# *De novo EHMT2* variants cause an autosomal dominant *EHMT2*-related Kleefstra syndrome via loss of G9a methyltransferase activity

**DOI:** 10.1101/2025.09.25.678439

**Authors:** Aleš Hnízda, Beatriz Martinez-Delgado, Diana Sanchez-Ponce, Javier Alonso, Jeanne Amiel, Tania Attie-Bitach, Ariadna Bada-Navarro, Beatriz Baladron, Eva Bermejo-Sanchez, Vítězslav Brinsa, Ivana Buková, Rosario Cazorla-Calleja, Sylvie Červenková, Shanshan Chow, Petr Dušek, Olha Fedosieieva, Marta Fernandez, Sourav Ghosh, Gema Gomez-Mariano, Andrea Gřegořová, Mark James Hamilton, Hana Hartmannová, Esther Hernandez-San Miguel, Marina Herrero-Matesanz, Kateřina Hodaňová, Alan Kádek, Jennifer Kerkhof, Tjitske Kleefstra, Didier Lacombe, Michael A. Levy, Estrella Lopez-Martin, Ruaud Lyse, Petr Man, Purificacion Marin-Reina, Ellen F. Macnamara, Haley McConkey, Petra Melenovská, Lidia M. Mielu, David Moore, Lenka Steiner Mrázová, Karolína Musilová, Kristýna Neffeová, Petr Nickl, David Pajuelo Reguera, Martina Pavlíková, Lea Pavlovičová, Manuel Posada, Jan Procházka, Kateryna Pysanenko, Sheila Ramos del Saz, Dmitrijs Rots, Jessica Rzasa, Radislav Sedláček, Viktor Stránecký, František Špoutil, Matthew L. Tedder, Louise Thompson, Cynthia J. Tifft, Frederic Tran Mau-Them, Helena Trešlová, Antonio Vitobello, Sarah Hilton, Christopher Campbell, Siddharth Banka, Daniel Jirák, Bekim Sadikovic, Jakub Sikora, Stanislav Kmoch, Maria J. Barrero, Lenka Nosková

**Affiliations:** Research Unit for Rare Diseases, Department of Pediatrics and Inherited Metabolic Disorders, First Faculty of Medicine and General University Hospital, Charles University in Prague, Czech Republic; Institute of Rare Diseases Research (IIER), Spanish National Institute of Health Carlos III (ISCIII), 28220 Madrid, Spain; Undiagnosed diseases program SpainUDP, Madrid, Spain; Centro de Investigación Biomedica en Red de Enfermedades Raras, (CIBERER), Madrid, Spain; Service de Médecine Génomique des Maladies Rares, Hôpital Necker-Enfants Malades, AP-HP, Paris, France; Department of Biochemistry, Faculty of Science, Charles University in Prague, Czech Republic; Czech Centre for Phenogenomics, Institute of Molecular Genetics, Czech Academy of Sciences, Prague, Czech Republic; Pediatric Neurology Department, Hospital Universitario Puerta de Hierro, Majadahonda, Madrid, Spain; Undiagnosed Diseases Program, National Institutes of Health, Bethesda, MD, USA; Department of Neurology and Center of Clinical Neuroscience, First Faculty of Medicine, Charles University and General University Hospital in Prague, Czech Republic; Verspeeten Clinical Genome Centre, London Health Sciences Centre, London, ON, Canada; Institute of Molecular and Clinical Pathology and Medical Genetics, University Hospital Ostrava, Czech Republic; West of Scotland Clinical Genetics Service, Queen Elizabeth University Hospital, Glasgow, UK; Institute of Microbiology, Czech Academy of Sciences, Prague, Czech Republic; Department of Clinical Genetics, Erasmus MC, Rotterdam, The Netherlands; Department of Human Genetics, Radboud University Medical Center, 6500 HB, Nijmegen, The Netherlands; Génétique Médicale, CHU Bordeaux, INSERM U1211 (MRGM), Université de Bordeaux, France; Département de génétique, APHP-Nord, Paris, France, Service de cytogénétique et génétique médicale, Hôpital de la Mère et de l’Enfant, CHU Limoges, Limoges France; University and Polytechnic Hospital La Fe, Valencia, Spain; Department of Pathology and Laboratory Medicine, Western University, London Ontario Canada; South East Scotland Genetic Service, Western General Hospital, Edinburgh, UK; Greenwood Genetic Center, Greenwood, SC, USA; Office of the Clinical Director and Medical Genetics Branch, National Human Genome Research Institute, National Institutes of Health, 10 Center Drive, Bethesda, MD, USA; Université Bourgogne Europe, CHU Dijon Bourgogne, Laboratoire de Génomique Médicale, FHU-TRANSLAD, Centre de recherche Translationnelle en Médecine moléculaire – Inserm UMR1231 équipe GAD, Dijon, France; Manchester Centre for Genomic Medicine, St Mary’s Hospital, Manchester University NHS Foundation Trust, Health Innovation Manchester, Manchester, UK; Division of Evolution, Infection and Genomics, School of Biological Sciences, Faculty of Biology, Medicine and Health, University of Manchester, Manchester, UK; Institute for Clinical and Experimental Medicine, 140 21 Prague 4, Czech Republic; Faculty of Health Studies, Technical University of Liberec, 461 17, Liberec, Czech Republic; Institute of Pathology, First Faculty of Medicine, Charles University and General University Hospital in Prague, Czech Republic

## Abstract

*EHMT1* and *EHMT2* genes encode human euchromatin histone lysine methyltransferase 1 and 2 (EHMT1 alias GLP; EHMT2 alias G9a) that form heteromeric GLP/G9a complexes with essential roles in epigenetic regulation of gene expression. While *EHMT1* haploinsufficiency has been established as the cause of Kleefstra syndrome 1, the pathogenesis of G9a dysfunction in human disease remains largely unknown. We identified seven *de novo EHMT2* variants in patients with clinical presentation, episignatures, histone modifications and transcriptomic profiles similar to those of Kleefstra syndrome 1. *In vitro* studies revealed that these variants encode for structurally stable G9a proteins that are catalytically incompetent due to aberrant interactions either with histone H3 tail or with S-adenosylmethionine. Heterozygous mice carrying a patient-derived variant exhibited growth retardation, facial/skull dysmorphia and aberrant behavior. Here we report pathogenic *EHMT2* variants that likely exert dominant-negative effect on GLP/G9a complexes and thus genocopy the *EHMT1* haploinsufficiency via a distinct molecular mechanism, defining an autosomal dominant *EHMT2*-related Kleefstra syndrome.

## Introduction

Histone methylation regulates chromatin compaction and dynamically orchestrates cellular transcription programs. Histone methylation status is maintained by the activity of histone methyltransferases and demethylases^1^. Their dysregulation has been implicated in many neurodevelopmental disorders including monogenic and multifactorial diseases (e.g. Alzheimer or Parkinson disease)^2^. So far, at least eleven distinct genetic syndromes caused by malfunction of histone methyltransferases have been described^3^.

*EHMT1* and *EHMT2* genes encode human euchromatin histone lysine methyltransferase 1 and 2 (EHMT1 alias GLP; EHMT2 alias G9a), respectively. GLP and G9a have identical domain architecture with high sequence similarity. Their main characteristic is the presence of a highly conserved SET domain that catalyzes the transfer of methyl group from S-adenosylmethionine to lysine 9 of histone H3 (H3K9)^4^. The SET domains assemble into functional dimers^5^ and govern the formation of heteromeric GLP/G9a complexes *in vivo*^6^. Despite the ability of these proteins to form functional homodimers, heterodimers are the preferred form *in vivo*, displaying the maximum enzymatic activity^6,7^. The proper formation and methylation activity of the GLP/G9a complexes is critical for epigenetic regulation of gene expression in many biological processes including oogenesis^8^, metabolic regulation^9^ or neuronal maturation during brain development^10^.

Haploinsufficiency of *EHMT1* was originally described in the *9q* subtelomeric deletion syndrome, a clinically recognizable neurodevelopmental disorder with congenital anomalies, specific dysmorphic features and other characteristic symptoms known as Kleefstra syndrome 1 (KLEFS1, OMIM# 610253)^11,12^. Over time, additional Kleefstra syndrome 1 patients have been identified with pathogenic frameshift, nonsense or missense *EHMT1* variants, which mostly lead to haploinsufficiency through the instability and decreased amount of functional GLP proteins^13,14^. Recently, a broader phenotypic spectrum of Kleefstra syndrome 1 associated with diverse biochemical properties of differently mutated GLP has been described^15^.

Homozygous loss of either GLP (*Ehmt1*) or G9a (*Ehmt2*) in mice results in embryonic lethality, underscoring the essential role of both proteins in early development^6^. Moreover, heterozygous *Ehmt1^+/-^* mice exhibit a spectrum of neurodevelopmental abnormalities, including facial/skull dysmorphia, behavioral deficits^16^, synaptic impairments and disrupted hippocampal neurogenesis^17^ recapitulating clinical features of Kleefstra syndrome 1^18^. Importantly, in contrast to severe effect observed for *Ehmt1^+/-^*, *Ehmt2^+/-^* mice present with only minimal abnormalities^19^.

While the molecular mechanisms of epigenetic dysregulation and pathogenesis involved in the clinical symptoms of Kleefstra syndrome 1 are still under investigation, the epigenetic hallmarks that characterize *EHMT1* haploinsufficiency are well described. Recent research revealed that patients with Kleefstra syndrome 1 show a specific DNA methylation signature (episignature) in peripheral blood^20^. This episignature can aid the diagnosis of patients with clinical symptoms suggestive of Kleefstra syndrome 1, but with no pathogenic *EHMT1* variants revealed by molecular genetic testing and/or with variants of uncertain clinical significance^15,20,21^. Additionally, abnormal epigenetic states have been described in Kleefstra syndrome 1 neurons derived from induced pluripotent stem cells (iPSCs) and in *Ehmt1^+/-^* mice, manifested by dysregulated H3K9me2 global levels^22,19^.

Despite the recognized functional cooperation and interdependency between GLP and G9a ^6,23–25^, clinical correlates of G9a dysfunction in humans have not yet been thoroughly described, although the potential involvement of *EHMT2* biallelic loss in a patient with neurodevelopmental delay has been recently suggested^21^.

Here we report clinical, structural, biochemical and molecular correlates of seven *de novo* heterozygous *EHMT2* variants that we identified through exome/genome sequencing. We show that all these *EHMT2* variants encode for structurally stable but catalytically inactive G9a proteins that seem to possess dominant negative effects on the function of GLP/G9a complexes. Patients harboring these variants showed a clinical phenotype consistent with Kleefstra syndrome 1. Patient-derived cell lines displayed alterations in episignatures, histone modifications and gene expression profiles closely resembling Kleefstra syndrome 1. Notably, the *EHMT2* patients showed a specific episignature in blood, enabling the diagnosis of an additional patient carrying a pathogenic *EHMT2* variant. Moreover, a knock-in mouse model harboring a patient-derived heterozygous deletion of four amino acids in the SET domain of G9a recapitulates multiple key phenotypic features observed in patients with *EHMT2* variants reported here.

## Results

### Multiple de novo variants in EHMT2 are associated with a neurodevelopmental phenotype and facial dysmorphism

Seven patients (P1-P7) carrying seven different *de novo* heterozygous variants in *EHMT2* gene have been identified through exome/genome sequencing in several national research or diagnostic programs for rare diseases. The investigators and health care professionals were gathered via GeneMatcher^26^, via Matchmaker Exchange using the RD-Connect Genome-Phenome Analysis Platform (GPAP)^27^ and personal communication. Five of the *EHMT2* variants encode for missense G9a mutations: NM_006709.5: c.3229G>T p.(Ala1077Ser), c.3229G>A p.(Ala1077Thr), c.3335A>G p.(Asn1112Ser), c.3472T>C p.(Phe1158Leu) and c.3485A>G p.(Lys1162Arg) and two variants encode for *in-frame* G9a deletions: NM_006709.5: c.3225_3236del p.Glu1076_Val1079del and c.3487_3492del p.(Ser1163_Lys1164del). None of these variants have been previously reported in the Genome Aggregation Database (gnomAD v4.1.0) and ClinVar database. The variants encode for evolutionary conserved residues and are predicted as pathogenic by *in silico* prediction tools (**Table 1**).

The patients’ cohort was composed of four females and three males aged 2 – 16 years. All patients presented with facial dysmorphism including midface hypoplasia, synophrys, hypertelorism, dysplastic ears, short philtrum and deeply set eyes, and cardiovascular anomalies including septal defects (ventricular or atrial), pulmonal valve stenosis and coarctation of the aorta. All but one patient presented with hypotonia, psychomotor and developmental delay, absence of speech and behavioral abnormalities. A single patient (P3) with the c.3335A>G p.(Asn1112Ser) variant had normal psychomotor, cognitive and speech abilities but exhibited paroxysmal hyperkinetic (choreoathetoid) movements. Individually, patients presented with various other abnormalities including pes planovalgus or equinovalgus, postaxial polydactyly or big halluxes, kidney cysts and other urogenital abnormalities. Detailed clinical description of patients is provided in **Table 1** and in **Supplementary Material**. Overall, the clinical presentation of the patients was very similar to those of Kleefstra syndrome 1 caused by *EHMT1* haploinsufficiency.

### Episignatures of patients with EHMT2 variants overlap with those of Kleefstra syndrome 1 yet reveal a specific hypomethylated episignature

Our initial discovery cohort consisted of patients P1 to P6. Analyses of patients’ blood DNA methylation profiles using the Episign assay^20,28–30^ revealed an episignature profile that matched the Kleefstra syndrome 1 episignature (Supplementary Figure 1). Unsupervised hierarchical clustering and multidimensional scaling (MDS) plots demonstrated that the patients with *EHMT2* variants clustered closely with the Kleefstra syndrome 1 cohort, with one sample (P3) showing complete overlap (Supplementary Figure 1A, B). Furthermore, a multi-class supervised classification model predicted the *EHMT2* patients as Kleefstra syndrome 1 with high confidence, while effectively distinguishing them from other established episignatures associated with genetic disorders in the EpiSign™ Knowledge Database (EKD) (Supplementary Figure 1C).

Next, we conducted a discovery analysis to assess the presence of an *EHMT2* specific episignature. For that, we compared the episignatures of the six patients carrying the EHMT2 variants and 54 matched controls, and identified 244 top differentially methylated probes. The mean methylation difference across the probes between patients and controls ranged from +45.52% to −35.77% (median = −18.60%), with 88.52% of the signature probes being hypomethylated (Supplementary Figure 2A). Interestingly, these probes were mapped to 155 genes that were enriched in the Gene Ontology term multicellular organism development at a p-value: 6.1E-7. Cluster analysis validated the feature selection, showing a clear separation between *EHMT2* patients and controls. The analysis also showed a clear separation from the testing Kleefstra syndrome 1 samples in both heatmap and multidimensional scaling (MDS) plots (**Figure 1A, B**). Furthermore, an unsupervised leave-one-out cross-validation on the discovery cases demonstrated that each hold-out test case consistently clustered with the remaining discovery cases in heatmaps and aligned closely with patient group centroids in MDS plots (Supplementary Figure 2B). This supports the reproducibility and robustness of the identified episignature. We then developed a Support Vector Machine (SVM)-based model to evaluate its diagnostic utility for *EHMT2*-related disorders. The model demonstrated high specificity, as nearly all EKD test samples yielded methylation variant pathogenicity (MVP) scores close to 0 (**Figure 1C**). It also showed sensitivity to an additional patient (P7) with a *de novo EHMT2* variant (EHMT2(NM_006709.5):c.3229G>A p.(Ala1077Thr) that was identified in the EKD database after the discovery cohort. This patient had an MVP score greater than 0.75, clustered with discovery cases in the MDS plot, and displayed similar profiles in the heatmap (**Figure 1A, B and C**). The clinical characteristics of this patient were consistent with those of the *EHMT2* patient cohort **(Table 1)**. Additionally, all testing Kleefstra samples had very low MVP scores (< 0.1), did not match the signature, and exhibited profiles more similar to controls. Overall, the episignature analysis suggest that *EHMT2*-related disorders are related to Kleefstra syndrome 1 but represent a distinct molecular entity with its own specific episignature.

**Figure 1.**
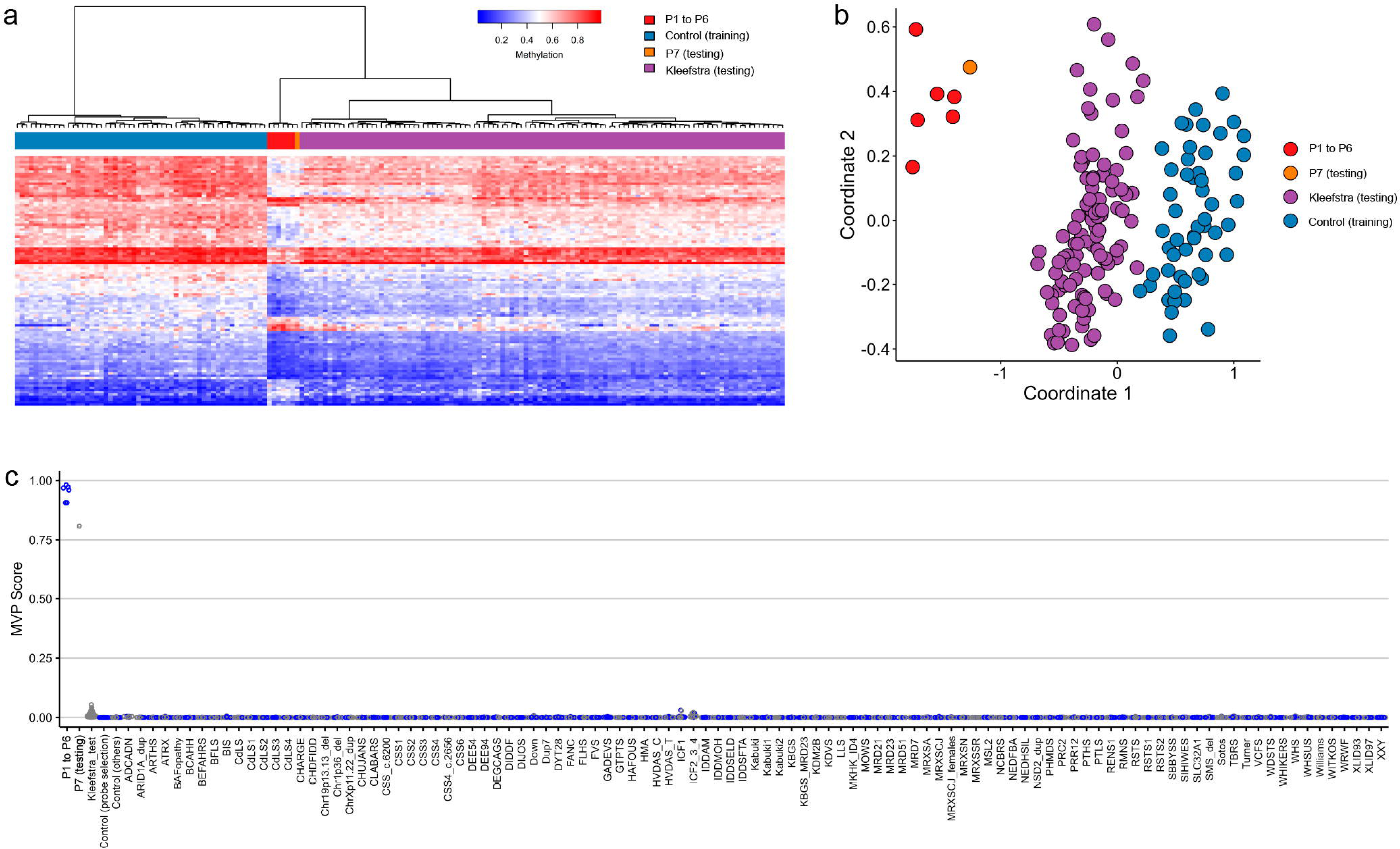
EpiSign (DNA methylation) analysis of peripheral blood from patients with *EHMT2* variants. Unsupervised clustering of the discovery cohort and assessment of *EHMT2* variants using the identified methylation signature, visualized via (A) heatmap and (B) multidimensional scaling (MDS) plot. Discovery samples from patients with *EHMT2* variants are shown in red, matched controls in blue, *EHMT2* test sample in orange, and Kleefstra (*EHMT1*) test samples in purple. (C) Methylation Variant Pathogenicity (MVP) scores from the SVM classifier for EpiSign™ Knowledge Database, *EHMT2*, and Kleefstra test samples, averaged over four-fold cross-validation (gray). Higher MVP scores indicate greater similarity to the discovery case methylation profile (blue).

### EHMT2 patients show histone modification and gene expression signatures that match those of Kleefstra syndrome 1

Next, we investigated whether the identified patients had a histone modification profile similar to the one reported for Kleefstra syndrome 1 and in accordance with the loss of G9a activity. We established fibroblast cultures from skin biopsy from patient P2, together with a healthy skin donor and a Kleefstra syndrome 1 patient (KSP) carrying a frameshift variant in *EHMT1* (NM_024757.5: c.1881delT p.(His629ThrfsTer12)). We further generated iPSCs from the established fibroblasts cultures and from peripheral blood mononuclear cells available from patient P1 that were characterized according to the established standards^31,32^ (Supplementary Figure 3 and 4). Quantification of histone modifications in fibroblasts and iPSCs lines by mass spectrometry revealed significantly (p-value<0.05) decreased levels of H3K9me1 or H3K9me2 in all patients’ cell lines compared to control cell lines (**Figure 2A**). While changes in histone modifications in iPSCs were rather subtle, more pronounced changes were observed in fibroblast cultures in which we detected significant downregulation of H3K9me1/2/3 and upregulation of unmodified and acetylated H3K9 in patients. The changes in histone modifications were consistent between Kleefstra syndrome 1 and P2 patient’s fibroblasts, showing coordinated upregulation of several histone marks involved in gene activation such as H3K56ac, H3K64ac and H3K79me3 (**Figure 2B**, Supplementary Figure 5A).

**Figure 2.**
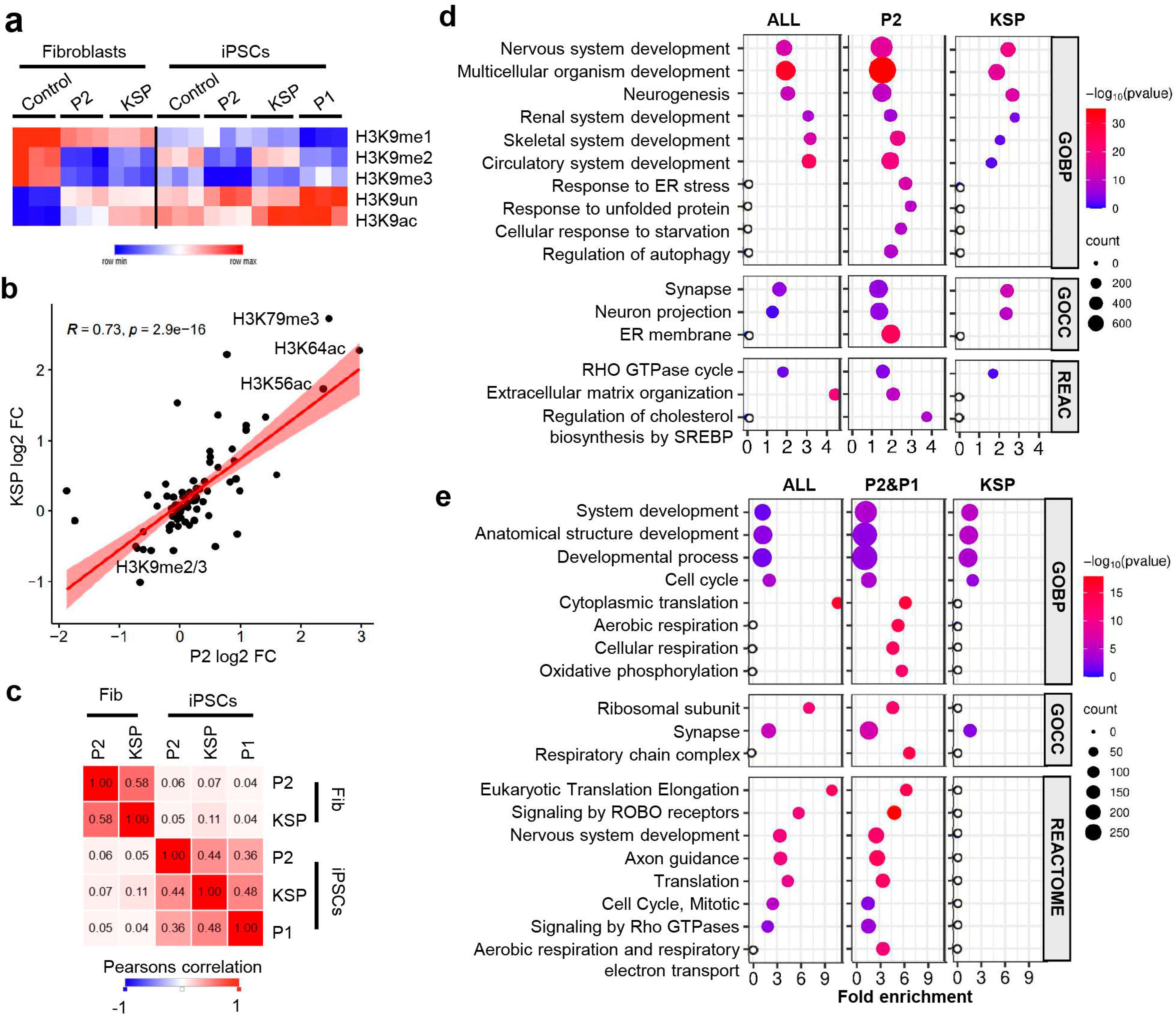
Epigenetic and transcriptomic analyses of cell lines derived from a healthy control, a Kleefstra Syndrome 1 patient (KSP) and patients P1 and P2 carrying *EHMT2* variants. (A) Relative abundance of the indicated histone H3K9 modifications quantified by mass spectrometry in fibroblasts cultures (Fib) and iPSCs in triplicates. (B) Correlation of changes in 92 histone modification states detected by mass spectrometry between KSP or P2 and control fibroblast. (C) Heatmap of Pearsons correlation coefficients of gene expression changes between the indicated cell lines. (D) DAVID enrichments of Gene Ontology Biological Processes (GOBP), Gene Ontology Cellular Compartments (GOCC) and Reactome (REAC) terms in genes commonly upregulated in KSP and P2 fibroblasts (ALL), upregulated in P2 fibroblasts only (P2) or KSP fibroblasts only (KSP). (E). DAVID enrichments of GOBP, GOCC and Reactome terms in genes commonly upregulated in P1, P2 and KSP iPSCs (ALL), upregulated in P2 and P1 (P1&P2) iPSCs only or KSP iPSCs only (KSP). Hollow circles indicate not significant at p-value>0.05.

In agreement with the altered histone modification patterns in the fibroblasts and iPSCs patient-derived cell lines, we observed significant changes in gene expression in patients compared to controls that correlated among patients (**Figure 2C**, Supplementary Figures 6A-C). Accordingly, there was a significant overlap between the patients’ cell lines in up- and downregulated genes (Supplementary Figure 6D-G). Upregulated genes, most likely G9a direct target genes^33–35^, were enriched in alternative lineages such as skeletal, muscle and neuronal systems development, and in cell cycle and Rho GTPase pathways. In addition, we identified genes exclusively upregulated in EHMT2 patients’ cell lines. Specifically, we observed a stress-related signature characterized by endoplasmic reticulum stress and unfolded protein response, accompanied by autophagy and increased cholesterol biosynthesis in fibroblasts and upregulated oxidative phosphorylation and aerobic respiration in iPSCs. None of these signatures were enriched among genes upregulated only in KSP cells or commonly upregulated in EHMT2 patients’ cell lines and KSP cell lines. (**Figure 2D and 2E**, Supplementary Figures 7 and 8). Overall, the histone modification and gene expression changes observed in patient-derived cells carrying EHMT2 variants largely overlap with those reported in Kleefstra syndrome 1, while also showing distinct differences, consistent with impaired H3K9 methyltransferase activity.

### EHMT2 variants have no effect on GLP and G9a transcript and protein levels

To explore the effects of the identified variants, we examined *EHMT1/2* transcripts and GLP/G9a protein amounts in the available patient-derived samples. RNA sequencing revealed very similar amounts of wild-type and mutant *EHMT2* transcripts in peripheral leukocytes of P1, P2 and P3 and similar transcript levels of EHMT1 and EHMT2 in peripheral leukocytes obtained from P1, P2 and P3, iPSCs from P1 and P2, and fibroblasts from P2 (Supplementary Figure 9A, B). The amounts of GLP and G9a protein were also very similar in patient-derived and control iPSCs as determined by western blot (Supplementary Figure 9C). Next, we immunoprecipitated G9a from iPSCs and analyzed co-precipitating proteins by western-blotting and mass spectrometry (Supplementary Figure 9D). We found that GLP, along with other known G9a interactors such as WIZ and ZNF462^36^ was efficiently co-immunoprecipitated in both control and patients’ cell lines. Overall, the identified *EHMT2* variants do not affect the formation of sufficient amounts of a structurally stable GLP-G9a complex.

### EHMT2 variants cluster at the G9a enzyme-substrate interface

Sequence alignment and structural modelling (**Figure 3**; Supplementary Figure 10) revealed that six of the predicted amino acid changes (Del 1076-9, A1077S, A1077T, F1158L, K1162R and Del 1163-4) affect the G9a binding site for histone H3 tail, whereas the N1112S substitution affects the G9a binding site for the methyl group donor (S-adenosylmethionine). Thus, all identified variants cluster at the G9a enzyme-substrate interface with potential effects on the histone H3 tail interaction and catalytic activity.

**Figure 3.**
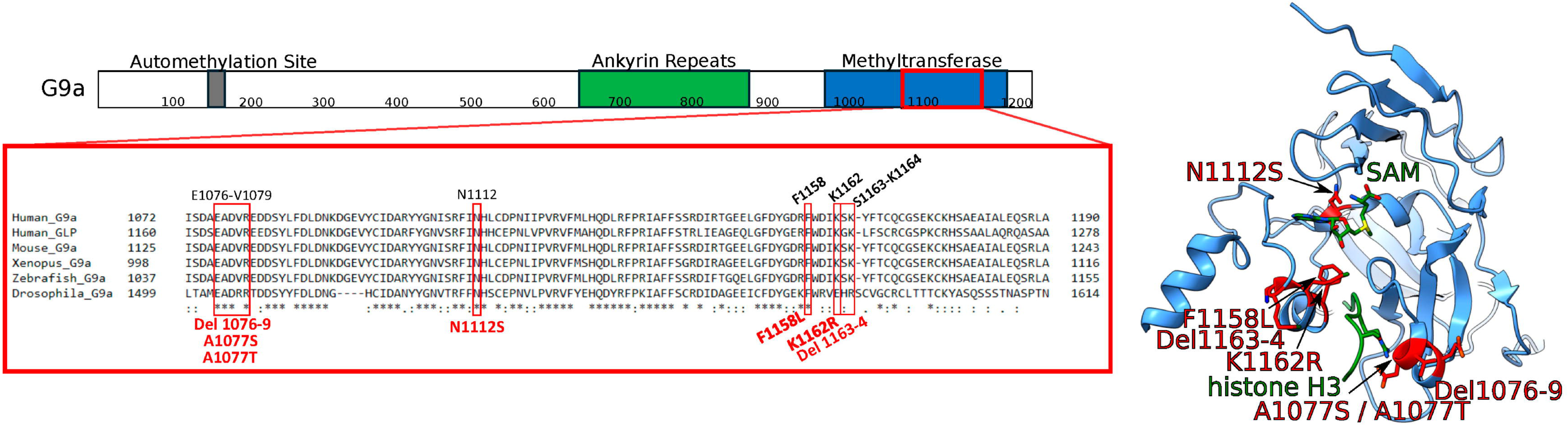
Sequence and structural positioning of mutations in the G9a protein. (A) Aminoacid sequence alignment shows highly conserved residues in the catalytic pocket of the methyltransferase domain. Positions of the mutations are highlighted by red squares. (B) Structural mapping demonstrates localization of the mutations in binding pockets for histone H3 tail and SAM in the G9a catalytic site. Mapping was performed using crystal structure of SET-domain of G9a in complex with SAM and histone H3 peptide (PDB ID 5JIN). The variant aminoacid residues are highlighted as red sticks; G9a substrates - histone H3 and SAM - are depicted in green color.

### All EHMT2 variants compromise the catalytic activity of G9a through defective enzyme-substrate interactions

Since structural modelling showed that all mutations cluster at the G9a enzyme-substrate interface, we cloned, expressed and purified recombinant methyltransferase domains and examined their catalytic activities. Specifically, we determined the extent of methylation of K9 in an N-terminal histone H3 peptide (amino acid residues 1-20) and nucleosome. Initially, using semi-quantitative analysis by MALDI-TOF mass spectrometry we observed that mono- and di-methylation of histone H3 peptide was decreased with A1077S, A1077T, K1162R and Del 1163-4 mutations and absent in Del 1076-9, N1112S and F1158L mutations (Supplementary Figure 11). Subsequently, we used a bioluminescence-based assay analysis to quantify the initial velocity rates of lysine methylation on histone H3 peptide or nucleosome. We observed that six of the mutations, Del 1076-9, A1077T, N1112S, F1158L and K1162R, Del 1163-4, decreased the methylation activity towards histone H3 peptide to less than 10% of the wild-type enzyme (**Figure 4A**). A higher (∼ 20%) residual activity was observed for the A1077S mutation. Notably, the methylation of mononucleosome was reduced to less than 5% of the wild-type enzyme activity with all studied mutations, including the A1077S. These data demonstrate that all studied mutations reduce the catalytic competence of G9a towards the histone H3 tail.

**Figure 4.**
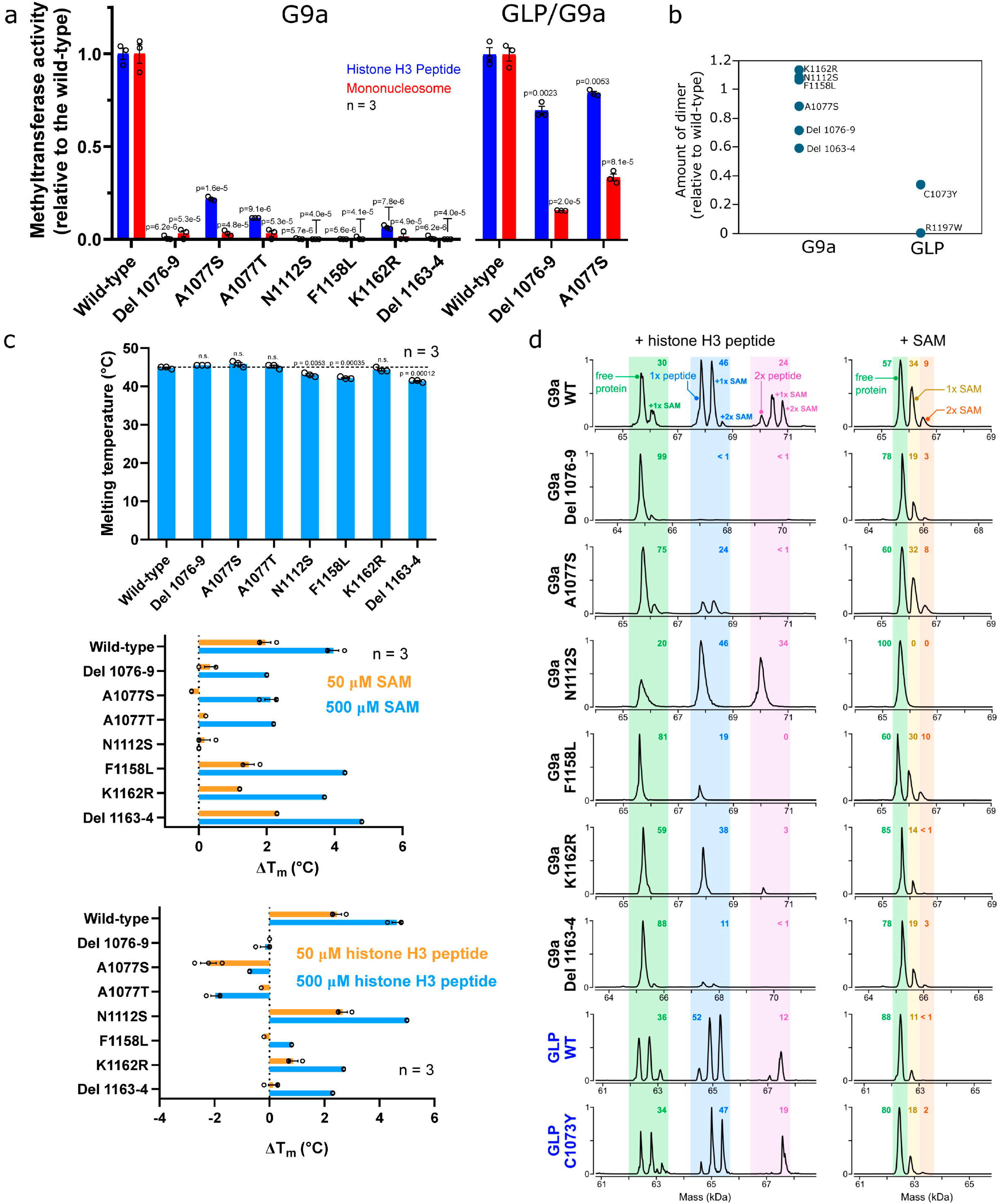
Functional characteristics of G9a variants assessed with recombinant proteins. (A) Methylation activity of the G9a mutants with histone H3 peptide (blue) and mononucleosome (red) as substrates studied by quantitative bioluminescence-based assay (Mtase-Glo by Promega). The activity was assessed for G9a homodimers (left) and GLP-G9a heterodimers (right). The values represent the means of relative activity to the wild-type enzyme with standard errors from three measurements. The p-values indicate variants with significant differences from the wild-type. whereas “n.s.“means changes were not significant, as determined by a two-tailed Student’s t-test (p<0.01). The symbol “#” indicates “activity not detected”. (B) Quantitative assessment for dimer formation of G9a and GLP mutants using native mass spectrometry (nESI-MS). Points represent relative amounts of dimers related to the wild-type proteins. Dimerization is severely affected in GLP but not in G9a mutants. (C) Thermostability of the G9a mutants in their free and substrate-bound states studied by differential scanning fluorimetry in at least in triplicates. The upper panel shows the melting temperatures of free G9a proteins are represented as mean values with standard errors. Dashed line indicates melting temperature of the wild-type protein. The p-values indicate variants with significantly different changes from the wild-type. whereas “n.s.“ means changes were not significant, as determined by a two-tailed Student’s t-test (p<0.01). The two lower panels show the stabilization of G9a proteins upon the addition of individual substrates (SAM or histone H3 peptide), as shown by differences in melting temperature in comparison to the free protein. The values represent the means with standard errors from three measurements. (D) Binding affinity of the G9a variants towards individual substrates assessed by native mass spectrometry. The protein and substrate concentrations were maintained at 5 µM. Representative deconvoluted nMS spectra revealed specific mass shifts that corresponded G9a complexes with SAM (+398 Da) or histone H3 peptide (+2100 Da) bound as shown by highlighted areas with relative abundance.

Next, we examined the impact of two G9a variants with compromised catalytic activity (Del 1076-9 and A1077S) in the context of the GLP-G9a heterodimer. In the GLP-G9a heterodimer, ankyrin repeat domains are required for efficient methylation with nucleosomal substrates^7^. Therefore, we co-expressed GLP and G9a constructs comprising the ankyrin repeat region and the methyltransferase domain and purified the formed heterodimeric complexes. A bioluminescence-based methylation assay revealed only a modest reduction in activity of the G9a mutant heterodimers with histone H3 peptide (≈ 70 % of wild-type), whereas activity with nucleosomes was markedly diminished (< 35 % of wild-type) (Figure 5A). These data indicate that the G9a variants substantially impair nucleosomal methylation activity of the GLP-G9a heterodimer and that heterodimer-associated GLP cannot compensate for catalytically deficient G9a mutants.

**Figure 5.**
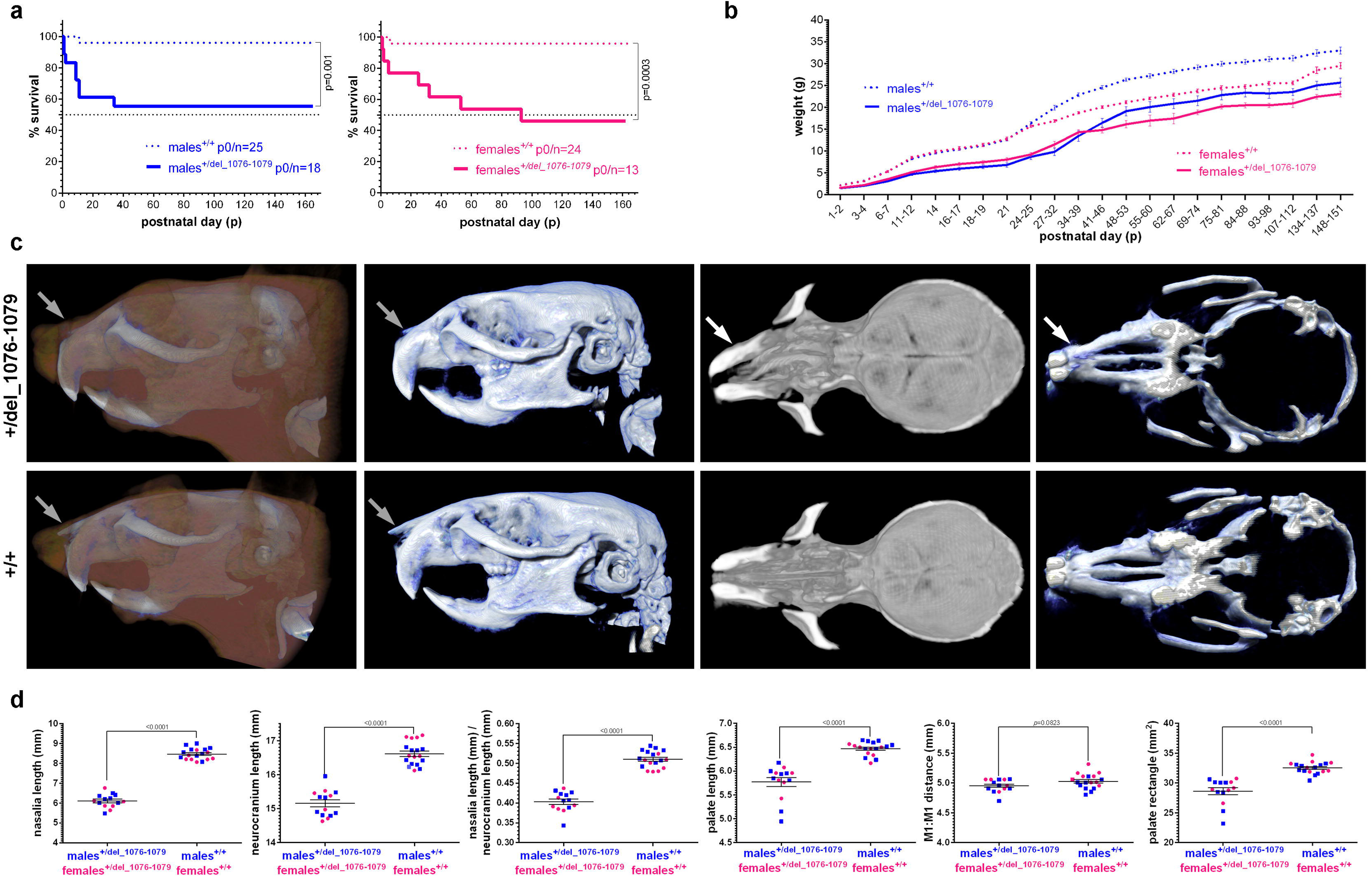
Key phenotypic features identified in *Ehmt2^+/del_1076–1079^* mice. (A) Survival data (5 months) comparing *Ehmt2^+/del_1076–1079^ and Ehmt2^+/+^* mice are shown for each sex. p0/n corresponds to the starting number of animals included into the study. Black dotted line highlights 50 % survival rate. Statistical significance was calculated using Log-rank (Mantel-Cox) test. (B) Weight curves (5 months) for all animals of both genotypes/sexes included in the survival study. Weights were significantly different (p<0.05 by a two-tailed Student’s t-test) between the *Ehmt2^+/del_1076–1079^* and *Ehmt2^+/+^*mice of each sex at all postnatal time points. Refer to Supplementary Figure 16 for detailed weight comparisons at p1-2 and p21 and weight curves of animals surviving the entire 5 months period. (C) MicroCT analyses in *Ehmt2^+/del_1076–1079^ and Ehmt2^+/+^* mice. Lateral views of the skulls with (in brown) and without soft facial tissues, horizontal views of the nasal cavity and neurocranium, and 3D reconstruction of the palate (bottom view). Nasalia (grey arrows) are shorter in *Ehmt2^+/del_1076–1079^* mice. White arrows highlight the “bent” nose phenotype in *Ehmt2^+/del_1076–1079^* animals. For the proportion and range of the shorter nasalia and the “bent nose” abnormalities see Supplementary Figures 18-19. (D) Quantitative microCT craniometric data comparisons in *Ehmt2^+/del_1076–1079^ and Ehmt2^+/+^* mice. *Ehmt2^+/del_1076–1079^* have significantly shorter nasalia and disproportionately shorter neurocranium, which results in a significantly lower ratio of these two values in comparison to *Ehmt2^+/+^* mice. Palate length is shorter in *Ehmt2^+/del_1076–1079^* while the palate width (molar1: molar1 – M1:M1 distance) remains comparable (p=0,083) to *Ehmt2^+/+^* mice. As a result, the palate (calculated as a square size of the latter two values) is significantly smaller in *Ehmt2^+/del_1076–1079^* animals. Statistical significance was calculated using two-tailed t-test. Values are color/shape-coded based on sex. See Supplementary Figure 17 for technical and acquisition details of the presented craniometric measurements. B and D show mean values with standard errors. Evaluators were blinded to the genotype of individual mice.

To identify the molecular mechanisms of this reduced catalytic competence, we assessed the structural stability and binding affinity of the recombinant G9a mutants towards their substrates. First, we used native mass spectrometry (nMS), which measures the mass of intact non-covalent protein complexes^37^, to quantify the relative amounts of properly folded dimeric forms of the corresponding mutated proteins. Native MS revealed that all G9a mutants predominantly assemble into compactly folded dimers (**Figure 4B** and Supplementary Figure 12A) and that GLP-G9a heterodimer formation is favored upon co-expression in all protein variants (Supplementary Figure 12B). Using differential scanning fluorimetry (DSF) performed under temperature gradient, we determined that the melting temperatures of free homodimeric G9a proteins were slightly reduced by N1112S, F1158L, and Del1163-4 mutations and not affected by Del 1076-9, A1077S, A1077T and K1162R mutations (**Figure 4C**). These data showed that the variants do not have major destabilizing effects on the overall G9a structure and its dimerization.

Next, we determined the binding affinities of individual G9a mutations for histone H3 peptide and SAM using nMS. Binding affinities were estimated from observed substrate binding visible as 398 Da (for SAM) or 2100 Da (for histone H3 peptide) mass increases of corresponding non-covalent protein-ligand complexes. This analysis revealed that mutants Del 1076-9, A1077S, F1158L, K1162R and Del 1163-4 had reduced binding to histone H3 peptide but unaffected binding to SAM (**Figure 4D**). Opposite results were observed for the N1112S mutation that exhibited normal binding to histone H3 peptide but failed to interact with SAM. Substrate binding behaviour of individual G9a mutations were further corroborated by binding affinity estimated from substrate concentration-dependent increased thermostability of corresponding protein-ligand complexes using DSF. This analysis further confirmed the decreased binding capacity of Del 1076-9, A1077S, F1158L, K1162R and Del 1163-4 mutations to histone H3 peptide and revealed that the Del 1076-9 and A1077S mutations had partially reduced binding capacities to SAM (**Figure 4C**). Consistently with our observations above, opposite results were recorded for the N1112S mutation that exhibited normal binding to histone H3 peptide but failed to interact with SAM.

Together, nMS and DSF revealed that pathogenic G9a mutations disturb the catalytic competence of the enzyme through defective interaction with one or both of enzyme substrates, while the structural integrity and stability of the protein is not affected.

Using the established protocol, we also examined the heterozygous G9a variant p.T961I previously reported as a variant of unknown significance in a patient with an autistic disorder and no phenotypic overlap with Kleefstra syndrome 1^38^. This variant, located outside the methyltransferase site, exhibited normal methyltransferase activity and protein stability (Supplementary Figure 13). Thus, its biochemical profile differs from that of the pathogenic variants identified in this study. Given these results, we classify this variant as likely benign.

### Pathogenic EHMT1 SET domain variants affect protein stability and dimer formation

To compare the *EHMT2* variants with previously identified pathogenic missense *EHMT1* variants, we assessed the effects of two previously described SET-domain GLP pathogenic mutants: C1073Y and R1197W^13^.

Our biochemical examinations revealed that both mutations render the GLP SET-domain considerably unstable when expressed in *E.coli*, with most of the mutant proteins accumulating in insoluble fractions, which either entirely abolished (R1197W) or decreased (C1073Y) purification yields (Supplementary Figure 14A, B). With reduced but sufficient amounts of purified C1073Y mutant, we were able to determine its decreased thermal stability and impaired dimerization (**Figure 4B, D**). Despite this, the C1073Y mutant showed binding affinities to histone H3 peptide and SAM close to those of wild-type GLP (**Figure 4D**) and had a relatively high (∼ 60%) methylation activity with histone H3 peptide but almost no (< 1%) activity with mononucleosome compared to the wild-type protein (Supplementary Figure 14C).

To further examine the aberrant catalysis of mutated methyltransferases, we compared the kinetic parameters of G9a (A1077S) and GLP (C1073Y) variants retaining relatively high residual activity toward histone H3 peptide. This analysis revealed that the C1073Y GLP mutation did not significantly affect enzyme turnover number (k_cat_), while the A1077S G9a mutation significantly decreased it. Specifically, k_cat_ values for A1077S G9a were 2.2-fold and 4.5-fold lower toward histone H3 peptide and SAM, respectively, compared to the wild-type enzyme. Therefore, our biochemical and enzymatic analyses confirm that pathogenic G9a variants are catalytically incompetent *per se,* while the tested pathogenic missense GLP mutations lose their activities due to structural instability.

### Ehmt2^+/del_1076–1079^ mice present with postnatal developmental abnormalities, craniofacial dysmorphia and gait alterations

To further validate the functional relevance of the *EHMT2* variants at the organismal level, we generated a mouse *knock-in* model (*Ehmt2^+/del_1076–1079^*) lacking four amino acids in the SET domain of G9a that corresponds to the deletion identified in patient P1 (Supplementary Figure 15).

*Ehmt2^+/del_1076–1079^* and littermate *Ehmt2^+/+^* control mice used in the studies were generated by *in-vitro* fertilization. None (out of 13) *Ehmt2^+/del_1076–1079^* females gave any progeny when mated with fertile *Ehmt2^+/+^* males. Two (out of 18) *Ehmt2^+/del_1076–1079^* males reproduced once with fertile *Ehmt2^+/+^* females.

Approximately half of the *Ehmt2^+/del_1076–1079^* animals of both sexes did not survive over postnatal day (p) 30-40 (**Figure 5A**). We were able perform urgent autopsies and/or retrieve carcasses for tissue analyses from 7 animals. Gross anatomical and histological analyses did not show any cardiac developmental defects. All surviving mice were terminated no later than p168. In comparison to sex-matched *Ehmt2^+/+^* mice, *Ehmt2^+/del_1076–1079^* presented with significant (and long-term lasting) weight-gain deficits detectable as early as p1-2 (**Figure 5B**, Supplementary Figure 16).

At 3-4 months of age, mice were subjected to facial/skull morphology assessment by microCT, SHIRPA scoring and a battery of behavioral tests (open field, gait analysis, rotarod and grip strength), and echocardiography. MicroCT and craniometric analyses identified a systematically occurring dysmorphia comprised of saddle and “bent” nose with shorter nasalia, nasal cavity disorganization, shorter and smaller palate and smaller (shorter) neurocranium in *Ehmt2^+/del_1076–1079^* mice (**Figure 5C** and **5D**, Supplementary Figures 17-19). Importantly, sphenoid bone formation did not seem to be impacted in *Ehmt2^+/del_1076–1079^* in comparison to *Ehmt2^+/+^* animals (Supplementary Figure 20).

The overall SHIRPA scores were comparable between the Ehmt2^+/del_*1076–1079*^ and Ehmt2^+/+^ animals of both sexes, but the locomotor activity was significantly lower in *Ehmt2^+/del_1076–1079^* animals (Supplementary Figure 21). This observation was consistent with results of the open-field test, in which *Ehmt2^+/del_1076–1079^* mice travelled significantly shorter distances at lower speeds and rested more than their *Ehmt2^+/+^* littermates. Interestingly, no difference in center zone preference/avoidance suggestive of anxious behavior was observed between the two alternative genotypes/sexes (Supplementary Figure 22). Gait analysis of *Ehmt2^+/del_1076–1079^* mice reveals a clear phenotype of compromised motor control and stability during the critical transitions of the locomotive cycle, evidenced by prolonged braking times and reduced maximal rates of paw contact area change during the braking phase. These deficits are compounded by postural abnormalities, including increased limbs midline distance and decreased paw overlap - that signal a fundamental breakdown in both intra- and inter-limb coordination (Supplementary Figure 23).

Similarly to the open field test, *Ehmt2^+/del_1076–1079^* mice had significantly shorter overall times spent, and distances travelled on the rotating rod (rotarod test). While not significantly different, *Ehmt2^+/+^* mice had higher average difference in the times-to-fall between the first and last (6^th^) trial on the beam (Supplementary Figure 24). Last, grip strength values were comparable between the two genotypes (Supplementary Figure 25). Transthoracic echocardiography did not identify any major structural or functional cardiac abnormalities in *Ehmt2^+/del_1076–1079^* mice, as left ventricular morphology and systolic function were largely preserved, and the observed reduction in cardiac output was secondary to a lower heart rate rather than intrinsic ventricular dysfunction (Supplementary Figures 26-29).

Together, the mouse model verified *in vivo* the pathogenicity of the identified heterozygous *EHMT2* variant.

## Discussion

Here, we report for the first time clinical and molecular correlates of *de novo* pathogenic heterozygous loss-of-function variants in the *EHMT2* gene in patients with clinical characteristics overlapping Kleefstra syndrome 1. Specifically, the probands reported here had recognizable facial dysmorphism (hypertelorism, synophrys, flat face, everted lips, protruding jaw) (See Figure 1) and cardiac anomalies (septal defects, pulmonal valve stenosis, aortal coarctation). Moreover, the probands exhibited additional phenotypic symptoms typical for Kleefstra syndrome 1 including psychomotor developmental delay, intellectual disability, absent speech, structural brain anomalies, hypotonia, dental anomalies, behavioral and sleep difficulties. These symptoms were observed in all patients except P3, whose clinically milder phenotype was reduced to dysmorphic features and paroxysmal hyperkinetic movement disorder. The clinical similarity of these patients to those with Kleefstra syndrome 1 caused *by EHMT1* haploinsufficiency was accompanied by their similar episignature profiles in blood indicating an overall similarity and overlap in the methylation profiles in these two syndromes. Similar clustering of episignatures has been reported for epigenetic disorders with malfunction of the BAF complex due to various mutations in its individual subunits^39^. Despite the overlap with the Kleefstra syndrome 1 episignatures, we identified a distinct set of differentially methylated probes that form a robust, condition-specific episignature for EHMT2 that can be used to reliably identify additional patients with pathogenic *EHMT2* variants as demonstrated for patient P7. This probe set provides a highly accurate, sensitive, and specific classifier that clearly differentiates EHMT2 patients from EHMT1 patients and patients with other disorders with known episignatures. Most probes in this episignature were hypomethylated compared to healthy controls and mapped to genes involved in development. These findings agree with the reported role of G9a in maintaining DNA methylation at germline genes during mouse embryogenesis^34^. Overall, our observations indicate that mutations in *EHMT1* and *EHMT2* affect the same epigenetic module, albeit with specific functional differences. These findings further highlight the value and clinical relevance of episignature analysis in assessing variants of uncertain significance.

Consistent with the aberrant episignature profiles, we observed alteration in histone modifications and gene expression profiles in patient-derived cells. In line with our findings in fibroblasts, coordinated downregulation of H3K9me2 and upregulation of H3K9ac levels have been described in mouse embryonic stem cells knock-out for G9a^24^. Moreover, similar global changes in histone modifications have been observed by mass spectrometry in HEK293 cells with *EHMT1* and *EHMT2* knock-down, including decreased H3K9 methylation and increased levels of methylated H3K79^40^. Patient-derived iPSCs exhibited only minor changes in histone modifications, limited to a subtle yet significant decrease in H3K9 methylation, which could be explained by low levels of this mark in iPSCs compared to fibroblasts (Supplementary Figure 5B). These differences are consistent with previous reports describing a significant reduction of the repressive marks H3K9me2/3 and H3K27me2/3 during the reprogramming of MEFs to pluripotency^41^ and their re-establishment during differentiation to silence lineage-inappropriate gene expression^42^. Accordingly, changes in histone modifications were accompanied by upregulation of genes involved in alternative lineages such as nervous, muscle and skeletal system development, as well as genes involved in cell cycle and Rho signaling pathways. This upregulation is concordant with previously reported gene expression changes resulting from G9a dysfunction^43–47^. Consistent with the presence of an *EHMT2*-specific episignature, we identified gene sets and functional enrichments that were selectively upregulated in EHMT2 patients’ cell lines. Whether these unique signatures reflect *EHMT2*-specific molecular functions or a stronger perturbation of the common *EHMT1/EHMT2* epigenetic program remains to be determined.

Kleefstra syndrome 1 phenotype was recapitulated in the mouse model (*Ehmt2^+/del_1076–1079^*) harboring a heterozygous *Ehmt2* variant equivalent to Del 1076-9 (identified in patient 1). *Ehmt2^+/del_1076–1079^* mice exhibited significantly impaired postnatal survival, growth and weight-gain retardation, decreased locomotor activity, gait abnormalities and were mostly infertile. Cranial/facial abnormalities identified in these mice resembled the dysmorphic features observed in the patients and in specific details (e.g. “bent” nose) overlapped with pathologies earlier detected in the *Ehmt1^+/-^* mouse model^18^.

All the identified *EHMT2* variants disturbed the catalytic site of the SET domain and resulted in a loss of methyltransferase activity. Interestingly, all patients presenting with classical symptoms of Kleefstra syndrome 1 carried *EHMT2* mutations that affected the histone H3 binding site. Conversely, patient P3 with the milder phenotype was heterozygous for the variant N1112S that affects the SAM binding pocket. In addition, the episignature of this patient clustered more closely with Kleefstra syndrome 1 than with the other *EHMT2* patients. These observations suggest that the severity of the phenotype depends on whether the defective enzyme-substrate interactions affect the binding affinity towards histone H3 tail or SAM.

Given the functional interdependency between GLP and G9a, it is remarkable that, twenty years after the initial description of Kleefstra syndrome 1, no individuals with *EHMT2* alterations have been reported. Kleefstra syndrome 1 is commonly caused by microdeletions or point mutations in *EHMT1* leading to decreased GLP protein levels^22,48^. In agreement with previous reports, we describe that GLP pathogenic mutants C1073Y and R1197W are structurally unstable and/or cause impaired dimerization^13,15^, which is compatible with the mechanism of haploinsufficiency. In contrast, the G9a mutants reported here formed structurally stable dimers that were catalytically inactive. Moreover, *EHMT2* variants did not affect the levels of transcript and protein of both *EHMT1* and *EHMT2* in blood and patient-derived cell-lines, nor did they affect the ability to form stable GLP-G9a complex in the patients’ cells. These findings suggest that the pathogenic mechanism of *EHMT2* variants differs from that of *EHMT1*, which is consistent with previously published data showing complementarity of GLP and G9a enzymes but non-overlapping functions^49^.

Population genomics also point out a different mechanism of pathogenesis for both genes. The Genome Aggregation Database (gnomAD 4.1.0) shows marked differences in the probability of loss-of-function intolerance scores for *EHMT1* (LOEUF = 0.16) and *EHMT2* (LOEUF = 0.49) suggesting a high level of intolerance to loss of *EHMT1* function, in contrast to *EHMT2*. Similarly, the Database of Genomic Variants (DGV) shows marked differences in occurrence of genic deletions in the healthy population that are sporadic for *EHMT1*, but frequent for *EHMT2* (Supplementary Figure 30A). Consistently, the Database of Chromosomal Imbalance and Phenotype in Humans Using Ensembl Resources (DECIPHER), a repository for variants identified in patients with rare diseases, shows a high frequency of *EHMT1* deletions, but not that of *EHMT2* (Supplementary Figure 30B). Additionally, gnomAD v4.1.0 documented numerous patients of variants (6 variants with 27 allele counts) positioned in methyltransferase active site that were specifically identified in *EHMT1* and not in *EHMT2*. Of particular note is the *EHMT1* variant N1200S, an equivalent to N1112S in EHMT2 patient described here, which was reported in 14 individuals in gnomAD v4.1.0. In summary, the population genomic data support a notion that *EHMT2* haploinsufficiency is not sufficient to cause disease, in contrast to *EHMT1*.

Additional evidence supporting a differential mechanism of pathogenesis for *EHMT1* and *EHMT2* is provided by the mouse model. *Ehmt1*⁺/⁻ mice have been reported to exhibit significant changes in histone methylation and transcriptome profiles, leading to severe cognitive impairments^17^. In contrast, *Ehmt2*⁺/⁻ mice display only mild alterations in epigenetic and transcriptomic profiles, resulting in minor phenotypic effects, according to two independent studies^19,24^. However, our mouse model (*Ehmt2^+/del_1076–1079^*) displays severe phenotype abnormalities. This suggests that the reported SET-domain variants are more pathogenic than the heterozygous loss of *EHMT2*. Overall, our experimental results, population genetics data and phenotypes of the mouse model suggest that the described *EHMT2* pathogenic variants probably act via dominant-negative effects (**Figure 6**).

**Figure 6.**
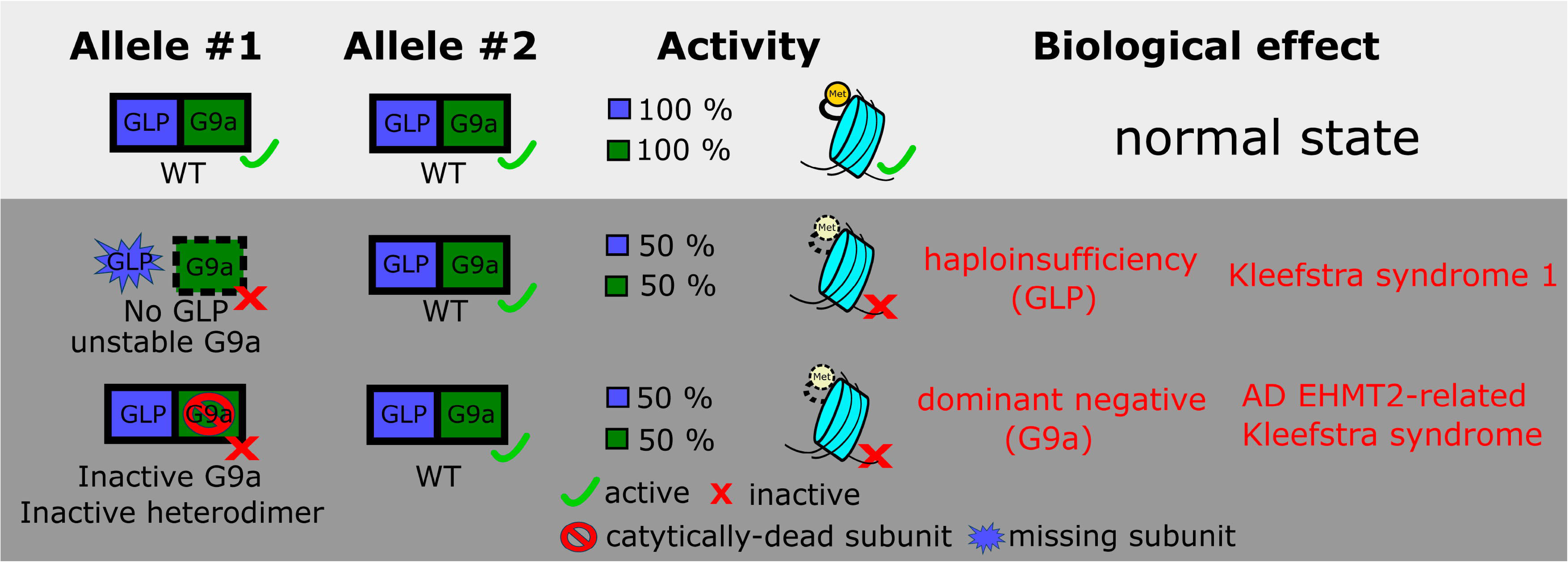
Scheme showing various scenarios for partially dysfunctional GLP/G9a heterodimer, that lead to physiological (upper part) or aberrant (lower part) H3K9 methylation.

Mechanistically, differences in the pathogenic mechanisms of *EHMT1* and *EHMT2* can be explained by the different biochemical properties and cellular functions attributed to both enzymes. While mouse embryonic stem cells deficient in GLP show significantly reduced levels of G9a protein, G9a-deficient cells exhibit normal GLP levels^6^. These findings indicate that G9a protein stability is highly dependent on the presence of GLP, while GLP stability is independent of G9a. Therefore, G9a’s biological function seems to be exclusively exercised through the G9a/GLP heterodimer, whereas GLP might be capable of forming both homo- and heterodimeric complexes.

In addition, each subunit of the GLP/G9a complex plays a distinctive role in governing the physiological levels of H3K9 methylation. Specifically, GLP contributes to the propagation of H3K9 methylation via recognition of H3K9me1/2 modified nucleosomes through its ankyrin-repeats domain (denoted as reader function)^49^, whereas G9a executes the methylation reaction *per se* by its enzymatically active SET domain^25^. Consistently, a recent report evaluating 209 individuals with rare variants in *EHMT1* found that missense variants in the GLP ankyrin repeat domain resulting in structurally stable GLP protein yet impairing its reader function were associated with Kleefstra syndrome 1^15^. In addition, variants that disrupted the GLP methyltransferase activity without affecting its structural stability could not be correlated with Kleefstra episignatures and phenotype^15^. These data indicate that both normal structural stability and reader function of GLP together with G9a writer activity are necessary conditions for the proper function of the GLP/G9a complexes. It is therefore plausible that pathogenic variants in *EHMT1*, but not in *EHMT2*, lead to haploinsufficiency caused by the reduced number of functional dimeric units of GLP/G9a. This reduction might be compensated by the formation of GLP homodimers in the case of limited G9a availability (e.g. in *Ehmt2^+/-^* mice), which seems unlikely to apply in the reverse scenario. In contrast, the pathogenic variants in *EHMT2* identified in this study give rise to stable GLP/G9a complexes with severely impaired nucleosomal methylation activity. These inactive mutant complexes might exert dominant-negative effects by binding to chromatin and blocking the access of the other functional complexes, resulting in disrupted epigenetic regulation.

In summary, we demonstrate that *EHMT2* missense variants located in the SET domain, which impair its catalytic function, lead to clinical symptoms that genocopy Kleefstra syndrome 1. This is supported by Kleefstra syndrome 1-like epigenetic changes, aberrant biochemical behavior of the mutant proteins and recapitulation of Kleefstra syndrome 1 phenotype in the mouse model. We also provide additional evidence that *EHMT2* and *EHMT1* variants act through different pathogenetic mechanisms in the context of the GLP/G9a complex. Therefore, the characterized patients with pathogenic *EHMT2* variants establish an autosomal dominant *EHMT2*-related Kleefstra syndrome. We believe that our findings will further facilitate the identification of additional individuals affected by pathogenic *EHMT2* variants and promote further research of the physiological and pathogenetic aspects of GLP/G9a biology.

### Methods (in supplementary file)

## Acknowledgments and Funding

This work was supported from the project MULTIOMICS_CZ (Programme Johannes Amos Comenius, Ministry of Education, Youth and Sports of the Czech Republic, ID Project CZ.02.01.01/00/23_020/0008540) – Co-funded by the European Union. L.N., A.H., and J.S. were supported by the project National Institute for Neurological Research (Programme EXCELES, ID Project No. LX22NPO5107) - Funded by the European Union – Next Generation EU and by institutional program UNCE/MED/007 of Charles University in Prague. A.H. was funded from the European Union’s Horizon 2020 research and innovation programme under the Marie Skłodowska-Curie grant agreement No 101003406 (HIPPOSTRUCT) and by the grant 26-21355S from the Czech Science Foundation. L.N., J.S. and S.K. were supported by grants NU23-07-00281 and NW24-04-00067 from the Ministry of Health of the Czech Republic. L.N., V.S. and S.K thank the National Center for Medical Genomics (LM2023067) for WES analyses. A.K. acknowledges funding by the European Union’s ERA fellowship (2D-TOPMASS, grant 101090276). The work was further supported by the grants from the Spanish Ministry of Science and Innovation PID2021-128087OB-I00 by MCIN /AEI /10.13039/501100011033 / FEDER, UE to M.J.B, Fundación Inocente grant number FII2024-125 and grant AESI PT20CIII/00009 (ISCIII Platform of Biobanks and Biomodels PT-20/23) from the Spanish National Institute of Health Carlos III. Funding was also partially provided by the Genome Canada and the Ontario Genomics Institute Genomics Applications Partnership Program Grant (OGI-188). Several co-authors are also members of the European Reference Network ITHACA. We further acknowledge the Solve-RD project, which has received funding from the European Union’s Horizon 2020 research and innovation programme under grant agreement No. 779257. We further acknowledge CMS-Biocev (“Biophysical techniques, Crystallization, Diffraction, Structural mass spectrometry”) of CIISB, Instruct-CZ Centre, supported by MEYS CR (LM2023042) and European Regional Development Fund-Project „UP CIISB“ (No. CZ.02.1.01/0.0/0.0/18_046/0015974). This study used tools provided through the RD-Connect GPAP, which received funding originally from the European Union Seventh Framework Programme (FP7/2007-2013) under grant agreement No. 305444. We also thank the Bioinformatics and the Advanced Optical Microscopy Units at ISCIII, the patients’ association Kleefstra España, the BioNER and the SpainUDP consortium. This study has also been delivered through the National Institute for Health and Care Research (NIHR) Manchester Biomedical Research Centre (NIHR203308). S.B. acknowledges the support of the MRC Epigenomics of Rare Diseases (EpiGenRare) Node (MR/Y008170/1). D.J. acknowledges the financial support from the Ministry of Health of the Czech Republic (MH CZ-DRO, Institute for Clinical and Experimental Medicine IKEM, IN 00023001). We acknowledge Lidia Lopez-Jimenez for her help in generating the iPSC lines and to Radka Kavánova for technical assistance. We would like to thank the patients’ families for their willingness to participate in this study.

## Author Contributions

AH – proteomic experiments, biochemical assays and protein analyses, project coordination, manuscript preparation; LN, MJB – genetic and transcriptomic analyses, data processing, project management, manuscript preparation; JS – design of mouse model experiments, manuscript preparation; SK, TK, DR – manuscript preparation; BMD, ELM, EHSM, GGM, PMR, RCC, EBS, AG, JA, TAB, MJH, DL, RL, DM, LT, FTMT, AV, PD, SC, EFM, CT – clinical description of the patients, phenotyping, contact with the families, genetic analyses; LMM, JA, MP, HH, KH, HT, LM, VS – genomic data analyses; BB, MF – biobanking, fibroblast cultures; DSP, ABN, MHM, SRS, PM- iPSCs establishment and characterization, in vitro assays; VB, AK, PM, KM, MP – biochemical assays, protein analyses; IB, SC, PN, LP, JP, KP, RS, FS, DJ, OF, KN, DPR – mouse model preparation and phenotyping; JK, JR, BS, SG, HMC, MLT, MAL, SB, CC, SH – epigenetic analyses

## Competing interests

B.S. is a shareholder in Episign, Inc. The other authors declare no conflicts of interest

## Ethics Declaration

Informed consent for genetic analyses was obtained for all individuals, and genetic studies were performed as approved by the review boards of corresponding institutes. The patient’s parents provided written informed consent for the participation in the study, clinical data and specimen collection, genetic analysis and publication of relevant findings. All data were de-identified. The study adheres to the principles set out in the Declaration of Helsinki. The generation of human iPSCs was approved by the ethical committee of Instituto de Salud Carlos III CEI PI 03_2022. Informed consents for sample donation and iPSCs generation were signed by the legal guardians.

## Notes

### Summary of Updates

Updated Author List, Main Text, Figures, Supplementary Figures, Methods, Tables

## References

1. Greer, E.L. & Shi, Y. Histone methylation: a dynamic mark in health, disease and inheritance. Nat Rev Genet 13, 343–57 (2012).

2. Younesian, S., Yousefi, A.M., Momeny, M., Ghaffari, S.H. & Bashash, D. The DNA Methylation in Neurological Diseases. Cells 11(2022).

3. Al Ojaimi, M. et al. Disorders of histone methylation: Molecular basis and clinical syndromes. Clin Genet 102, 169–181 (2022).

4. Jenuwein, T., Laible, G., Dorn, R. & Reuter, G. SET domain proteins modulate chromatin domains in eu- and heterochromatin. Cell Mol Life Sci 54, 80–93 (1998).

5. Wei, H. & Zhou, M.M. Dimerization of a viral SET protein endows its function. Proc Natl Acad Sci U S A 107, 18433–8 (2010).

6. Tachibana, M. et al. Histone methyltransferases G9a and GLP form heteromeric complexes and are both crucial for methylation of euchromatin at H3-K9. Genes Dev 19, 815–26 (2005).

7. Sanchez, N.A., Kallweit, L.M., Trnka, M.J., Clemmer, C.L. & Al-Sady, B. Heterodimerization of H3K9 histone methyltransferases G9a and GLP activates methyl reading and writing capabilities. J Biol Chem 297, 101276 (2021).

8. Demond, H. et al. Multi-omics analyses demonstrate a critical role for EHMT1 methyltransferase in transcriptional repression during oogenesis. Genome Res 33, 18–31 (2023).

9. Ohno, H., Shinoda, K., Ohyama, K., Sharp, L.Z. & Kajimura, S. EHMT1 controls brown adipose cell fate and thermogenesis through the PRDM16 complex. Nature 504, 163–7 (2013).

10. Ciceri, G. et al. An epigenetic barrier sets the timing of human neuronal maturation. Nature 626, 881–890 (2024).

11. Kleefstra, T. et al. Loss-of-function mutations in euchromatin histone methyl transferase 1 (EHMT1) cause the 9q34 subtelomeric deletion syndrome. Am J Hum Genet 79, 370–7 (2006).

12. Kleefstra, T. et al. Further clinical and molecular delineation of the 9q subtelomeric deletion syndrome supports a major contribution of EHMT1 haploinsufficiency to the core phenotype. J Med Genet 46, 598–606 (2009).

13. Yamada, A., Shimura, C. & Shinkai, Y. Biochemical validation of EHMT1 missense mutations in Kleefstra syndrome. J Hum Genet 63, 555–562 (2018).

14. Blackburn, P.R. et al. A Novel Kleefstra Syndrome-associated Variant That Affects the Conserved TPL. J Biol Chem 292, 3866–3876 (2017).

15. Rots, D. et al. Comprehensive EHMT1 variants analysis broadens genotype-phenotype associations and molecular mechanisms in Kleefstra syndrome. Am J Hum Genet 111, 1605–1625 (2024).

16. Balemans, M.C. et al. Reduced exploration, increased anxiety, and altered social behavior: Autistic-like features of euchromatin histone methyltransferase 1 heterozygous knockout mice. Behav Brain Res 208, 47–55 (2010).

17. Balemans, M.C. et al. Hippocampal dysfunction in the Euchromatin histone methyltransferase 1 heterozygous knockout mouse model for Kleefstra syndrome. Hum Mol Genet 22, 852–66 (2013).

18. Balemans, M.C. et al. Reduced Euchromatin histone methyltransferase 1 causes developmental delay, hypotonia, and cranial abnormalities associated with increased bone gene expression in Kleefstra syndrome mice. Dev Biol 386, 395–407 (2014).

19. Iacono, G. et al. Increased H3K9 methylation and impaired expression of Protocadherins are associated with the cognitive dysfunctions of the Kleefstra syndrome. Nucleic Acids Res 46, 4950–4965 (2018).

20. Aref-Eshghi, E. et al. Evaluation of DNA Methylation Episignatures for Diagnosis and Phenotype Correlations in 42 Mendelian Neurodevelopmental Disorders. Am J Hum Genet 106, 356–370 (2020).

21. Carvalho, L.M.L. et al. EHMT2 as a Candidate Gene for an Autosomal Recessive Neurodevelopmental Syndrome. Mol Neurobiol (2024).

22. Frega, M. et al. Neuronal network dysfunction in a model for Kleefstra syndrome mediated by enhanced NMDAR signaling. Nat Commun 10, 4928 (2019).

23. Tachibana, M., Sugimoto, K., Fukushima, T. & Shinkai, Y. Set domain-containing protein, G9a, is a novel lysine-preferring mammalian histone methyltransferase with hyperactivity and specific selectivity to lysines 9 and 27 of histone H3. J Biol Chem 276, 25309–17 (2001).

24. Tachibana, M. et al. G9a histone methyltransferase plays a dominant role in euchromatic histone H3 lysine 9 methylation and is essential for early embryogenesis. Genes Dev 16, 1779–91 (2002).

25. Tachibana, M., Matsumura, Y., Fukuda, M., Kimura, H. & Shinkai, Y. G9a/GLP complexes independently mediate H3K9 and DNA methylation to silence transcription. Embo j 27, 2681–90 (2008).

26. Sobreira, N., Schiettecatte, F., Valle, D. & Hamosh, A. GeneMatcher: a matching tool for connecting investigators with an interest in the same gene. Hum Mutat 36, 928–30 (2015).

27. Laurie, S. et al. The RD-Connect Genome-Phenome Analysis Platform: Accelerating diagnosis, research, and gene discovery for rare diseases. Hum Mutat 43, 717–733 (2022).

28. Levy, M.A. et al. Novel diagnostic DNA methylation episignatures expand and refine the epigenetic landscapes of Mendelian disorders. HGG Adv 3, 100075 (2022).

29. Sadikovic, B. et al. Clinical epigenomics: genome-wide DNA methylation analysis for the diagnosis of Mendelian disorders. Genet Med 23, 1065–1074 (2021).

30. Kerkhof, J. et al. Diagnostic utility and reporting recommendations for clinical DNA methylation episignature testing in genetically undiagnosed rare diseases. Genet Med 26, 101075 (2024).

31. Martí, M. et al. Characterization of pluripotent stem cells. Nat Protoc 8, 223–53 (2013).

32. Suresh Babu, S., et al. Characterization of human induced pluripotent stems cells: Current approaches, challenges, and future solutions. Biotechnol Rep (Amst) 37, e00784 (2023).

33. Mozzetta, C. et al. The histone H3 lysine 9 methyltransferases G9a and GLP regulate polycomb repressive complex 2-mediated gene silencing. Mol Cell 53, 277–89 (2014).

34. Auclair, G. et al. EHMT2 directs DNA methylation for efficient gene silencing in mouse embryos. Genome Res 26, 192–202 (2016).

35. Papait, R. et al. Histone Methyltransferase G9a Is Required for Cardiomyocyte Homeostasis and Hypertrophy. Circulation 136, 1233–1246 (2017).

36. Bian, C., Chen, Q. & Yu, X. The zinc finger proteins ZNF644 and WIZ regulate the G9a/GLP complex for gene repression. Elife 4(2015).

37. Britt, H.M. & Robinson, C.V. Traversing the drug discovery landscape using native mass spectrometry. Curr Opin Struct Biol 91, 102993 (2025).

38. Balan, S. et al. Exon resequencing of H3K9 methyltransferase complex genes, EHMT1, EHTM2 and WIZ, in Japanese autism subjects. Mol Autism 5, 49 (2014).

39. Aref-Eshghi, E. et al. BAFopathies’ DNA methylation epi-signatures demonstrate diagnostic utility and functional continuum of Coffin-Siris and Nicolaides-Baraitser syndromes. Nat Commun 9, 4885 (2018).

40. Plazas-Mayorca, M.D. et al. Quantitative proteomics reveals direct and indirect alterations in the histone code following methyltransferase knockdown. Mol Biosyst 6, 1719–29 (2010).

41. Sridharan, R. et al. Proteomic and genomic approaches reveal critical functions of H3K9 methylation and heterochromatin protein-1γ in reprogramming to pluripotency. Nat Cell Biol 15, 872–82 (2013).

42. Wen, B., Wu, H., Shinkai, Y., Irizarry, R.A. & Feinberg, A.P. Large histone H3 lysine 9 dimethylated chromatin blocks distinguish differentiated from embryonic stem cells. Nat Genet 41, 246–50 (2009).

43. Roopra, A., Qazi, R., Schoenike, B., Daley, T.J. & Morrison, J.F. Localized domains of G9a-mediated histone methylation are required for silencing of neuronal genes. Mol Cell 14, 727–38 (2004).

44. Maze, I. et al. G9a influences neuronal subtype specification in striatum. Nat Neurosci 17, 533–9 (2014).

45. Schaefer, A. et al. Control of cognition and adaptive behavior by the GLP/G9a epigenetic suppressor complex. Neuron 64, 678–91 (2009).

46. Katoh, K., Yamazaki, R., Onishi, A., Sanuki, R. & Furukawa, T. G9a histone methyltransferase activity in retinal progenitors is essential for proper differentiation and survival of mouse retinal cells. J Neurosci 32, 17658–70 (2012).

47. Wilson, C. et al. The Histone Methyltransferase G9a Controls Axon Growth by Targeting the RhoA Signaling Pathway. Cell Rep 31, 107639 (2020).

48. Yamada, A. et al. Derepression of inflammation-related genes link to microglia activation and neural maturation defect in a mouse model of Kleefstra syndrome. iScience 24, 102741 (2021).

49. Liu, N. et al. Recognition of H3K9 methylation by GLP is required for efficient establishment of H3K9 methylation, rapid target gene repression, and mouse viability. Genes Dev 29, 379–93 (2015).

