## Supplementary material for "*De novo EHMT2* variants cause an autosomal dominant *EHMT2*-related Kleefstra syndrome via loss of G9a methyltransferase activity": Methods

#### Identification of *EHMT2* variants

Variants in *EHMT2* were discovered by exome or genome analyses. Detailed data analysis workflow for individual patients is summarized in Supplementary Material.

#### DNA Methylation Data Analysis

##### *Data generation and processing*

Methylation analyses were conducted using the clinically validated EpiSign assay, following previously established protocols[1-4]. All computational analyses were performed using R statistical software[5] (version 4.4.1) along with relevant libraries. Raw intensity data files containing methylated and unmethylated signal values underwent quality control using the SeSAMe package[6]. Standard preprocessing of Illumina microarray data involves dye bias correction, background subtraction, and masking of low-quality probes[7]. Detection p-value-based probe masking is further applied using Infinium I out-of-band signal calibration, with a significance threshold set at $p = 0.05$.

##### *Episignature discovery*

A signature discovery analysis was conducted using six case samples (P1-P6) carrying variants in the SET domain of *EHMT2*. Prior to analysis, probes were excluded if they met any of the following criteria: located on the X or Y chromosomes, targeting CpG sites overlapping known single nucleotide polymorphisms (SNPs), identified as cross-reactive, exhibiting high variability in previous Illumina DNA methylation array versions following manufacturing changes, or listed among EPICv1 probes that were omitted in the updated EPICv2 array.

Matched control samples for the discovery analysis were selected using the 'matchIt' package[8], with matching criteria based on age, sex, and the array type used for methylation profiling. Samples from the EpiSign™ Knowledge Database (EKD) previously identified as contributing to batch effects, as well as those with more than 5% failed probes, were excluded during the matching process. A total of 54 matched controls were included in the analysis, yielding a match ratio of 1:9. Principal component analysis (PCA) was subsequently performed to assess data structure and identify potential outliers.

Differential analysis was conducted on the discovery cohort using the 'limma' package[9]. Methylation beta values were first transformed into M-values via a logit transformation. The data was then modeled using multivariate linear regression, where methylation levels served as predictors, case/control status as the response variable, and estimated blood cell compositions as covariates. To enhance statistical reliability, empirical Bayes moderation was applied, and adjusted p-values were calculated using the Benjamini-Hochberg method to control the false discovery rate. To identify probes that define the episignature, a multi-step ranking process was employed. Initially, probes were ranked based on a composite score derived from the absolute mean methylation difference and the negative logarithm of their adjusted p-value. The top subset of these probes was then selected using ROC curve analysis implemented via the 'caret' package[10]. Highly correlated probes were subsequently removed to reduce redundancy. The resulting probe lists were evaluated using unsupervised clustering, including Euclidean distance-based clustering with Ward's method and multidimensional scaling. Visualization was performed using the 'ggplot2'[11] and 'gplots'[12] packages in R. The probe set that yielded the most distinct clustering was chosen to define the final signature.

The selected probes were utilized to construct a support vector machine (SVM) classification model using the e1071 package[13] as previously described[1,14,15]. For each sample, the classifier generates a Methylation Variant Pathogenicity (MVP) score ranging from 0 to 1. An MVP score approaching 1 indicates a methylation pattern closely resembling the EHMT2 episignature, whereas scores near 0 reflect a pattern similar to that of control samples. The SVM classifier was trained using the discovery case samples, matched controls, 75% of the remaining control samples, and 75% of the neurodevelopmental disorder samples from the EKD. The remaining 25% of control and neurodevelopmental disorder samples were reserved for model testing. SVM hyperparameters, including class weights, regularization parameters, and kernel functions were optimized via grid search.

To evaluate the reproducibility and robustness of the identified methylation profile, six rounds of leave-one-out cross-validation were performed on the discovery case samples. In each round, one of the six EHMT2 samples was excluded from probe selection, and the remaining five were used to train the model. The excluded sample was then used for testing and visualized using heatmaps and multidimensional scaling (MDS) plots. To provide an additional independent assessment, the seventh sample in the cohort was subsequently used as a test validation case.

**Cell cultures**

***Fibroblast culture***

Human dermal fibroblasts cultures were established from skin tissue samples. Briefly, fibroblasts were mechanically isolated by dissecting the dermal layer of the skin and the resulting fragments were incubated at 37°C in Dulbecco's modified Eagle's medium (DMEM) containing 2% of fetal calf serum. Cells were expanded in 75 $cm^2$ culture flasks at 37°C with 5% $CO_2$ and 95% humidity in DMEM containing 10% fetal bovine serum.

***Reprogramming to pluripotency***

Cell lines were generated from cultured fibroblasts or peripheral blood mononuclear cells using the CytoTune iPS 2.0 Sendai Reprogramming Kit (Invitrogen). Growing colonies were manually selected according to morphology and passed for at least seven passages before assessing the presence of viral mRNAs.

***Maintenance of iPSCs***

Established cell lines were grown in Essential E8 Media (Gibco) in Vitronectin (Gibco) coated plates. Cells were split twice a week using ReLeSR (Stem Cell Technologies) at a ratio of 1:4-1:10 in 5 μM Rock inhibitor Y-27632 (Tocris) for 1-3 hours before changing to fresh media.

***RNA preparation and real-time PCR***

RNA was extracted using RNeasy kit (Qiagen) and cDNA prepared using the Maxima First Strand cDNA Synthesis Kit. Oligonucleotides for detecting the expression of transgenes were the ones recommended in the reprogramming kit. Clones free from viral expression were selected for further characterization. Oligonucleotides used to detect the expression of pluripotency genes and reprograming factors have been previously described[16] .

***Immunofluorescence staining***

Cells were fixed using 4% PFA for 20 min at room temperature and washed with phosphate buffered saline (PBS) three times, followed by permeabilization and blocking with 0.25% Triton X-100 and 5% normal horse serum at room temperature for one hour. Subsequently, cells were incubated with primary antibodies (anti-Sox2 AB5603 Sigma and PA1-094 Fisher Scientific (1:400), anti-NANOG AF1997-SP R&D (1:50), anti-Oct3/4 (C-10) sc-5279 Santa Cruz Biotechnology, anti-TRA-1-60 sc-21705 Santa Cruz Biotechnology, anti-Alpha-1-fetoprotein A000829-2 Agilent Dako (1:500), Alpha-Smooth Muscle A5228 Sigma (1:500), TUBB3 Biolegend 801202 (1:500)) overnight at 4°C. The next day, cells were washed with PBS and incubated with secondary antibodies for 1 h at room temperature in the dark and stained with DAPI. Images were captured using a laser confocal microscope (Stellaris 8, Leica Microsystems).

***Embryoid body (EB) differentiation***

For in vitro differentiation, cells were trypsinized using ReLeSR and 50,000 cells per well were seeded into low attachment 96-well v-bottom plates in Essential 8 media and 5 µM Rock inhibitor Y-27632 (Tocris). After two days, EBs were moved from the plate to a 10 cm low attachment plate with fresh Essential 8 media. The next day they were transferred to gelatin-coated dishes and cultured in differentiation media consisting of DMEM media supplemented with 20% fetal bovine serum for up to twenty days.

***Karyotype and genotype***

The generated iPSC lines were free from genetic abnormalities according to the hPSC Genetic Analysis Kit from Stem Cell Technology and karyotype from 20 metaphases. To confirm the presence of the mutations, genomic DNA was extracted with the DNeasy Blood and Tissue Kit from Qiagen and mutations verified by amplification and Sanger sequencing. Oligonucleotides used for amplifying and sequencing the *EHMT1* gene were primer forward: 5'-CTTCTTCTCTGTGGGGCGAG-3' primer reverse: 5'-CACCATAAGCATCAGCATCAGC-3' and the *EHMT2* gene primer forward 5'TCCATATCGCCCATTCCTGC-3' and primer reverse 5'-ATGAAGCGGCTGATGTTGCC-3'.

**RNA-seq analysis**

Total RNA was extracted from fibroblasts and iPSCs using the RNeasy mini kit from Qiagen. RNA-seq was performed at BGI Tech Solutions with two or three biological replicates per condition. Briefly, ribosomal RNAs were removed using a RNase H-based method and first-strand cDNA was generated using random hexamer-primed reverse transcription, followed by a second-strand cDNA synthesis with dUTP instead of dTTP, end repair, A addition and adaptor ligation. The U-labeled second-strand template was digested with Uracil-DNA-Glycosylase (UDG) and amplified by PCR. The resulting library was validated by quality control. The PCR products were then heat denatured and circularized by the splint oligo sequence to generate a single strand circle DNA followed by rolling circle replication to create DNA nanoballs (DNB) for sequencing on the MGI DNBSEQ platform. Raw sequencing data with adapter sequences or low-quality sequences were trimmed or filtered and examined by FastQC for basic quality controls. The sequencing analysis was carried out using Galaxy (https://usegalaxy.eu/). Paired reads were aligned to the human hg19 genome build using STAR[17]. Gene counts were calculated using HTseq-count[18] and differential expression between patients and healthy fibroblasts was interrogated using DEseq[19]. Differentially expressed genes were considered at an adjusted p-value <0.05. PCA plots were generated using the DESeq2 package in R software (version 4.4.1). Enrichment analysis in differentially expressed genes was carried out using DAVID[20]. Scatterplots showing correlations were generated with ggplot2 (https://ggplot2.tidyverse.org) in Galaxy. Pearson's correlation and bubble plots were calculated using SRPlot[21]. Heatmaps were generated using Morpheus (https://software.broadinstitute.org/morpheus/).

**Proteomic mass spectrometry for histone extracts**

Proteomic analyses were performed by Active Motif (Waterloo, Belgium). Histones were acid extracted from a cell pellet containing $2.5x10^6$ cells, derivatized via propionylation, digested with trypsin, newly formed N-termini were propionylated as previously described[22]. Histones were extracted by incubating samples at room temperature for 1 hour in 0.2M sulfuric acid with intermittent vortexing. Histones were then precipitated by the addition of trichloroacetic acid (TCA) on ice, and recovered by centrifugation at 10,000 × g for 5 minutes at 4°C. The pellet was then washed once with 1mL cold acetone/0.1% HCl and twice with 100% acetone, and then air dried in a clean hood. The histones were propionylated by adding 1:3 (v/v) propionic anhydride/2-propanol and incrementally adding ammonium hydroxide to keep the pH around 8, and subsequently dried in a SpeedVac concentrator. The pellet was then resuspended in 100 mM ammonium bicarbonate and adjusted to pH 7-8 with ammonium hydroxide. The histones were then digested with trypsin, resuspended in 100 mM ammonium bicarbonate overnight at 37°C, and dried in a SpeedVac concentrator. The pellet was resuspended in 100 mM ammonium bicarbonate and propionylated a second time by adding 1:3 v/v propionic anhydride/2-propanol and incrementally adding ammonium hydroxide to keep the pH around 8, and subsequently dried in a SpeedVac concentrator. Histone peptides were resuspended in 50 µL of 0.1% TFA and 3 µl were injected in 3 technical replicates in a Thermo Scientific TSQ Quantum Ultra mass spectrometer (Thermo Scientific) coupled with an UltiMate 3000 Dionex nano-liquid chromatography system. The data was quantified using Skyline[23] and represents the percent of each modification within the total pool of that amino acid residue. A total of 92 modification states were quantified including unmodified ones.

**Western blot analyses of fibroblasts and iPSCs**

Cell extracts were prepared in a lysis buffer (0.9× CelLytic MT reagent supplemented with 200 mM NaCl, 1.5 mM magnesium chloride, 20 mM EDTA and protease inhibitor cocktail) with gentle shaking for 30 minutes at 4 °C. Crude cell extracts were centrifuged for 16,000×g for 10 minutes at 4 °C. Supernatants were collected and protein concentration was determined using Bradford assay with bovine serum albumin as standard. Subsequently, extracts were analyzed using SDS-PAGE (4–12% gradient gels) followed by western blotting (with 20 µg and 10 µg of total protein loaded in case of fibroblasts and iPSCs, respectively) with immunodectection against G9a (rabbit anti-G9a, 1:1000; Sigma, G 6919), GLP (rabbit anti-GLP, 1:1000, Sigma, 09-078) and GAPDH (mouse Anti-GAPDH, 1:10000; Sigma G8795) as a reference.

**Immunoprecipitation (IP) of G9a**

Dynabeads Protein A (ThermoFisher Scientific) were coated with rabbit anti-G9a or rabbit anti-PFAS IgG as a negative control (1 ug per 50 ul of Dynabeads suspension; ab185050 and ARP46181 from Abcam and Aviva Systems Biology, respectively) in PBS containing 0.1% Tween 20 for 10 minutes at room temperature. Whole cell extracts from iPSCs (100 µg of total protein) were added to antibody-coated Dynabeads and incubated with rotation for 45 minutes at 4 °C. After three washing cycles (0.9× CelLytic MT reagent supplemented with 200 mM NaCl, 1.5 mM magnesium chloride and 20 mM EDTA), immunoprecipitated proteins were eluted by 50 mM Glycine (pH 2.8) followed by an addition of 100 mM Tris-HCl (pH 7.5). The eluate was analysed by western blotting against G9a and GLP (see above) and by mass spectrometry.

Mass spectrometric analysis of eluates from IP

Tris(2-carboxyethyl)phosphine (TCEP) and 2-chloroacetamide were added to final concentrations of 10 mM and 40 mM, respectively, and samples were incubated at 70 °C for 10 min. Samples were then added to Sera-Mag Carboxylate SpeedBeads (Cytiva), diluted with ethanol to a final concentration of 50% (v/v) and incubated for 10 min on a rocker at 1000 rpm to allow protein binding. Beads were washed three times with 80% (v/v) ethanol. Proteins bound to beads were digested with Lys-C (Promega) for 2 h at 37°C, followed by the addition of trypsin (Promega) and overnight digestion at 37°C. Peptides were desalted using homemade StageTips containing three punches of Empore SPE C18 disks (Merck).

LC–MS/MS analysis was performed using a Vanquish Neo UHPLC system (Thermo Fisher Scientific) coupled to a timsTOF SCP mass spectrometer (Bruker Daltonics). Peptides were separated on a reversed-phase PepSep C18 analytical column (15 cm × 75 µm; Bruker Daltonics) using solvent A (2% ACN, 0.1% formic acid) and solvent B (80% ACN, 0.1% formic acid). A 60-min linear gradient from 1% to 35% solvent B was applied. Data were acquired in data-dependent acquisition (DDA) mode.

Raw data were analyzed using PEAKS 12.5 software with a parent mass error tolerance of 20 ppm and a fragment mass error tolerance of 0.05 Da. Carbamidomethylation of cysteine was specified as a fixed modification, while N-terminal acetylation and methionine oxidation were set as variable modifications, with a maximum of three variable modifications per peptide. Database searches were performed against the human UniProt database. This analysis was performed in three replicates.

### Functional studies

#### *In-silico structural mapping*

Mutations were mapped onto the crystal structure of G9a SET-domain bound to the N-terminal peptide of histone H3 and S-adenosylmethionine (PDB ID 5JIN)[24]. Structural models were visualized and assessed by Pymol Viewer 2.1 (Schrodinger, LLC.). Representative images were prepared using UCSF ChimeraX 1.6.1[25].

#### *Preparation of recombinant proteins*

The methyltransferase domains of G9a (aa residues 913-1193) and GLP (aa 1001-1266) with N-terminal hexa-histidine tag followed by recognition site for TEV protease was expressed in *E.coli* (BL21-CodonPlus (DE3)-RIPL strain) using pET28a plasmid. For co-expression of GLP (aa residues 710-1285) and G9a (aa residues 601-1193), both constructs comprising N-terminal hexa-histidine tags with recognition site for TEV protease, the ankyrin repeats and the methyltransferase domain in pETDuet-1 were used. Bacterial culture was cultivated in Terrific Broth medium supplemented by 0.1 mM zinc chloride. When cells reached OD600 of 1.0, protein expression was induced by the addition of 0.5 mM IPTG at 15 °C for 24 hours.

After cell lysis by sonication in lysis buffer (G9a: 20 mM potassium phosphate (pH 7.5) containing 500 mM NaCl, 20 mM imidazole, 10 mM beta-mercaptoethanol, 5% glycerol and SigmaFast inhibitor cocktail; GLP: 50 mM Tris-HCl (pH 7.5) containing 500 mM NaCl, 20 mM imidazole, 0.5 mM TCEP, 10% glycerol and SigmaFast inhibitor cocktail), proteins were purified by affinity chromatography using an imidazole gradient from 20 to 500 mM over 20 column volumes with Profinity IMAC column (Biorad) followed by cleavage of the histidine tag by TEV-protease overnight at 4 °C (protein:protease w/w ratio of 50:1). In the next step, proteins were separated using size-exclusion chromatography (Enrich SEC 650 column, BioRad) in SEC buffer (G9a: 50 mM potassium phosphate (pH 7.5) containing 100 mM NaCl, 2 mM dithiotreitol, 20 mM imidazole and 5% glycerol; GLP: 50 mM Tris-HCl (pH 7.5) containing 100 mM NaCl, 0.5 mM TCEP and 10% glycerol) and a second round of affinity chromatography to remove TEV protease containing uncleavable histidine tag. Finally, proteins were dialyzed against storage buffer (G9a: 50 mM potassium phosphate (pH 7.0) containing 100 mM NaCl, 1 mM TCEP and 5% glycerol; GLP: 25 mM Tris-HCl (pH 8.0) containing 50 mM NaCl, 0.5 mM TCEP and 10% glycerol), and stored in -80C. The GLP–G9a heterodimer was purified using the same protocol as for G9a, as described above. Aminoacid sequence of the purified proteins was verified using on-line pepsin digestion followed by LC-MS/MS analysis (details described in[26]).

***Solubility test of GLP mutants expressed in E.coli***

Bacterial pellets were resuspended in the lysate buffer (as described above) and cells were disintegrated by sonication or by an addition of BugBuster reagent, lysozyme (0.5 mg/ml) and benzonase (25 mU/µl) followed by gentle shaking at room temperature for 20 minutes. Soluble and insoluble fractions were separated by centrifugation at 4 °C at 16,000 G for 20 minutes. Amount of soluble GLP proteins in supernatant was determined using western blotting (with 10 µg of total protein loaded) followed by immunodetection using Anti-6× His tag primary antibody (ab9108 from Abcam) and Rabbit IgG (H+L) Secondary Antibody (31460 from Thermo Fisher Scientific). Insoluble fraction was washed four times for by BugBuster solution followed by centrifugation 4 °C at 5,000×g for 15 minutes and analysed by SDS-PAGE with Coomassie-Blue staining.

***Bioluminescence-based methyltransferase assay***

Enzymatic activity of G9a, GLP and GLP-G9a variants was assessed using MTase-Glo Methyltransferase Assay (Promega) according to manufacturer instructions. The activity assay was performed with a peptide derived from the N-terminal part of histone H3 (aa residues 1-20; manufactured by Merck Life Science) or mononucleosome (prepared according to[27]). Concentration of the histone H3 peptide and S-adenosylmethionine was kept at 10 μM. Methyltransferase reactions were performed at room temperature for 30 minutes with 5 nM G9a or GLP enzymes. In assay with mononucleosome, concentration was kept at 0.5 μM and 5 μM for mononucleosome and SAM, respectively. Reaction was executed at 37 °C for 1 hour with 0.5 μM enzyme. For the GLP–G9a heterodimer, nucleosome methylation assays were performed at room temperature for 1 hour with 0.25 μM enzyme. Obtained signals were recorded using Spark plate reader (TECAN). Quantification was performed using external calibration standard of S-adenosyl homocysteine solution. Kinetic data were fitted according to Michaelis-Menten equation using OriginPro 2015 (OriginLab).

***MALDI-TOF MS-based methyltransferase assay***

Reactions were performed in 50 mM HEPES (pH 7.9) containing 0.5 mM DTT, 0.25 mM PMSF and 2 mM magnesium chloride with 25 μM histone H3 peptide, 50 μM S-adenosylmethionine and G9a or GLP enzyme (25 or 100 nM) for 30 min at 30 °C. The reaction was quenched by the addition of 0.1% TFA. The resulting mixture was 10-fold diluted in 0.1% TFA, mixed with saturated solution of α-cyano-4-hydroxycinnamic acid in a ratio of 1:1 (v/v) and deposited onto MALDI target plate (MTP 384 target plate ground steel, Bruker).. Samples were measured using Autoflex Speed (Bruker Daltonics) in a reflecton positive mode with parameters as follows: mass range 700-3500 m/z, frequency 1000 Hz and ion extraction delay

130 ns. Instrument was externally calibrated using Bruker Peptide calibration standard mix II. Spectra were processed in FlexAnalysis 3.4 (Bruker Daltonics).

***Differential scanning fluorimetry***

Proteins (3 μM) dissolved in an assay buffer (G9a: 50 mM potassium phosphate (pH 7.0) containing 100 mM NaCl, 1 mM TCEP; GLP: 25mM Tris-HCl (pH 8.0) containing 50 mM NaCl and 0.5 mM TCEP) were analyzed in a free state or bound to histone H3 peptide (50 or 500 μM) or S-adenosylmethionine (50 or 500 μM). After preincubation of the G9a proteins with ligands at room temperature for 10 minutes, 5× Sypro-Orange dye (Invitrogen) was added to the samples which were subsequently subjected to melting analysis using CFX96 Real-Time System (BioRad). The proteins were heated from 20 °C to 90 °C in increments of 0.5 °C and with 1-minute hold intervals. The signal was monitored using fluorescence detection (excitation wavelength of 470 nm and emission wavelength of 570 nm). The melting temperatures (Tm) of the proteins were determined as minima from first derivative curves.

***Native mass spectrometry***

In order to examine the protein dimerization and substrate binding, we performed native MS analysis on a Waters Synapt G2Si instrument. Analysed G9a, GLP and GLP-G9a variants were first transferred into 150 mM ammonium acetate pH 7.5 containing 0.5 mM DTT by one or two cycles of spin gel filtration (Micro Bio-Spin P-6 Gel columns, 6-kDa cut off; Bio-Rad). Next, the protein samples were diluted to 5 μM concentration, optionally mixed with SAM or histone H3 peptide in an equimolar ratio, and incubated for 10 min at 4 °C. The samples were electrosprayed at 0.9-1.2 kV from an in-house prepared borosilicate glass spraying tips[28]. (Kwik-Fil 1B120F-4, World Precision Instruments) pulled with P-97 platinum-wire Flaming/Brown Micropipette puller (Sutter Instrument) and coated with 40 nm thick layer of gold using an ACE600 sputter coater (Leica). Data acquisition was performed in positive ionization sensitivity mode. For assessing the dimerization and substrate binding of the G9a and GLP variants, the trap collision energy was optimized to maximize sensitivity while keeping ion activation minimal at 10 V and 40 V, respectively. The argon flow as collision gas was maintained at 6 ml/min, sampling cone voltage set to 80 V, source offset to 0 V, and source temperature to 80 °C. Quadrupole was operated in broad transmission mode up to 4000 m/z. Acquired data were externally mass recalibrated on cesium iodide clusters, summed over 50 scans, and smoothed using two passes of Savitzky-Golay filtering (window 50 scans)[29] in MassLynx 4.1 (Waters). Subsequently, mass deconvolution of the spectra was performed in UniDec v6.0.4 (10.1021/acs.analchem.5b00140). Additionally, spectra were background subtracted in UniDec v6.0.4 and used to quantify the monomer-dimer distribution of the analyzed proteins in their apo-form (Supplementary Figure 12).

### Mouse model

#### *Generation of the knock-in mouse model*

To model the effects of the variant (c.3225_3236del, p.Glu1076_Val1079del) identified in patient 1, the *Ehmt2* c.3385_G3396del (p.E1129_V1132del, (C57BL/6NCrl-Ehmt2$^{em1Ccpcz}$/Ph) mouse model (further referenced in the manuscript as *Ehmt2*$^{+/del_1076-1079}$) was generated on the C57BL/6NCrl background (Charles River Laboratories) by targeting exon 25 (ENSMUSE00001282962) (Supplementary Figure 15A). *Ehmt2* codons E1129, A1130, D1131 and V1132, that are complementary to human *EHMT2* codons 1076-1079, were deleted using CRISPR/Cas9 technology in combination with single-stranded oligonucleotides (ssODN) template: 5'-GGTGCTGCATCCCACCTTGTTATCTAAATCGAAGAGGTAAGAATCATCCTCTCTC GCATCAGAGATCAGCTCTCCTACGTACCTGTTGGACGGAAACTTCAGTTCTGCAT GAACAGTGAGC -3' as described by[30]. The guide RNAs (gRNAs) of highest score and specificity were designed using http://crispor.tefor.net/.The following guide was selected:

gRNA- AGAGCTGATCTCTGATGCCG. The gRNAs (100ng/µl, Integrated DNA Technologies) were assembled into a ribonucleoprotein (RNP) complex with Cas9 protein (500ng/µl; 1081058, 1072532, Integrated DNA Technologies), electroporated into 1-cell zygotes, and transferred into ICR (CD1) pseudopregnant foster females. Putative founders were analyzed by PCR and sequencing. A founder carrying a E1129-V1132del mutation also harbors *SnaBI* (Eco105I) restriction enzyme site enabling direct detection of the mutation by the restriction enzyme digest. Genotyping was performed by PCR with forward (F) 5′-TTCCCTCAACTTCTCAGACACC -3′ and reverse (R)5’- ACCCGGACAGGGATGATGTT -3’ primers. The F and R product is 775 bp, *SnaBI* digestion of the E1129-D1132del homozygote mice PCR product results in 439bp and 324bp fragments (Supplementary Figure 15B, C). Selected founder was confirmed by sequencing and bred with C57BL/6NCrl wild-type to confirm germ-line transmission of the target deletion. The only male founder carrying E1129-V1132del gave rise to infertile females and sub-viable males, often dying in 10 days after birth. Due to this limitation, the founder was archived in form of the frozen sperm. To propagate this line, in vitro fertilization was performed as described previously[31] and viable embryos we implanted into ICR (CD1) surrogate females (Charles River Laboratories). Animals born after IVF were genotyped and sequenced (Sanger sequencing) within 5 days after birth to detect and monitor positive mice. Bulk sequencing data were further analyzed using DECODR.org online software[32]. The sperm from founder male was sequenced using NGS sequencing to detect all possible *Ehmt2* gene variants in germ line.

Evaluators were blinded to the genotype of individual mice in all the following tests.

***SHIRPA and locomotor activity scoring***

Standardized SHIRPA scoring scale[33] was used to generally asses the phenotype of the $Ehmt2^{+/del\_1076\text{-}1079}$ and $Ehmt2^{+/+}$ mice. Locomotor Activity Test was used for quick evaluation of spontaneous exploratory behavior and general activity level of mice. The test was performed in a room with 100-120 lux light intensity using a Maneko rat cage (55 × 33 cm) with a floor divided into 15 equal squares. Each mouse was placed in the arena and allowed to move freely. Locomotor activity was quantified by counting the number squares the animal entered during the first 30 seconds of exploration.

***In-vivo microCT mouse scanning***

All mice were anesthetized via intramuscular administration of 20% Zoletil-Xylazine (Virbac). In-vivo scans of mice were performed using the SkyScan 1278 (Bruker) as continuous 7 connected scans. Scanning parameters included a voxel size of 50 µm, a 0.5 mm aluminum filter, 180° rotation and no averaging. Source voltage was set to 54 kV and source current to 909 µA. Data reconstruction was carried out using NRecon software, version 2.2.0.6 (Bruker) with the following parameters: ring artifact correction at 3, beam hardening at 28%, threshold for defect pixel mask at 5%, no smoothing and intensity range set from 0.0025 to 0.10 AU[34]. CTvox software, version 3.3.0.0 (Bruker) was used for data visualization. Craniometric measurements (Supplementary Figure 17) were performed in DataViewer, version 1.5.2 (Bruker).

***Open Field test***

The activity of the animals in a novel environment and the level of anxiety displayed were evaluated in the open field test[35]. The area of the open field was a square of 42 × 42 cm uniformly illuminated with a light intensity of 200 lux in the center of the field. The testing arena was virtually divided into periphery and center zones, where the center zone constituted 38% of the whole arena. Each mouse was placed in the corner of the arena for a 20 min period of free maze exploration. The time spent in each zone, the distance travelled, and other indices were automatically computed based on video recordings (Viewer software 3.0.1.452, Biobserve GmbH).

***Gait analysis***

Gait analysis was conducted using the DigiGait system (Mouse Specifics Inc., Framingham, MA, USA) to assess the specific differences in gait between *Ehmt2*$^{+/+}$ (males *n* = 15 and females *n* = 15) and *Ehmt2*$^{+/del\_1076\text{-}1079}$ (males *n* = 8 and females *n* = 6) mice at 3-4 months of age.

Briefly, the DigiGait system consisted of a transparent treadmill, horizontally fixed at 0 ° (5 cm in width, 25 cm in length), which contained the mice as they ran at a constant velocity. The velocity selected for the assessment was fixed at 13 cm/s, a speed at which the *Ehmt2*$^{+/del\_1076\text{-}1079}$ animals were able to consistently ambulate throughout the experiment. A minimum of five fluent strides were recorded during each trial. A high-temporal-resolution video was recorded (~5 s) for each mouse and video analysis was performed using the DigiGait Imaging and Analysis software ver. 16 (Mouse Specifics Inc., Framingham, MA, USA). The software automatically analyzed images to create a digital paw print and dynamic gait signals. Spatial and temporal gait parameters were measured from the generated signals. Data were averaged between left and right paws in both the forelimbs and hindlimbs. To acclimatize to the apparatus, mice were trained on the DigiGait treadmill at speeds of 10 to 15 cm/s for a maximum of 1 min per mouse for 5 days prior to testing. The average of 3 days was used for subsequent calculations.

***Grip strength***

A grip strength meter was used to determine the maximal peak force (×g) exerted by the mouse forelimbs as well as all four limbs when the examiner tried to pull it out of a specially designed grid for mice (Bioseb, BIO-GS3 & BIO-GRIPGS, Bioseb). The maximal effort (g) was used as absolute force (×g) and was corrected for body mass (×g/×g). The experiment was conducted as a singly-anonymized trial. and a total of 3 measurements were applied to each mouse with a 10-15 min interval between measurements. The average of the 3 values was used for subsequent calculations.

#### *Rotarod*

The rotarod system was used to assess the sense of balance, motor learning, and motor coordination in the animals. Three trials per day for each mouse were recorded on a rod with an accelerating speed of rotation (4–40 rpm/5 min, RotaRod, TSE Systems) during two consecutive days. The average latency to fall was determined from three trials with 15 min intertrial intervals (ITI).

#### *Echocardiography*

Transthoracic echocardiography was performed using a high-resolution ultrasound imaging system (Vevo® F2, FUJIFILM VisualSonics) equipped with an ultra-high-frequency transducer (VisualSonics UHF46x, 46–20 MHz). Fifteen-week-old animals were anesthetized with 5% isoflurane for induction and positioned in the supine position on a temperature-controlled imaging platform maintained at 38 °C. During image acquisition, the isoflurane concentration was reduced to 1.5–2% to maintain stable and comparable heart rates (425 ± 50 bpm) and respiration rates.

Following the removal of chest hair and the application of pre-warmed ultrasound gel, two-dimensional B-mode cine loops were acquired. Parasternal long-axis (LAX) views were obtained to visualize the left ventricle (LV) from apex to base. Parasternal short-axis (SAX) views were subsequently recorded at the mid-ventricular level, typically at the level of the papillary muscles, to assess LV morphology and function.

After sonography, animals were returned back to their cages placed on heating pad and monitored until full recovery from anesthesia.

All measurements and calculations were made using AutoLV mode with the VevoLab software (FUJIFILM VisualSonics Inc.).


1. Aref-Eshghi, E. *et al.* Diagnostic Utility of Genome-wide DNA Methylation Testing in Genetically Unsolved Individuals with Suspected Hereditary Conditions. *Am J Hum Genet* **104**, 685-700 (2019).
2. Kerkhof, J. *et al.* Diagnostic utility and reporting recommendations for clinical DNA methylation episignature testing in genetically undiagnosed rare diseases. *Genet Med* **26**, 101075 (2024).
3. Levy, M.A. *et al.* Novel diagnostic DNA methylation episignatures expand and refine the epigenetic landscapes of Mendelian disorders. *HGG Adv* **3**, 100075 (2022).
4. Sadikovic, B. *et al.* Clinical epigenomics: genome-wide DNA methylation analysis for the diagnosis of Mendelian disorders. Genet Med 23, 1065-1074 (2021).
5. R & Team, C. R: A Language and Environment for Statistical Computing, (2022).
6. Zhou, W., Triche, T.J., Laird, P.W. & Shen, H. SeSAMe: reducing artifactual detection of DNA methylation by Infinium BeadChips in genomic deletions. *Nucleic Acids Res* **46**, e123 (2018).
7. Zhou, W., Laird, P.W. & Shen, H. Comprehensive characterization, annotation and innovative use of Infinium DNA methylation BeadChip probes. *Nucleic Acids Res* **45**, e22 (2017).
8. Ho, D.a.I.K.a.K.G.a.S.E.A. MatchIt: Nonparametric Preprocessing for Parametric Causal Inference. *Journal of Statistical Software* **42**, 1–28 (2011).
9. Ritchie, M.E. *et al.* limma powers differential expression analyses for RNA-sequencing and microarray studies. *Nucleic Acids Res* **43**, e47 (2015).
10. Kuhn, M. Building Predictive Models in R Using the caret Package. *Journal of Statistical Software* **28**, 1–26 (2008).
11. Wickham, H. Data analysis. in *ggplot2: elegant graphics for data analysis* 189--201 (2016).
12. gplots: Various R Programming Tools for Plotting Data. (The R Foundation, 2005).
13. Dimitriadou, E.a.H.K.a.L.F.a.M.D.a.W.A. *E1071: Misc Functions of the Department of Statistics (E1071), TU Wien*, (2009).
14. Aref-Eshghi, E. *et al.* BAFopathies' DNA methylation epi-signatures demonstrate diagnostic utility and functional continuum of Coffin-Siris and Nicolaides-Baraitser syndromes. *Nat Commun* **9**, 4885 (2018).
15. Aref-Eshghi, E. *et al.* Evaluation of DNA Methylation Episignatures for Diagnosis and Phenotype Correlations in 42 Mendelian Neurodevelopmental Disorders. *Am J Hum Genet* **106**, 356-370 (2020).
16. Aasen, T. *et al.* Efficient and rapid generation of induced pluripotent stem cells from human keratinocytes. *Nat Biotechnol* **26**, 1276-84 (2008).
17. Dobin, A. *et al.* STAR: ultrafast universal RNA-seq aligner. *Bioinformatics* **29**, 15-21 (2013).
18. Anders, S., Pyl, P.T. & Huber, W. HTSeq--a Python framework to work with high-throughput sequencing data. *Bioinformatics* **31**, 166-9 (2015).
19. Anders, S. & Huber, W. Differential expression analysis for sequence count data. *Genome Biol* **11**, R106 (2010).
20. Sherman, B.T. *et al.* DAVID: a web server for functional enrichment analysis and functional annotation of gene lists (2021 update). *Nucleic Acids Res* (2022).
21. Tang, D. *et al.* SRplot: A free online platform for data visualization and graphing. *PLoS One* **18**, e0294236 (2023).
22. Garcia, B.A. *et al.* Chemical derivatization of histones for facilitated analysis by mass spectrometry. *Nat Protoc* **2**, 933-8 (2007).

23. MacLean, B. *et al.* Skyline: an open source document editor for creating and analyzing targeted proteomics experiments. *Bioinformatics* **26**, 966-8 (2010).
24. Jayaram, H. *et al.* S-adenosyl methionine is necessary for inhibition of the methyltransferase G9a by the lysine 9 to methionine mutation on histone H3. *Proc Natl Acad Sci U S A* **113**, 6182-7 (2016).
25. Pettersen, E.F. *et al.* UCSF ChimeraX: Structure visualization for researchers, educators, and developers. *Protein Sci* **30**, 70-82 (2021).
26. Pacheco-Garcia, J.L. *et al.* Structural basis of the pleiotropic and specific phenotypic consequences of missense mutations in the multifunctional NAD(P)H:quinone oxidoreductase 1 and their pharmacological rescue. *Redox Biol* **46**, 102112 (2021).
27. Luger, K., Rechsteiner, T.J. & Richmond, T.J. Expression and purification of recombinant histones and nucleosome reconstitution. *Methods Mol Biol* **119**, 1-16 (1999).
28. Hernández, H. & Robinson, C.V. Determining the stoichiometry and interactions of macromolecular assemblies from mass spectrometry. *Nat Protoc* **2**, 715-26 (2007).
29. Savitzky, A.a.G.M.J.E. Smoothing and Differentiation of Data by Simplified Least Squares Procedures. *Analytical Chemistry* **36**, 1627-1639 (1964).
30. Inui, M. *et al.* Rapid generation of mouse models with defined point mutations by the CRISPR/Cas9 system. *Sci Rep* **4**, 5396 (2014).
31. Takeo, T. & Nakagata, N. Combination medium of cryoprotective agents containing L-glutamine and methyl-{beta}-cyclodextrin in a preincubation medium yields a high fertilization rate for cryopreserved C57BL/6J mouse sperm. *Lab Anim* **44**, 132-7 (2010).
32. Bloh, K. *et al.* Deconvolution of Complex DNA Repair (DECODR): Establishing a Novel Deconvolution Algorithm for Comprehensive Analysis of CRISPR-Edited Sanger Sequencing Data. *CRISPR J* **4**, 120-131 (2021).
33. Rogers, D.C. *et al.* Behavioral and functional analysis of mouse phenotype: SHIRPA, a proposed protocol for comprehensive phenotype assessment. *Mamm Genome* **8**, 711-3 (1997).
34. Spoutil, F. *et al.* Semi-Automated MicroCT Analysis of Bone Anatomy and Mineralization in Mouse Models. *Curr Protoc* **4**, e980 (2024).
35. Syding, L.A. *et al.* Generation and Characterization of a Novel Angelman Syndrome Mouse Model with a Full Deletion of the. *Cells* **11**(2022).
