## Supplementary Figure for "*De novo EHMT2* variants cause an autosomal dominant *EHMT2*-related Kleefstra syndrome via loss of G9a methyltransferase activity"

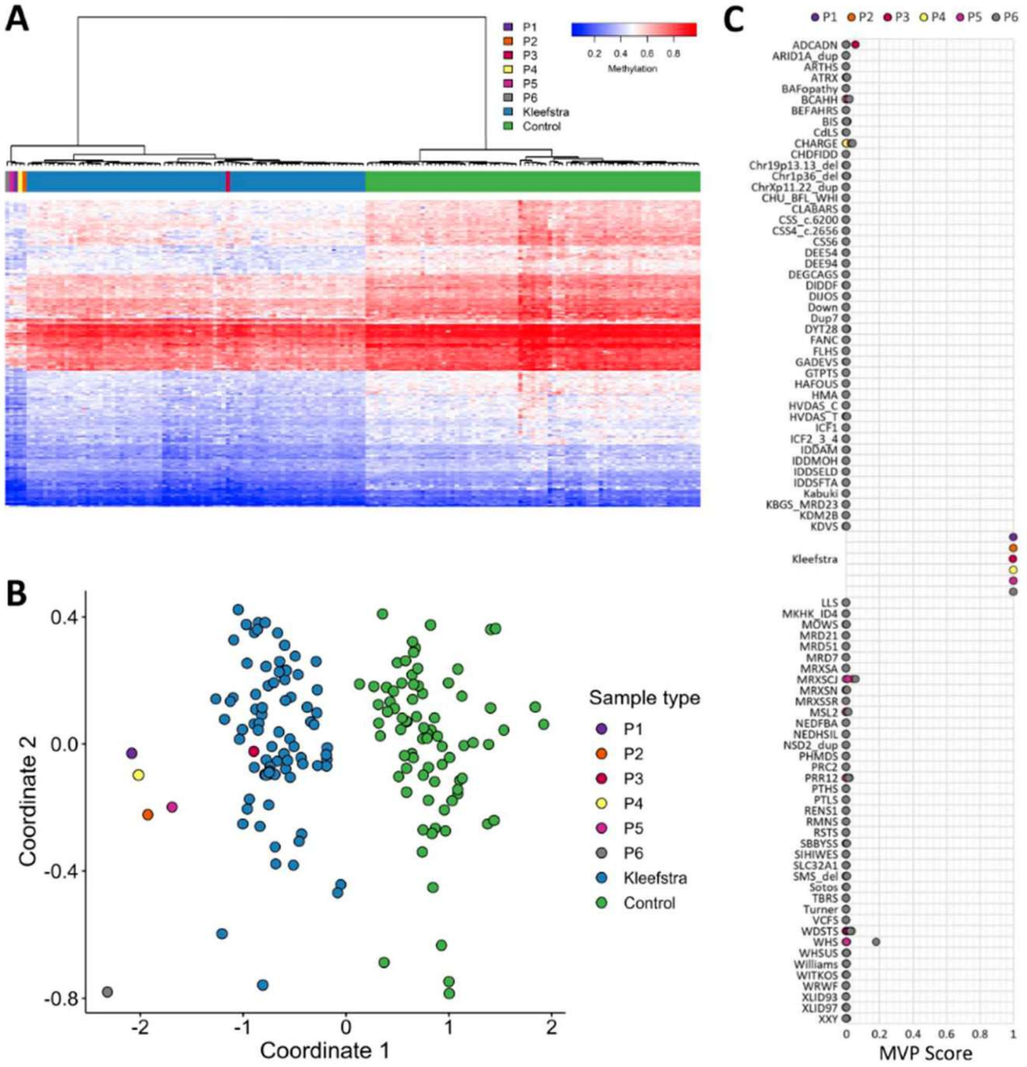


**Supplementary Figure 1**. EpiSign (DNA methylation) analysis of peripheral blood from patients with EHMT2 variants.

(A) Hierarchical clustering and (B) multidimensional scaling plots indicate the patients (purple, orange, red, yellow, pink and grey) have a DNA methylation profile similar to subjects with a confirmed Kleefstra episignature (blue) and distinct from controls (green). (C) MVP score, a multi-class supervised classification system capable of discerning between multiple episignatures by generating a probability score for each episignature. The elevated score for Kleefstra shows an episignature similar to reference patients with pathogenic variants in EHMT1.

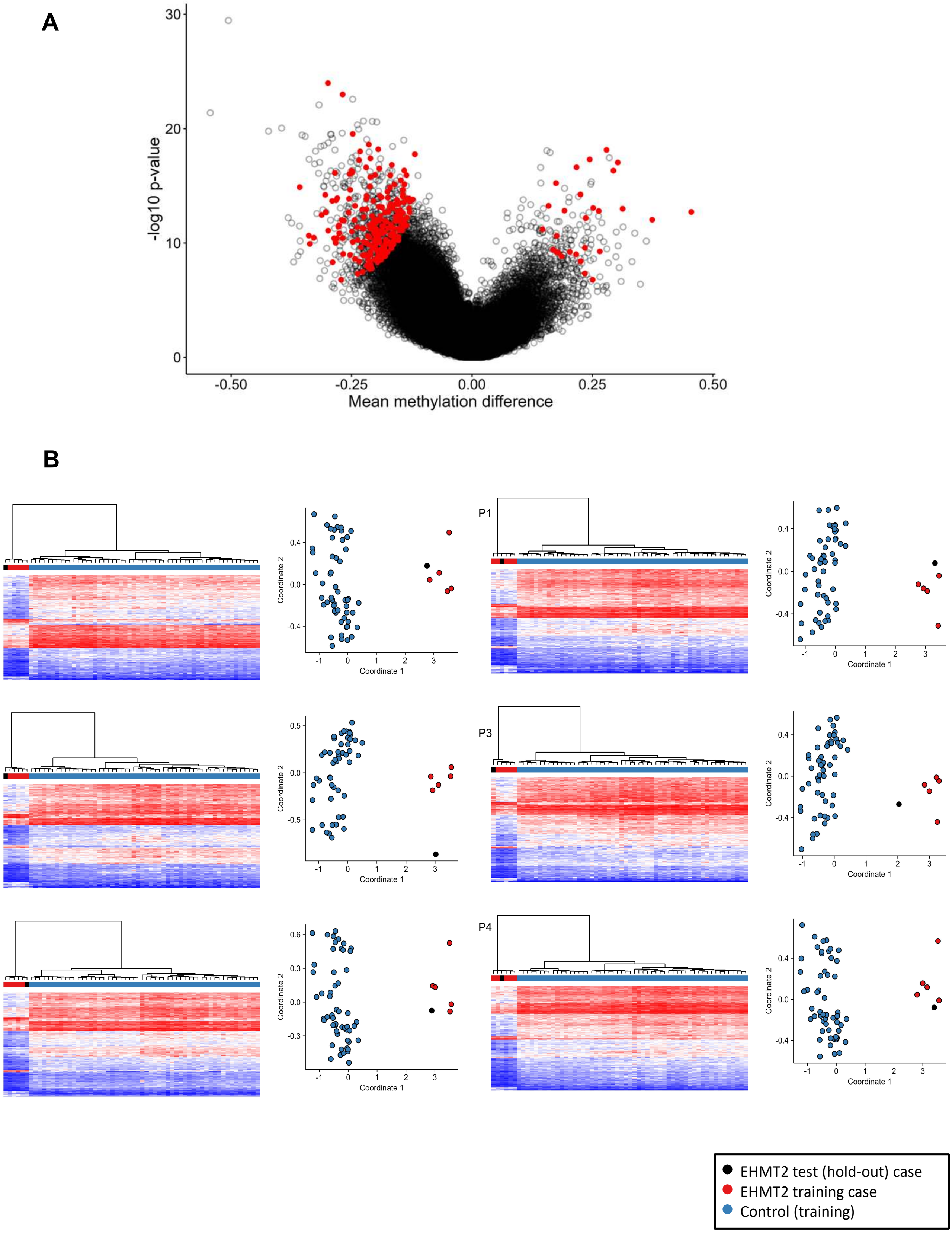


**Supplementary Figure 2**. A. Visualization of mean methylation differences and p-values for the episignature probes. The volcano plot displays all episignature probes, with mean methylation differences on the x-axis and –log10 p values on the y-axis. Probes selected through the multi-step ranking procedure for the EHMT2 episignature are highlighted in red.

B. Unsupervised clustering results for leave-one-out cross-validation on PTBP1 discovery cases. In each cross-validation iteration, a single PTBP1 start-loss case was held-out from the discovery data as a test case and a sub-signature was generated using the same feature selection parameters of the identified signature. Reproducibility and robustness of the episignature is validated by consistent clustering of the hold-out case (black) with the remaining discovery cases (red) instead of the matched controls (blue).

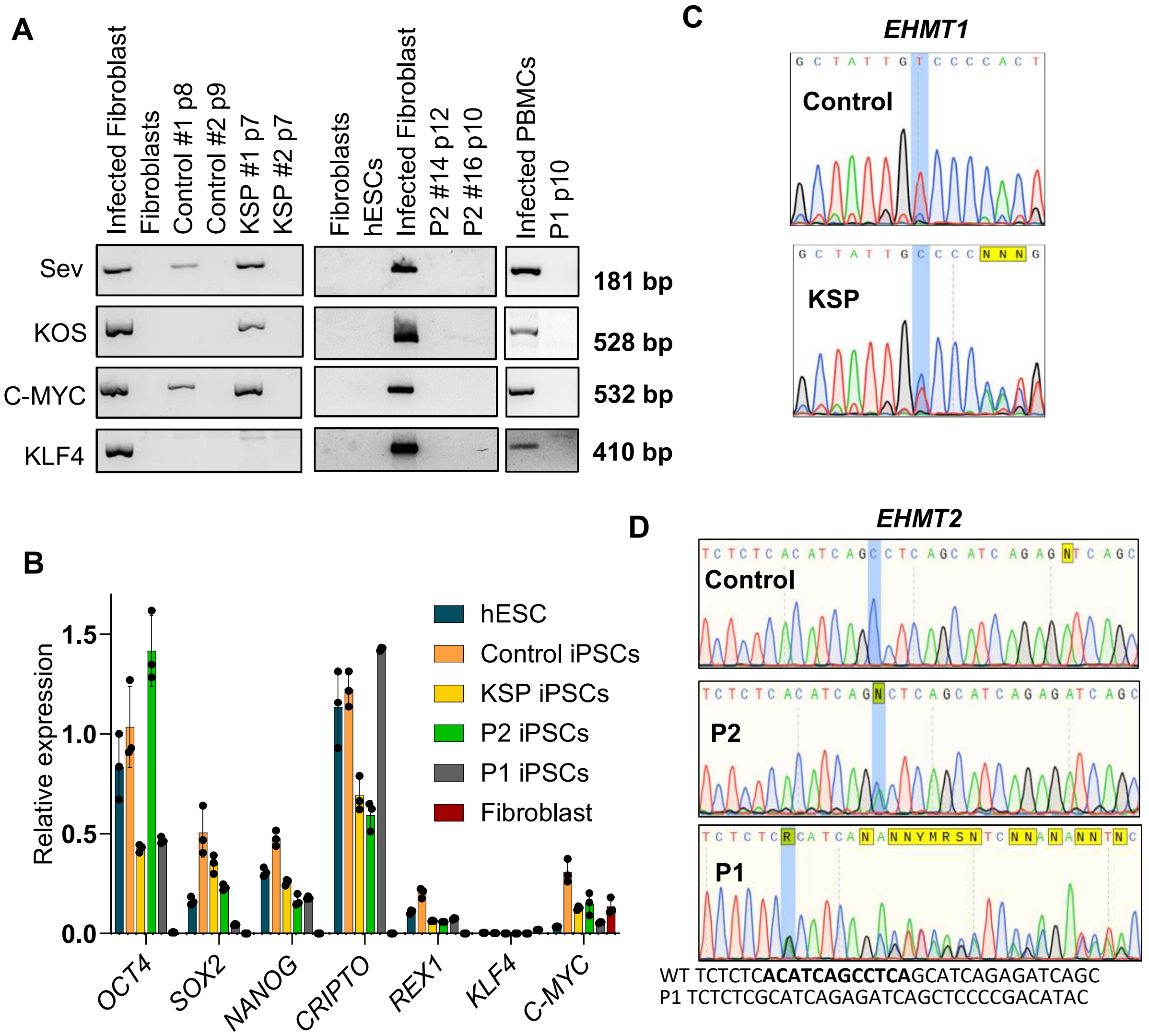


**Supplementary Figure 3**. Characterization of iPSCs lines. A. Transgenes expression in the different lines at the indicated passages. Transgene-free clones Control #2, KSP #2 and P2 #16 were selected for subsequent analysis. B. Pluripotency markers expression in the indicated cell lines obtained by qPCR. Three replicates are shown. The human embryonic stem cell hESC (AND-2) is included as a positive control. C. Sanger sequencing of EHMT1 mutation in control and KSP iPSC lines D. Sanger sequencing of EHMT2 mutations in control and P1 and P2 iPSC lines. For patient P1 missing nucleotides are highlighted in bold.

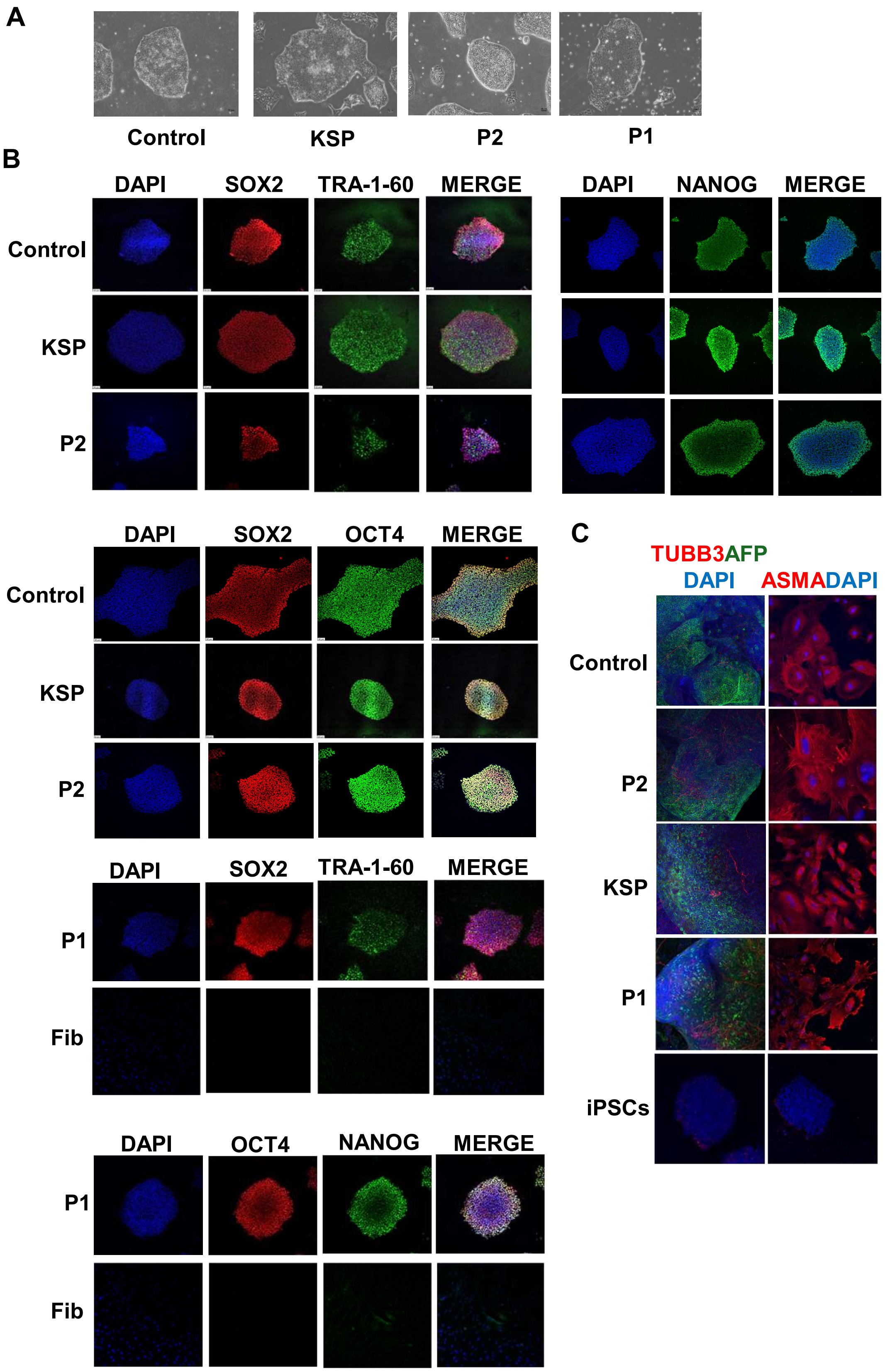


**Supplementary Figure 4.** Characterization of iPSCs lines by immunochemistry. A. Phase contrast pictures of growing cell lines. B. Immunohistochemistry of pluripotency markers in the indicated cell lines. All pictures correspond to iPSCs except negative controls performed in fibroblasts (Fib) C. Detection of differentiation markers in EBs at day 20 of in vitro differentiation. Negative controls were performed in undifferentiated iPSCs.

**A**

| Histone PTM | Log2 P2/C | Log2 KSP/C |
|---|---|---|
| H1.4: k25me2 | -1.87 | 0.28 |
| H3: k79ac | -1.74 | -0.14 |
| H3R2un: q5me1 | -0.73 | -0.50 |
| H3: k9me2 | -0.71 | -0.53 |
| H1.4: k25me1 | -0.66 | -1.01 |
| H3: k9me3 | -0.60 | -0.55 |
| H3: k79un | -0.60 | -0.30 |
| H3.3: k27ac | -0.53 | 0.22 |
| H3.3: k36un | -0.48 | -0.56 |
| H4: k20un | -0.38 | 0.07 |
| H3: q55me1 | -0.23 | 0.38 |
| H3.1: k27un | -0.21 | 0.21 |
| H3: k9me1 | -0.18 | -0.23 |
| H3.1: k36un | -0.17 | -0.28 |
| H3: k79me1 | -0.14 | -0.20 |
| H4: k20me1 | -0.14 | 0.03 |
| H3: k14un | -0.12 | -0.08 |
| H3.3: k36me1 | -0.11 | -0.56 |
| H3.3: k36me3 | -0.11 | 0.19 |
| H3R2un: k4me1 | -0.07 | 0.08 |
| H4: k16un | -0.06 | -0.05 |
| H3: r49un | -0.05 | -0.06 |
| H3: k23un | -0.05 | -0.04 |
| H3.3: k27me1 | -0.04 | -0.12 |
| H3: k122ac | -0.04 | 1.54 |
| H4: k8un | -0.04 | -0.04 |
| H2A: k5ac | -0.04 | 0.04 |
| H3: r42un | -0.04 | -0.03 |
| H3.1: k27me2 | -0.03 | -0.22 |
| H4: k12un | -0.03 | -0.03 |
| H4: k5un | -0.02 | -0.01 |
| H3: k18un | -0.02 | -0.03 |
| H3R2un: k4me2 | -0.01 | -0.07 |
| H3.1: k27me1 | -0.01 | 0.08 |
| H3: k56un | 0.00 | 0.00 |
| H2A: k9un | 0.00 | 0.00 |
| H2A3: k15un | 0.00 | 0.00 |
| H3: q19un | 0.00 | 0.00 |
| H2A: k36un | 0.00 | 0.00 |
| H3: k64un | 0.00 | 0.00 |
| H2A1: k13un | 0.00 | 0.00 |
| H2A1: k15un | 0.00 | 0.00 |
| H2A3: k13un | 0.00 | 0.00 |
| H3: k122un | 0.00 | 0.00 |
| H3: q55un | 0.00 | 0.00 |
| H2A: k5un | 0.00 | 0.00 |
| H3R2un: q5un | 0.01 | 0.00 |
| H3R2un: k4un | 0.01 | -0.01 |
| H3.3: k27me3 | 0.01 | -0.10 |
| H1.4: k25un | 0.02 | 0.03 |
| H4: k20ac | 0.04 | 0.05 |
| H3.1: k36me3 | 0.04 | 0.26 |
| H3.1: k36me2 | 0.06 | 0.20 |
| H3.3: k27un | 0.07 | 0.19 |
| H3.3: k27me2 | 0.09 | -0.16 |
| H4: k16ac | 0.13 | 0.11 |
| H3.1: k27ac | 0.13 | 0.05 |
| H4: k20me2 | 0.16 | -0.20 |
| H3.3: k36ac | 0.16 | 0.26 |
| H3: k79me2 | 0.22 | 0.14 |
| H3: k18ac | 0.23 | 0.42 |
| H3: k18me1 | 0.24 | -0.14 |
| H3: k23ac | 0.24 | 0.24 |
| H3.3: k36me2 | 0.25 | 0.15 |
| H2A3: k13ac | 0.27 | 0.32 |
| H3: k14ac | 0.28 | 0.19 |
| H3.1: k36me1 | 0.28 | 0.02 |
| H3: k9un | 0.33 | 0.30 |
| H3.1: k27me3 | 0.38 | 0.13 |
| H2A: k9ac | 0.41 | 0.41 |
| H4: k5ac | 0.46 | 0.28 |
| H3: q19me1 | 0.48 | -0.07 |
| H3: k9ac | 0.50 | 0.70 |
| H3: k56me1 | 0.50 | 0.85 |
| H3: k23me1 | 0.50 | 0.77 |
| H2A: k36ac | 0.59 | -0.51 |
| H3.1: k36ac | 0.63 | 1.36 |
| H4: k8ac | 0.63 | 0.62 |
| H2A3: k15ac | 0.78 | 2.22 |
| H2A1: k15ac | 0.83 | 0.41 |
| H4: k12ac | 0.86 | 0.88 |
| H4: k20me3 | 0.90 | 0.72 |
| H3R2un: k4me3 | 0.93 | 0.45 |
| H3R2un: k4ac | 0.95 | -0.33 |
| H2A1: k13ac | 0.99 | 0.28 |
| H1.4: k25ac | 1.09 | 1.15 |
| H3: r49me2 | 1.10 | 1.22 |
| H3: r42me2 | 1.42 | 1.33 |
| H1.4: k25me3 | 1.60 | 0.51 |
| H3: k56ac | 2.37 | 1.73 |
| H3: k64ac | 2.46 | 2.73 |
| H3: k79me3 | 2.97 | 2.27 |

**B**

| Histone PTM | Log2 FIB/iPSCs |
|---|---|
| H3: q19me1 | -4.56 |
| H3: k64ac | -4.21 |
| H3: q55me1 | -3.66 |
| H4: k20me3 | -3.62 |
| H2A: k9ac | -3.53 |
| H1.4: k25me3 | -3.51 |
| H4: k12ac | -3.16 |
| H4: k8ac | -2.83 |
| H3.1: k27ac | -2.78 |
| H3R2un: k4ac | -2.52 |
| H4: k20me1 | -1.80 |
| H3: k18ac | -1.80 |
| H4: k20me2 | -1.72 |
| H3: k23ac | -1.65 |
| H2A: k5ac | -1.63 |
| H4: k5ac | -1.52 |
| H3.1: k36me2 | -1.41 |
| H4: k16ac | -1.17 |
| H3.1: k27un | -1.09 |
| H2A3: k15ac | -0.87 |
| H3: k14ac | -0.75 |
| H3: k9ac | -0.74 |
| H3.3: k27ac | -0.73 |
| H3: k23me1 | -0.59 |
| H3.3: k36me2 | -0.59 |
| H2A: k36ac | -0.51 |
| H3: k79un | -0.51 |
| H3.1: k27me1 | -0.45 |
| H3: k9un | -0.33 |
| H3.3: k27un | -0.31 |
| H3R2un: k4me1 | -0.27 |
| H3R2un: k4me2 | -0.20 |
| H3.3: k27me1 | -0.16 |
| H3: k9me2 | 0.19 |
| H3: k18me1 | 0.20 |
| H2A: k5un | 0.23 |
| H3: k18un | 0.25 |
| H3: k79me1 | 0.29 |
| H4: k12un | 0.31 |
| H3.3: k36me1 | 0.35 |
| H3.1: k36me1 | 0.39 |
| H3.1: k36me3 | 0.41 |
| H3: k9me3 | 0.41 |
| H3: k14un | 0.42 |
| H3.3: k36un | 0.44 |
| H3: k9me1 | 0.46 |
| H3.3: k36me3 | 0.46 |
| H4: k8un | 0.49 |
| H3: k23un | 0.64 |
| H3: r49me2 | 0.68 |
| H4: k16un | 1.06 |
| H3.1: k36ac | 1.08 |
| H3.3: k27me2 | 1.08 |
| H3: r42me2 | 1.17 |
| H1.4: k25me1 | 1.32 |
| H3.3: k27me3 | 1.37 |
| H4: k20un | 1.54 |
| H3.1: k36un | 1.64 |
| H3.3: k36ac | 1.65 |
| H3: k56me1 | 1.68 |
| H3.1: k27me2 | 1.88 |
| H3: k79me2 | 2.96 |
| H3.1: k27me3 | 3.36 |
| H4: k20ac | 3.66 |
| H1.4: k25me2 | 5.12 |
| H3: k79me3 | 5.16 |

**Supplementary Figure 5.** Fold change of histone modifications determined by mass spectrometry. A. Log2 of fold change in the indicated modifications in P2 or KSP fibroblasts relative to healthy control fibroblasts. B. Log2 of fold change in the indicated modifications in control fibroblasts relative to control iPSCs. Modifications that significantly change at least 10% in abundance at a p-value<0.05 are shown.

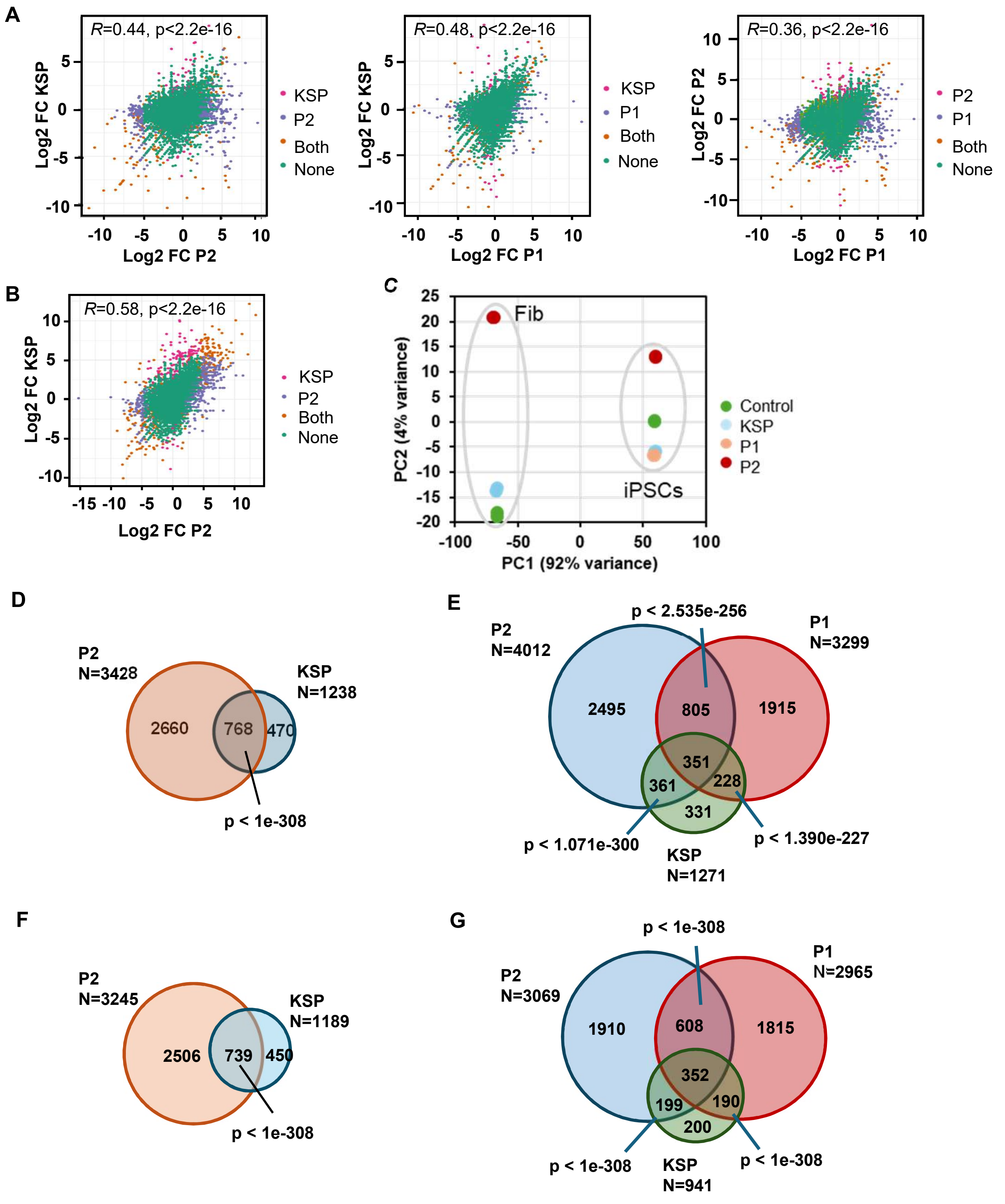


**Supplementary Figure 6 .** Transcriptomic analysis of cell lines derived from a healthy control, a Kleefstra Syndrome patient (KSP) and patients P1 and P2 carrying EHMT2 variants. A. Scatter plot showing correlation between expression fold change of all genes in patients compared to control iPSCs. Pearson's correlation coefficient (R) and p-value is indicated. Genes changing expression at an adjusted p-value<0.05 only in the indicated patients only (pink or purple), in both (orange) or not significantly changing expression (green) are shown. B. Scatter plot showing correlation between expression fold change of all genes in the patients' compared to control fibroblasts. Pearson's correlation coefficient (R) and p-value is indicated. Genes changing expression at an adjusted p-value<0.05 only in KSP (pink), only in P2 (purple), in both (orange) or not significantly changing expression (green) are shown. C. Principal component analysis of healthy and patient's fibroblasts and iPSCs. D. Overlap of differentially upregulated genes compared to control in patient's fibroblasts. P-value shows the significance of the overlap. E. Overlap of differentially upregulated genes compared to control in patient's iPSCs. P-value shows the significance of the overlap. F. Overlap of differentially downregulated genes compared to control in patient's fibroblasts. P-value shows the significance of the overlap. G. Overlap of differentially downregulated genes compared to control in patient's iPSCs. P-value shows the significance of the overlap.

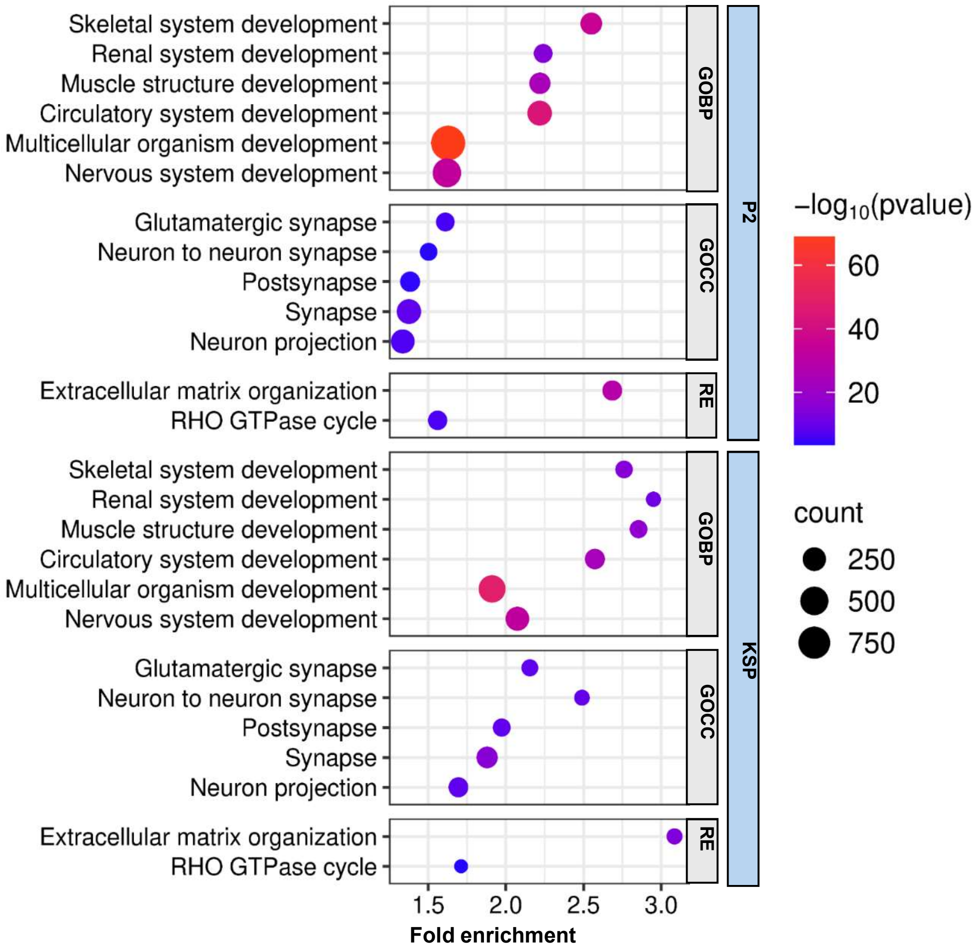


**Supplementary Figure 7.** DAVID enrichments of GOBP, GOCC and Reactome (RE) terms in genes upregulated in KSP or P2 fibroblasts compared to healthy control fibroblasts.

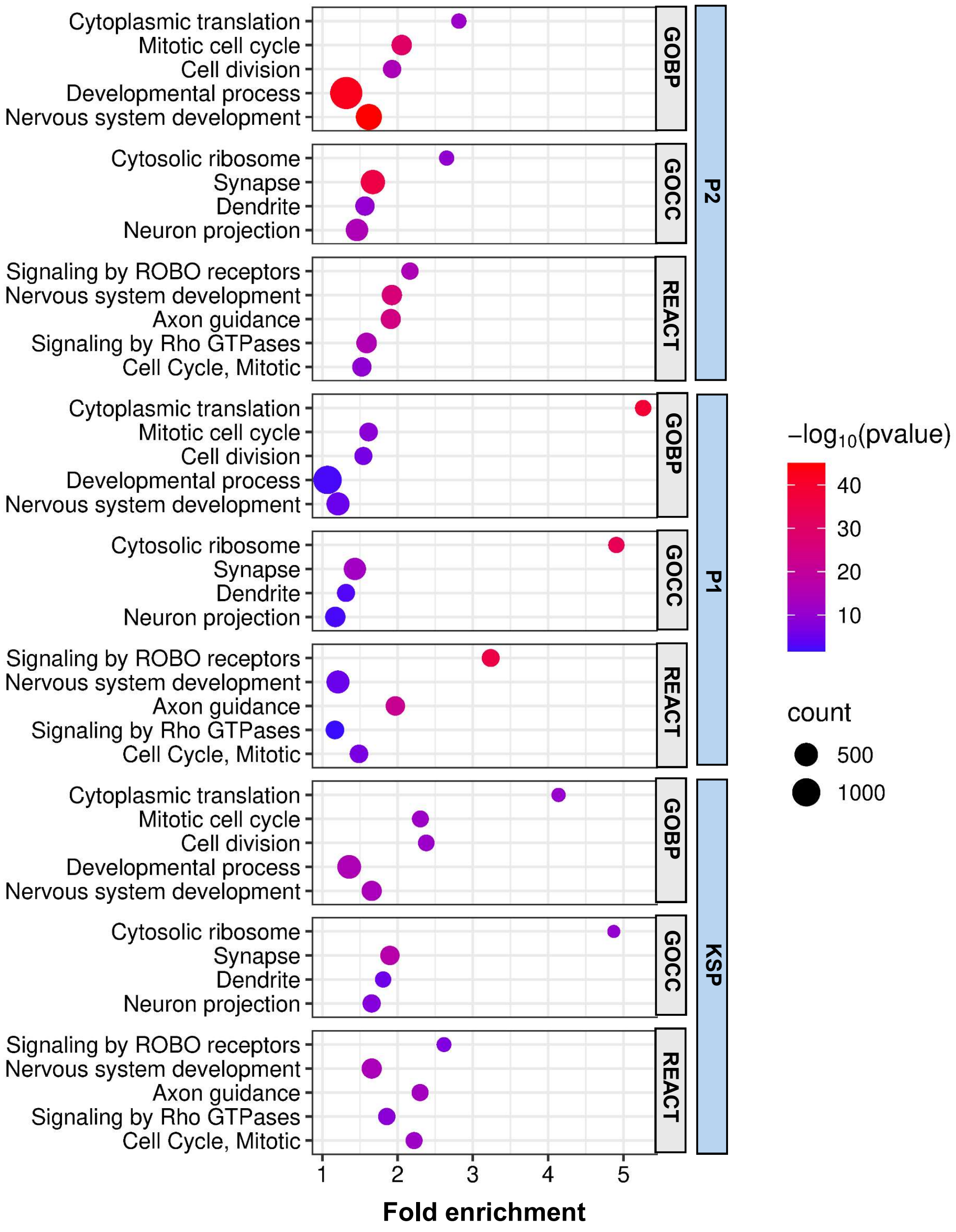


**Supplementary Figure 8 .** DAVID enrichments of GOBP, GOCC and Reactome (REACT) terms in genes upregulated in P2, P1 or KSP iPSCs compared to healthy control iPSCs.

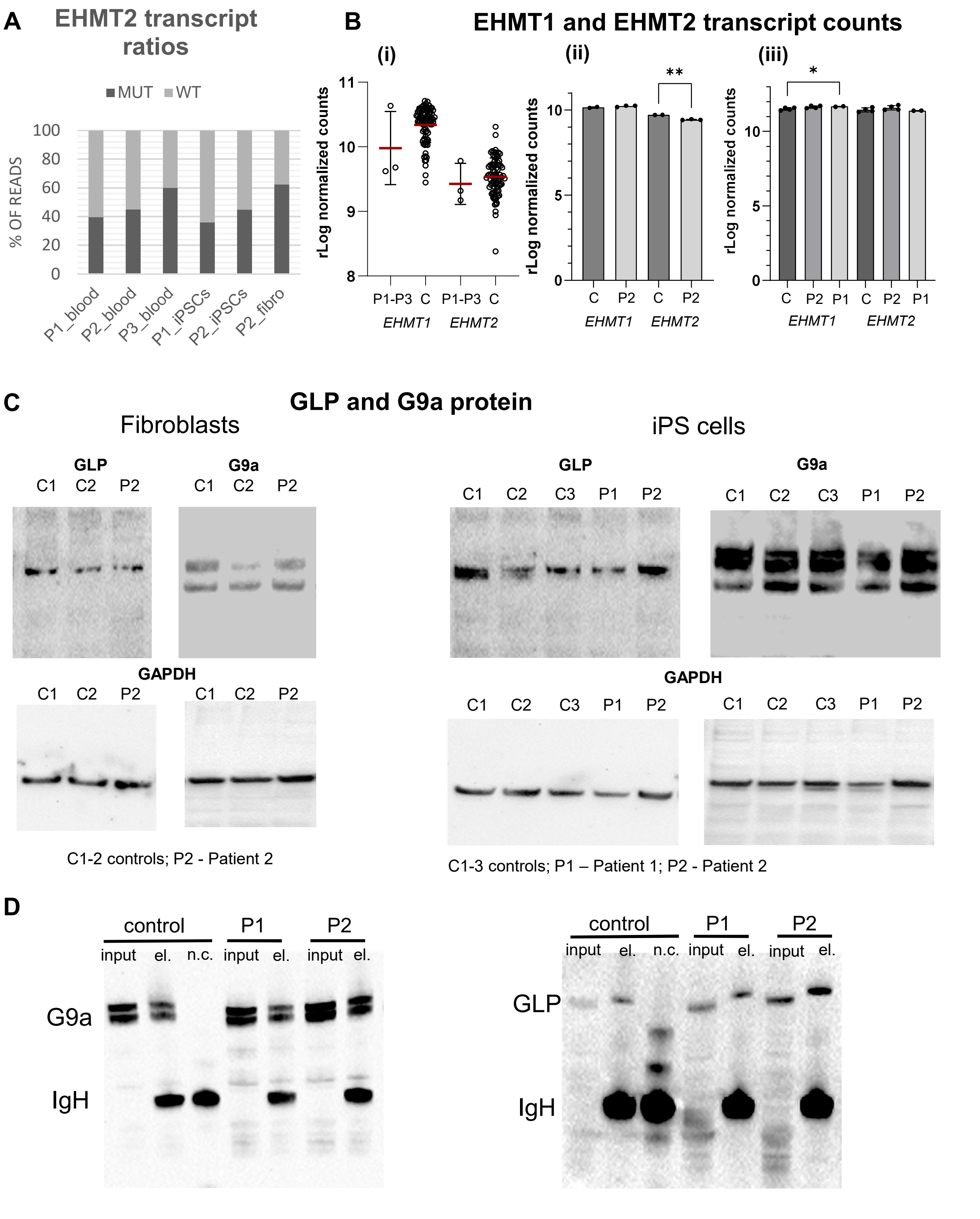


**Supplementary Figure 9.**

A. Percentage of mutant (MUT) and wild type (WT) reads of *EHMT2* transcript detected by RNA-seq from leukocytes of P1, P2 and P3, iPSCs from P1 and P2 and fibroblasts from P2.

B. (i) mRNA levels of EHMT1 and EHMT2 in the blood of patients P1, P2 and P3 and 83 control samples.

(ii) mRNA levels of EHMT1 and in fibroblasts derived from patient P2 (n=3) and control fibroblasts (C) n=2.

(iii) mRNA levels of EHMT1 and in iPSCs derived from patient P2 (n=4), P1 (n=2) and control fibroblasts (n=4). All plots show means with standard deviation. Significant differences by t-test are indicated. * $p<0.05$, ** $p<0.005$

C. Western blot analysis of whole- cell extracts of patient-derived fibroblasts and iPSCs to determine the amount of GLP and G9a protein. GAPDH protein was used as a housekeeping reference.

D. Western blot analysis of G9a immunoprecipitation in iPSCs. “el.” and “n.c.” mean elution and negative control, respectively.

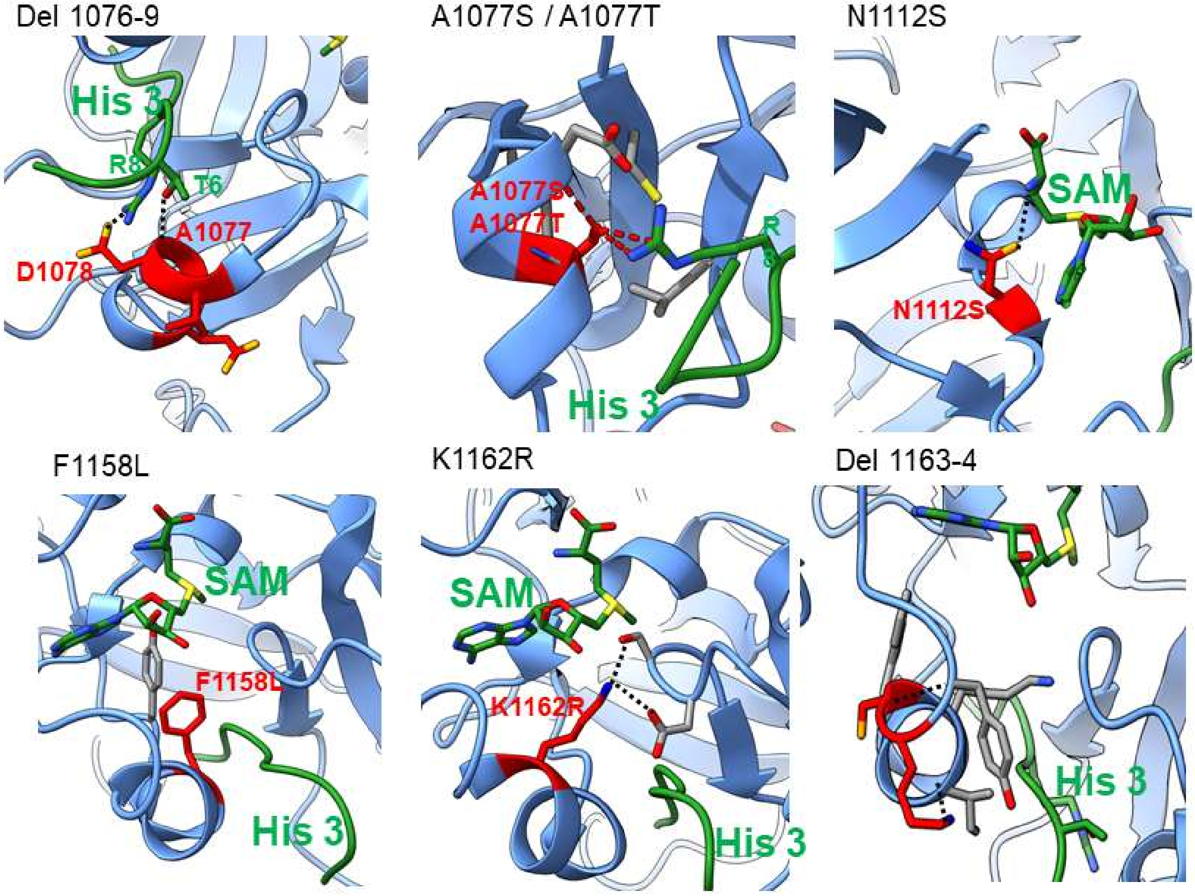


**Supplementary Figure 10.** Structural analyses of G9a mutants using available crystal structure of the G9a SET-domain in complex with SAM and histone 3 peptide (PDB ID 5JIN). Mutated residues are highlighted by red color; lost interactions and sterical clashes are depicted as dashed black and red lines, respectively.

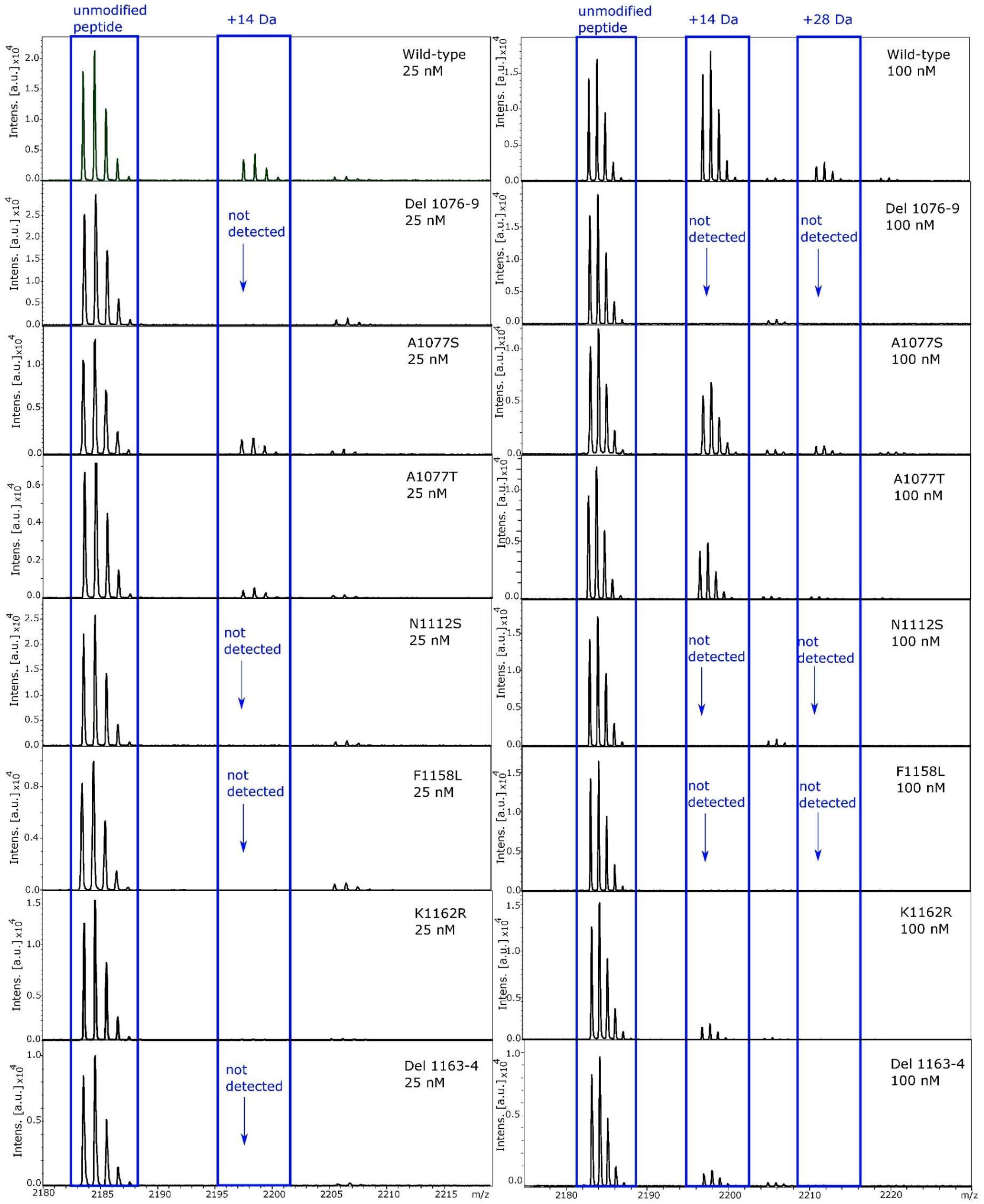


**Supplementary Figure 11.** Methyltransferase activity of G9a variants with histone 3 peptide (aa 1-20) assessed by MALDI-TOF MS. Reactions were performed for 30 minutes at 30 °C with two different concentrations of the enzyme (25 and 100 nM). Methylation of histones 3 peptides causes an increase in molecular mass by 14 (monomethylation) and 28 Da (dimethylation). Series of histone 3 peptides are highlighted by blue boxes. The mutants exhibited undetectable or decreased residual activity compared to the wild-type protein.

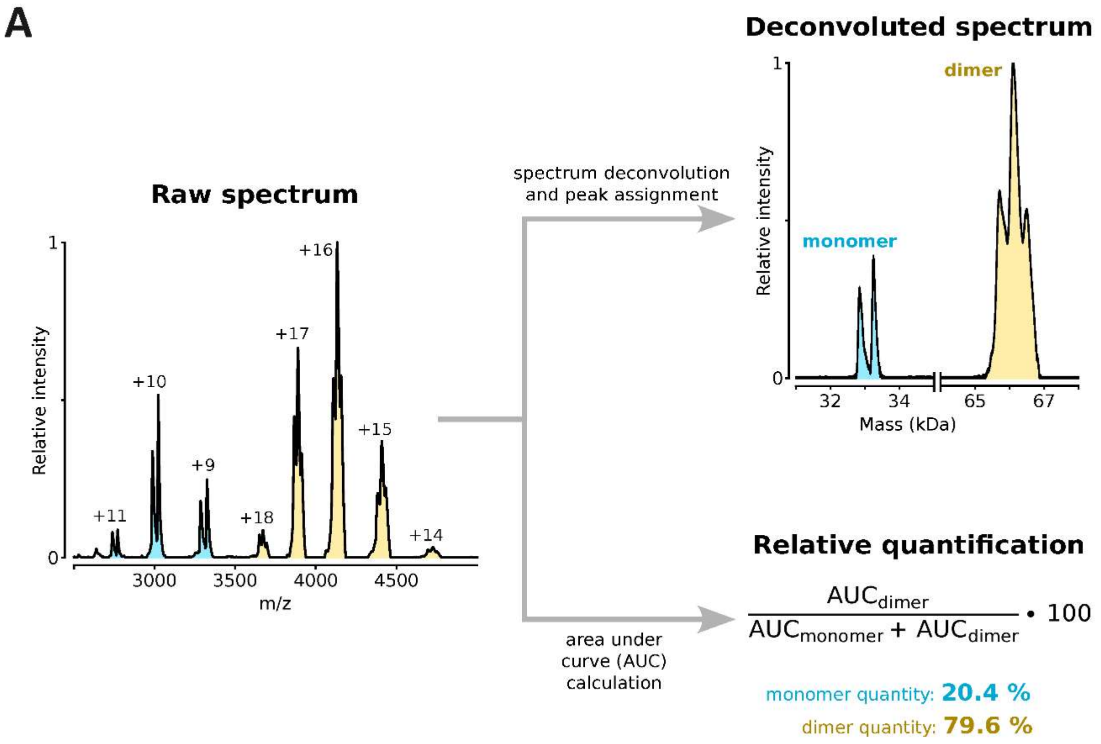


$$\frac{AUC_{dimer}}{AUC_{monomer} + AUC_{dimer}} \cdot 100$$

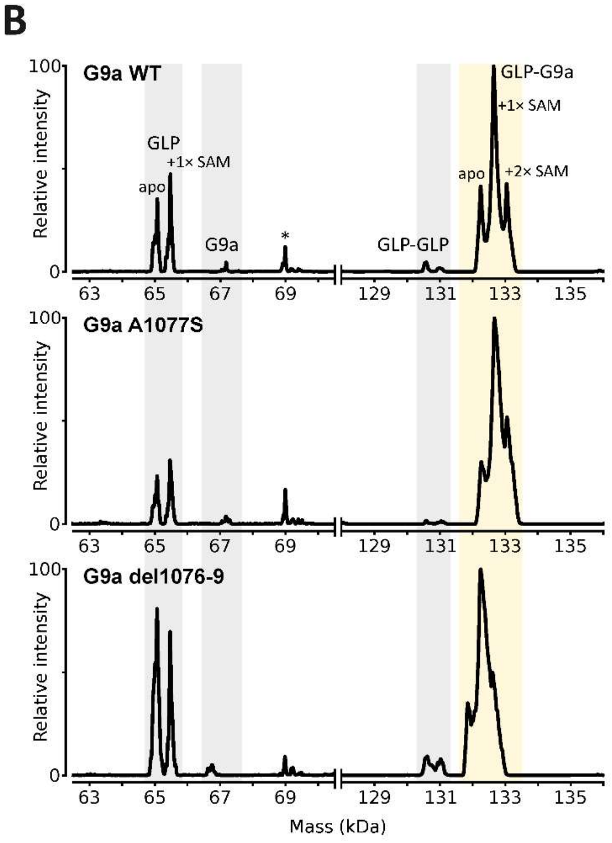


| G9a variant | Relative amount of GLP-G9a heterodimer (%) |
|---|---|
| WT | 80.7 |
| A1077S | 87.3 |
| del1076-9 | 65.5 |

**Supplementary Figure 12.** Mass spectrometric analysis of G9a and GLP dimerization. **A** Evaluation of the monomer-dimer distribution in G9a and GLP variants using nESI-MS. The procedure is illustrated using representative data from methyltransferase domain of the G9a wild-type. Initially, raw spectra were deconvoluted to facilitate peak assignment. Subsequently, relative quantities of monomeric and dimeric forms were calculated from the areas under curve (AUC) of the corresponding peaks in raw MS spectra. B. Deconvoluted nMS spectra of GLP-G9a heterodimer, comprising ankyrin repeat region and methyltransferase domain, with variants in G9a. Yellow area highlights properly assembled heterodimer. Peak denoted with asterisk was not assigned and likely represents protein contaminant from recombinant expression.

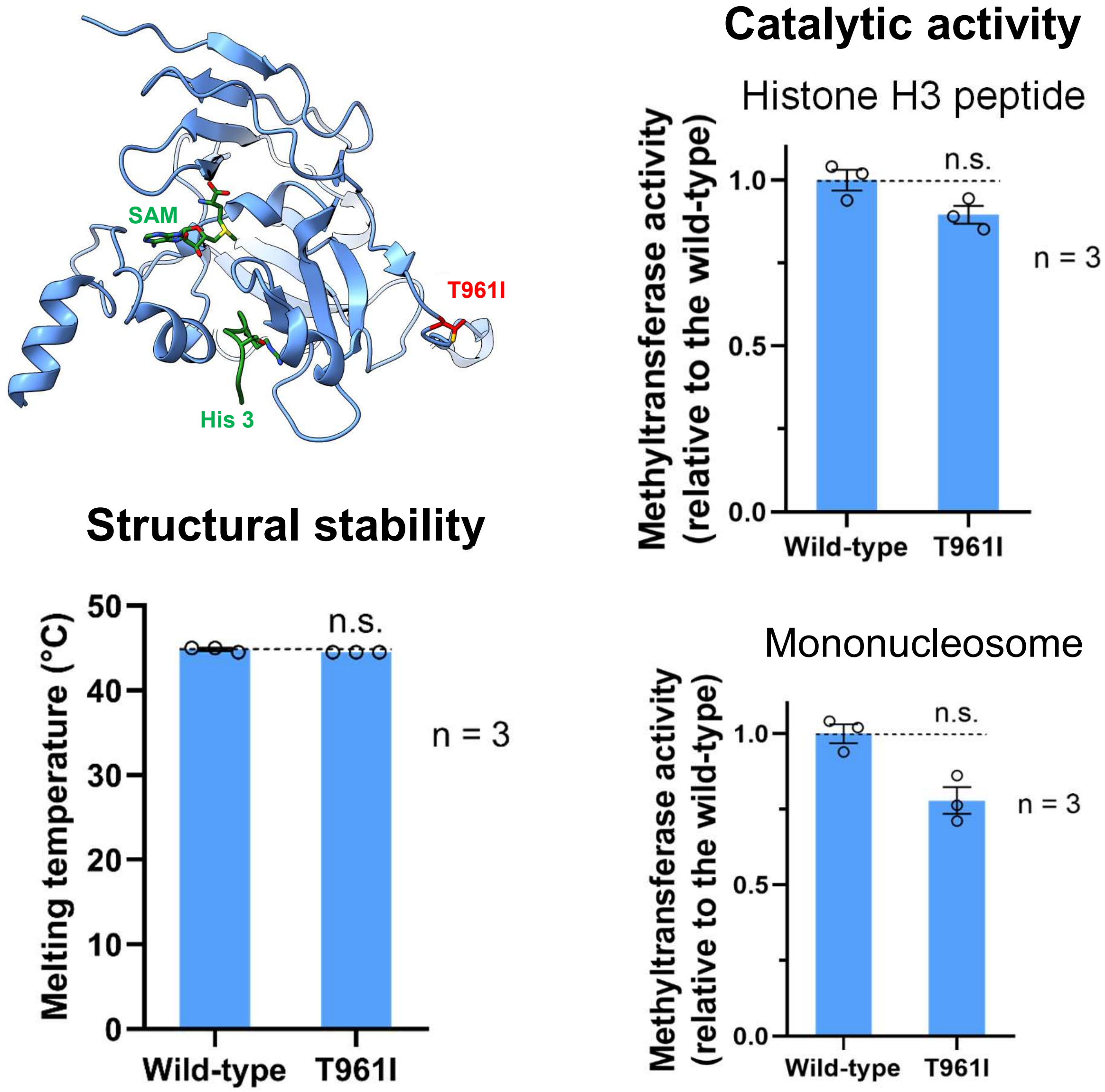


**Supplementary Figure 13.** Structural topology, catalytic activity and structural stability of the variant T961I with unknown clinical significance. Values represent means with standard errors from three measurements. Statistical significance was determined using two-tailed Student's t-test ($p < 0.01$).

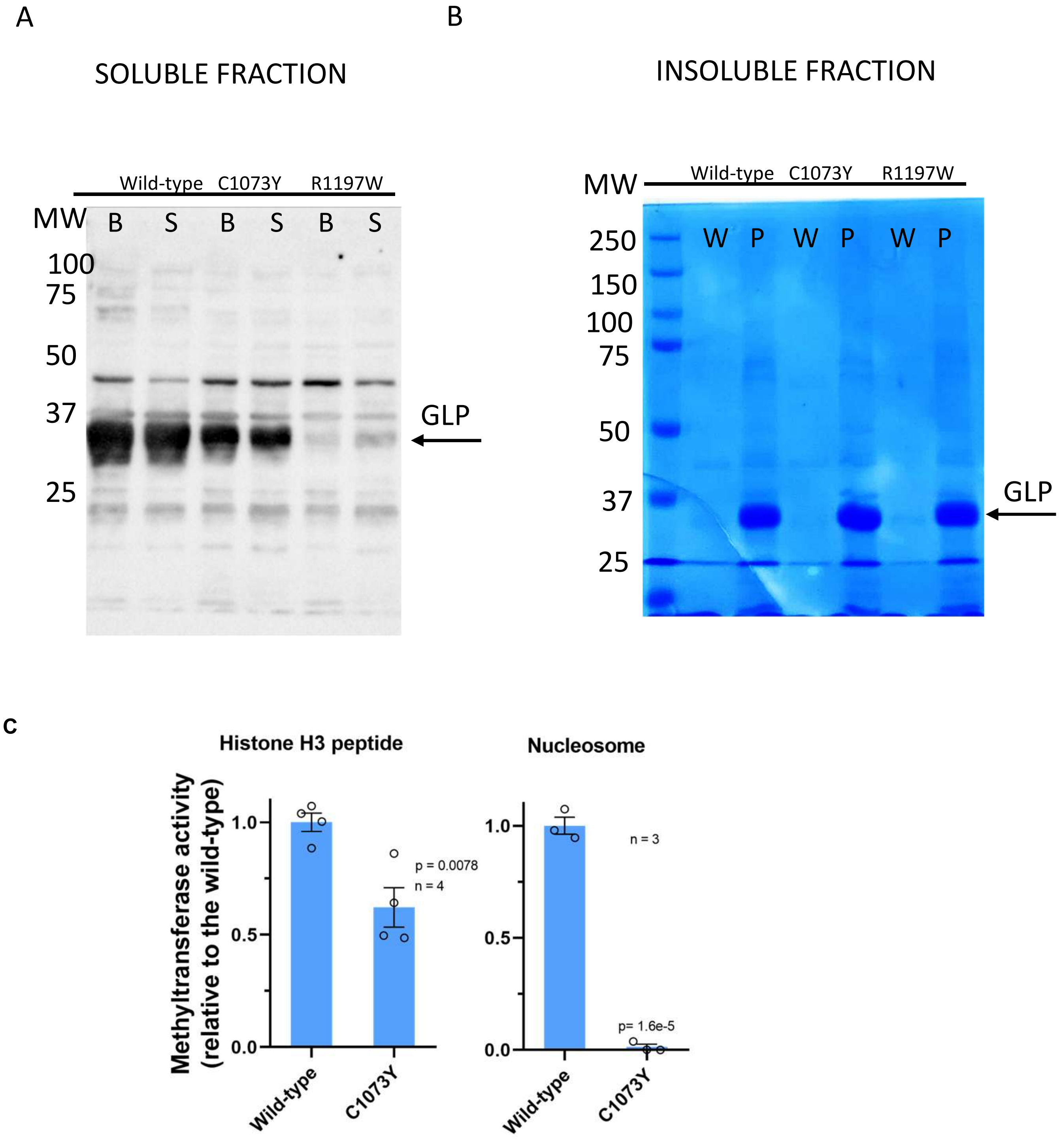


**Supplementary Figure 14.** Production of GLP methyltransferase domain with histidine tag in *E.coli* and its distribution in soluble and insoluble fraction and its catalytic activity. A. Bacterial lysates were prepared by sonication (S) or extraction using BugBuster (B) followed by centrifugation. The soluble fraction represents supernant (10 µg of total protein) analyzed by western blotting followed by immunodetection against histidine tag. B. The insoluble fraction represents pellet after centrifugation analyzed by Coomassie Blue-stained SDS-PAGE (W – wash of insoluble fraction; P – pellet solubilized by heating) (C) Methylation activity of purified GLP variants with histone H3 peptide and mononucleosome as substrates studied by quantitative bioluminescence-based assay (Mtase-Glo by Promega). The values represent the means of relative activity to the wild-type enzyme with standard errors. Statistical significance was determined using two-tailed Student's t-test ($p < 0.01$).

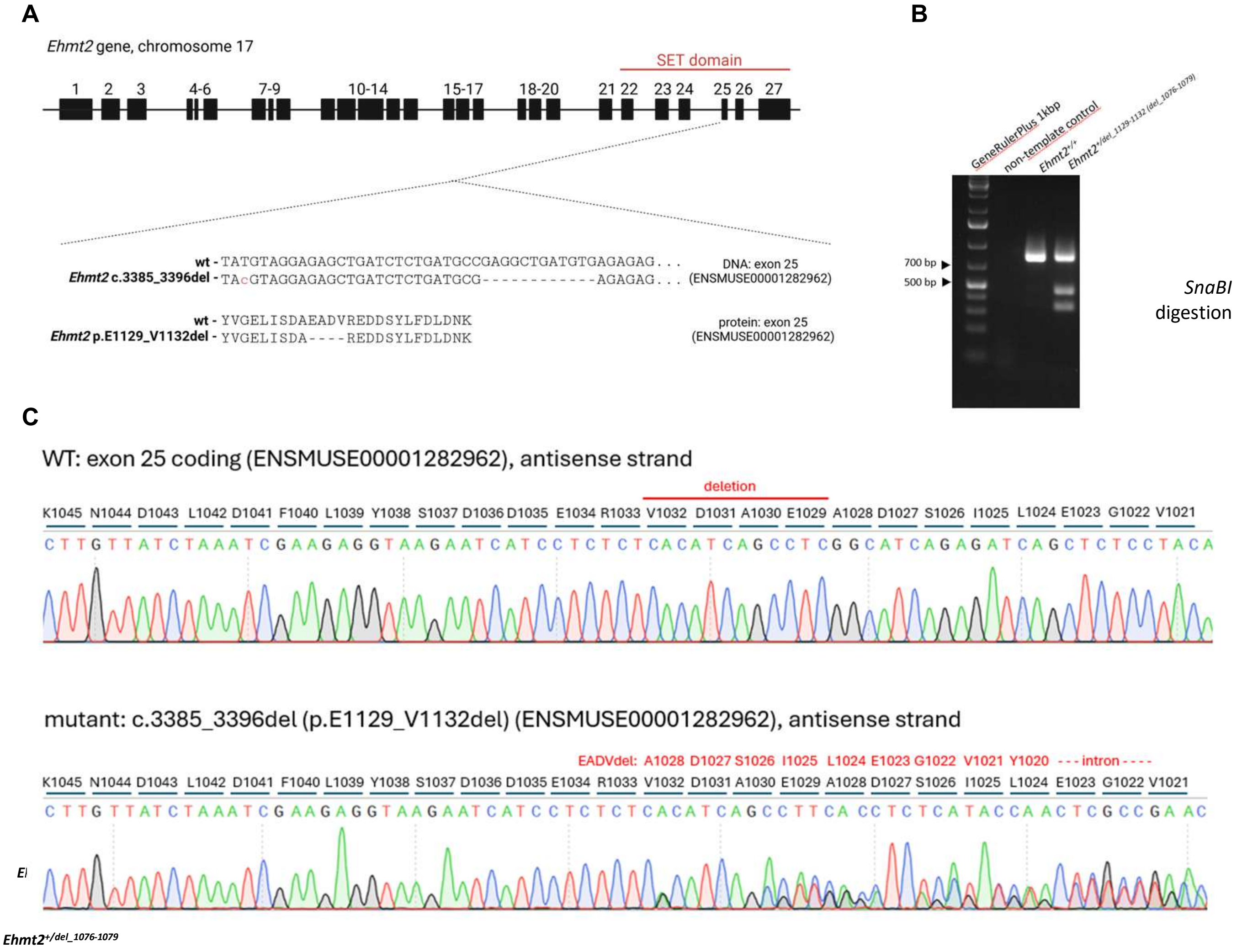


**Supplementary Figure 15**: Generation of a mouse model. A. Localization and sequence characteristics of the *Ehmt2* c.3385_G3396del (p.E1129_V1132del, [C57BL/6NCrl-Ehmt2$^{em1Ccpcz}$/Ph) variant that was designed to mimic the c.3225_3236del (p.Glu1076_Val1079del) *EHMT2* variant detected in patient 1.
B. Genotyping strategy using *SnaBI* digestion.
C. Example of direct Sanger sequencing of the critical targeted *Ehmt2* exon 25 in an *Ehmt2*$^{+/+}$ and *Ehmt2*$^{+/del\_1076\text{-}1079}$ mouse. Frame-shift triggered by the deletion of the 12 nucleotides can be seen in the sequence from the *Ehmt2*$^{+/del\_1076\text{-}1079}$ mouse.

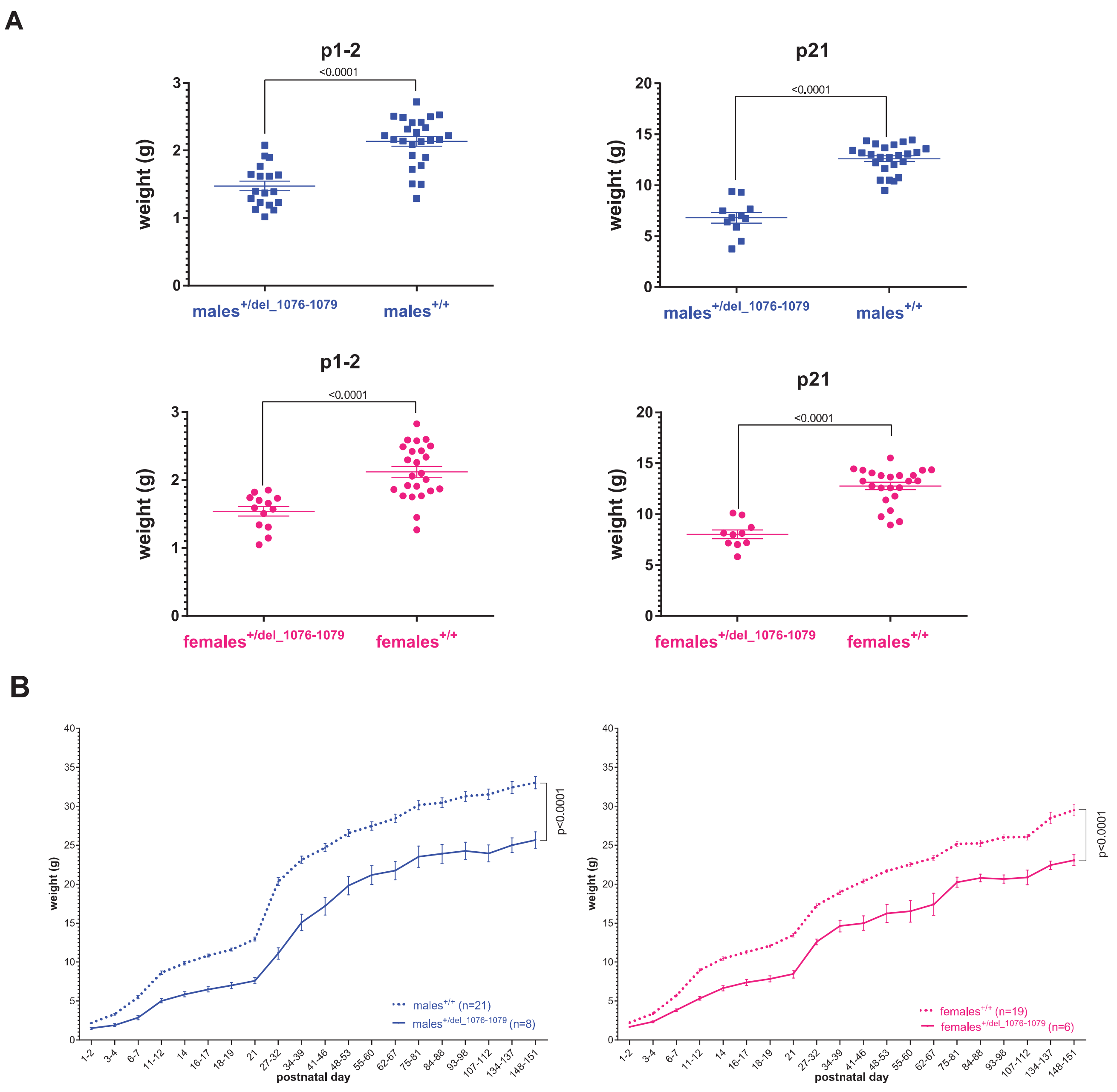


**Supplementary Figure 16**: A. p1-2 and p21 weight data by sex of the mice. *Ehmt2*$^{+/del\_1076\text{-}1079}$ mice have significantly lower body weight regardless of sex at both time points. Statistical significance was calculated using two-tailed Student's t-test.

B. Weight curves of mice surviving the entire 5 month (p151) duration of the study. Compare to weight curve data provided in Figure 6A and 6B. Statistical significance of the differences between the genotypes for each sex of the animals was calculated using two-way repeated measures ANOVA. Average values ±SEM are shown in A and B.

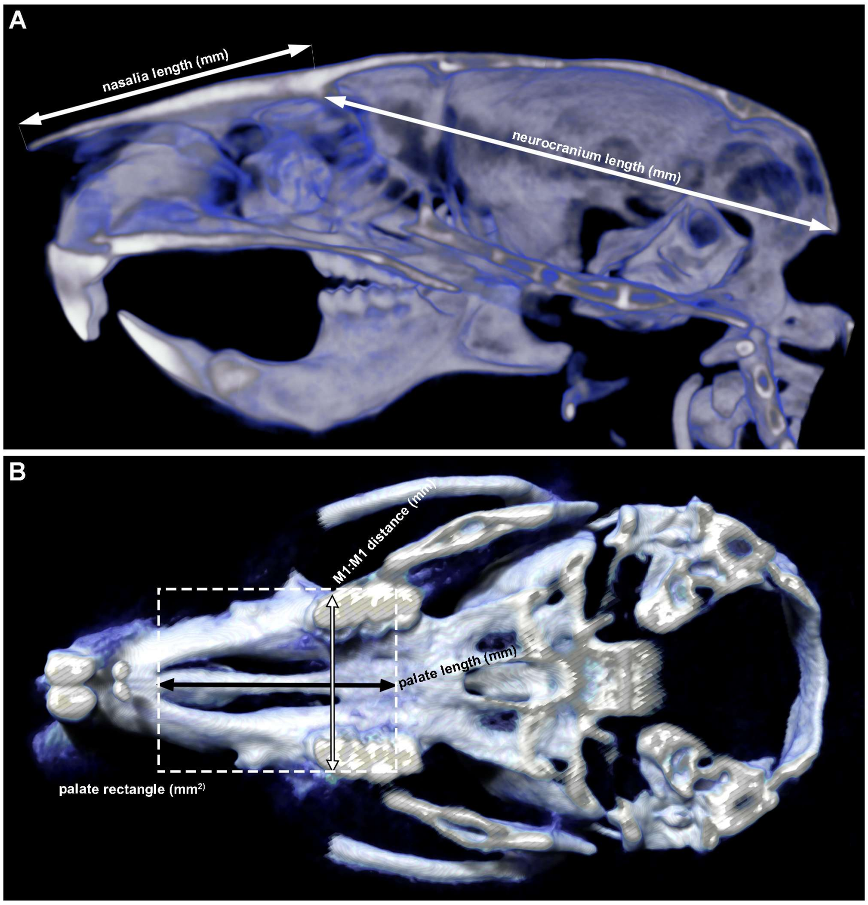


**Supplementary Figure 17**: Craniometric analyses: A. Sagittal skull section. White arrows show the strategy of nasalia and neurocranium length measurements. B. 3D palate reconstruction. Palate length was assessed from the distal rim of the palate arch (fissure) to the suture between maxilla and palate bone. Palate width corresponds to molar 1: molar 1 (M1:M1) distance. Palate rectangle (dashed) was calculated by multiplying the latter two values. See Figure 6D for further details.

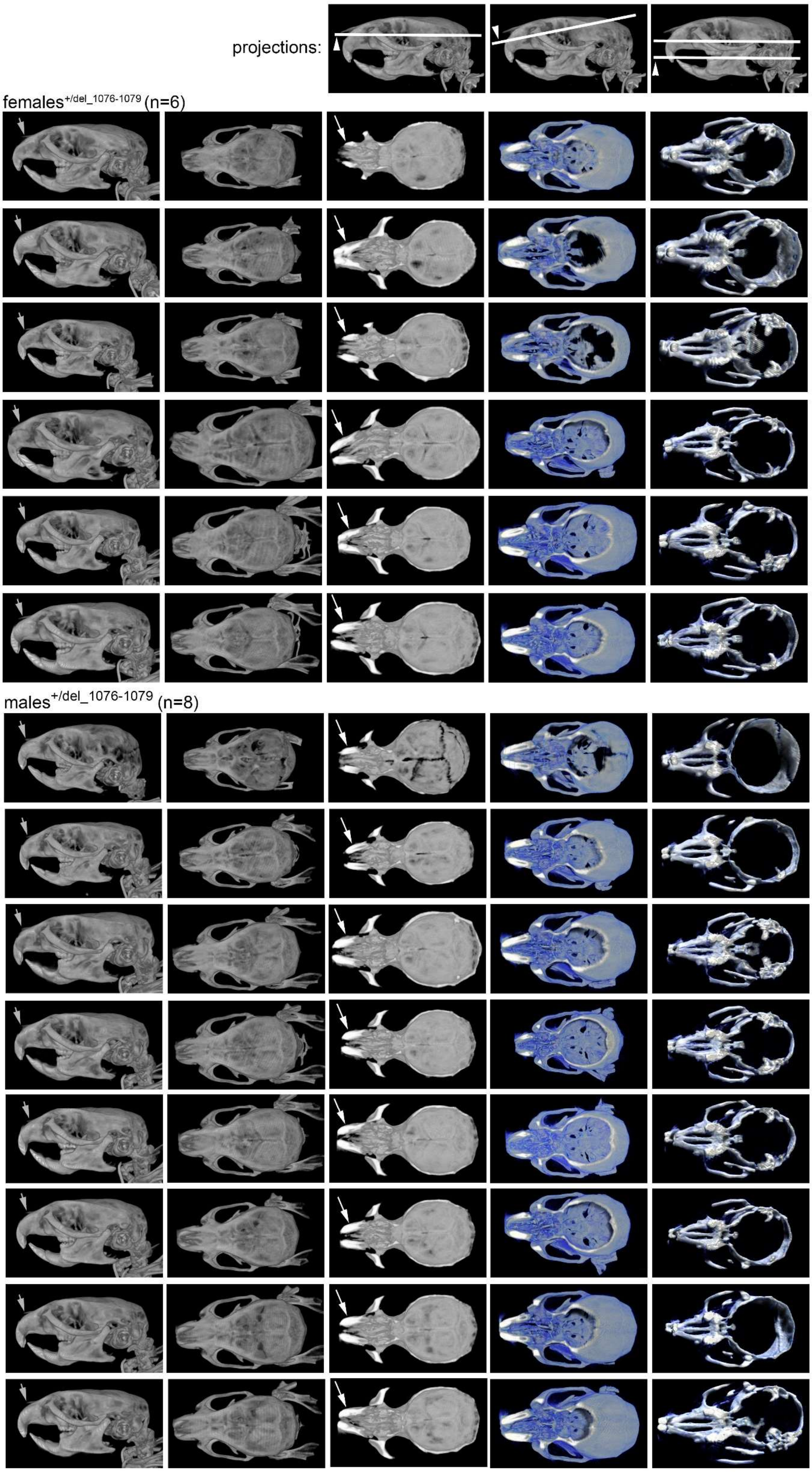


**Supplementary Figure 18**: Overview of microCT data from all tested *Ehmt2*$^{+/del\_1076-1079}$ animals. Left to right. Lateral view at the 3D skull reconstruction. Top view at the 3D skull reconstruction. For the remaining three columns of the projections - white lines show the section plain and arrowheads highlight the view direction (summarized in three images above the layout). Grey arrows in the first column highlight short nasalia, white arrows in the third column highlight the variability of the “bent” nose phenotype in the tested cohort. Compare to Figure 6C and 6D and Supplementary Figure 19.

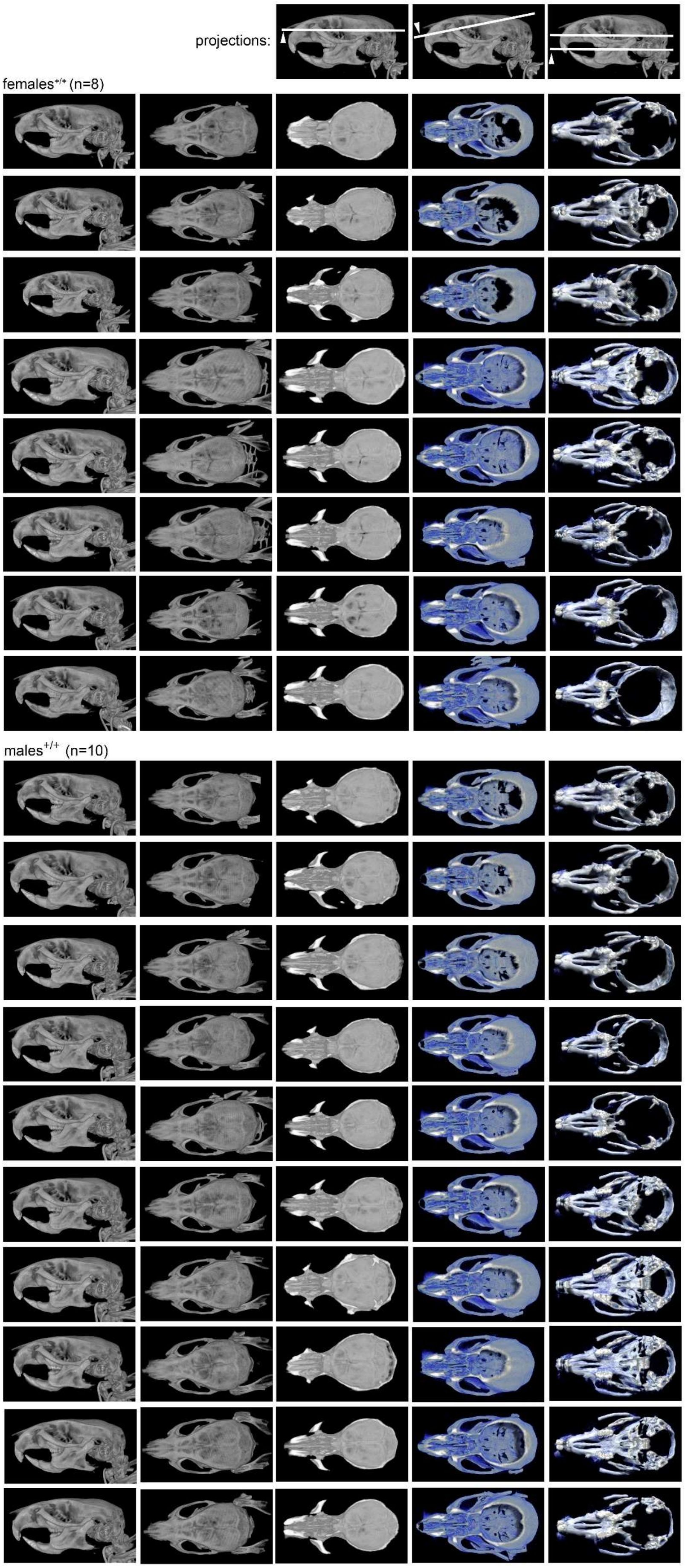


**Supplementary Figure 19**: Overview of microCT data from all tested *Ehmt2*$^{+/+}$ animals. Left to right. Lateral view at the 3D skull reconstruction. Top view at the 3D skull reconstruction. For the remaining three columns of the projections - white lines show the section plain and arrowheads highlight the view direction (summarized in three images above the layout). Compare to Figure 6C and 6D and Supplementary Figure 18.

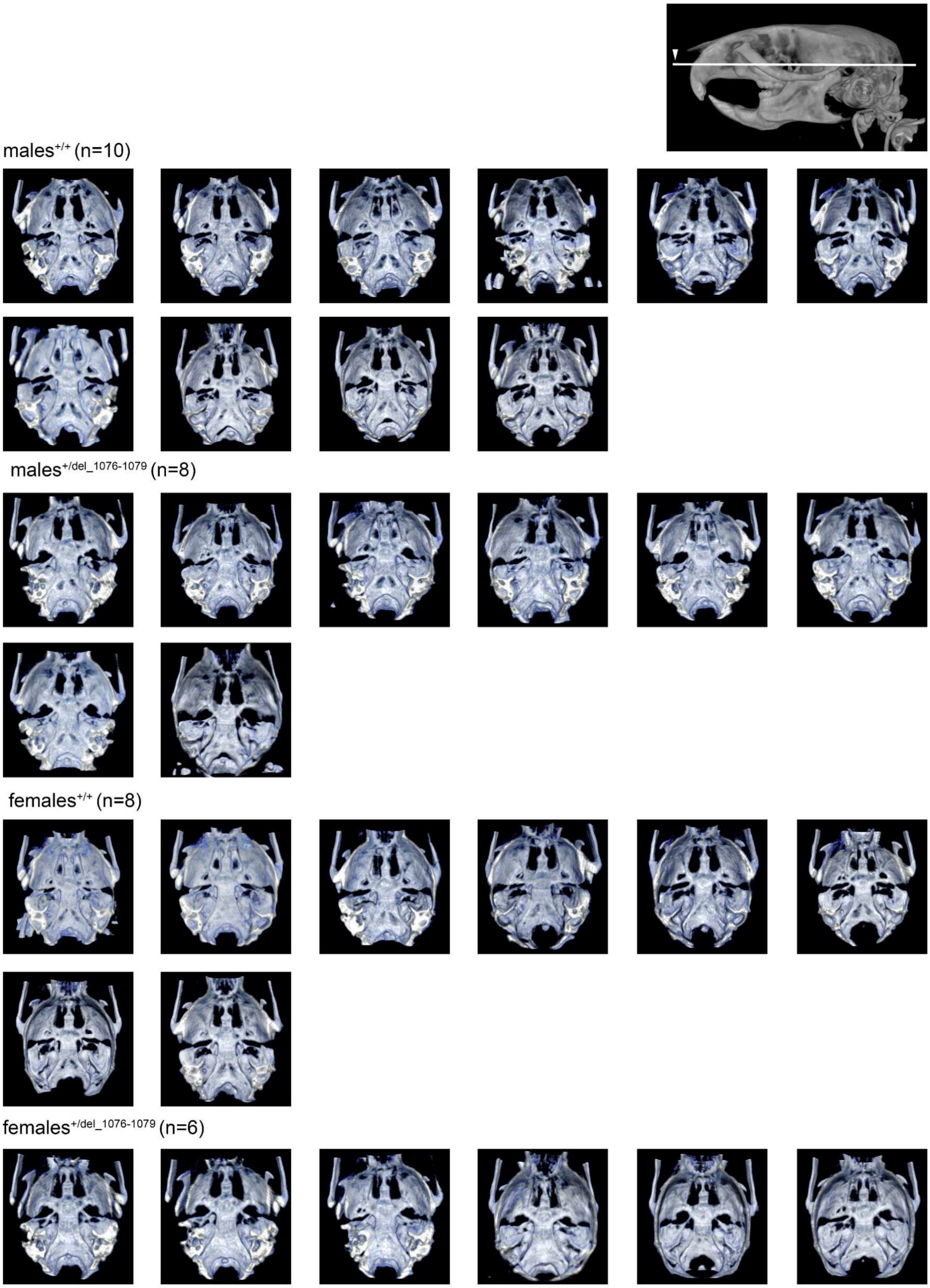


**Supplementary Figure 20**:3D reconstruction of the skull base viewed from the top. Section plane and view direction is shown in the image above the layout. There do not seem to be any abnormalities in sphenoid bone formation in *Ehmt2*$^{+/del\_1076-1079}$ mice in comparison to *Ehmt2*$^{+/+}$ animals.

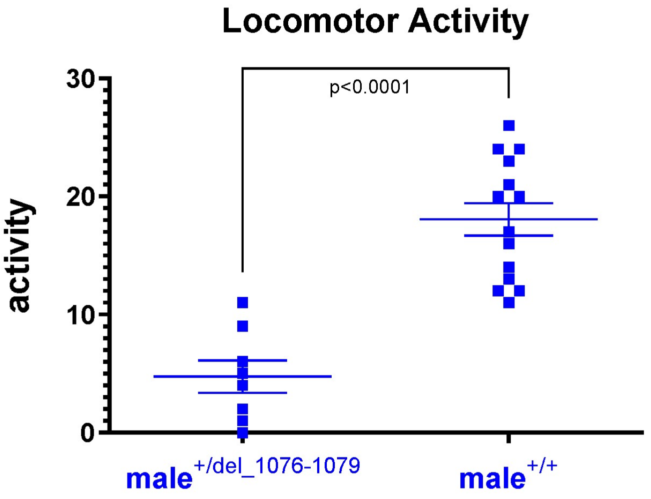


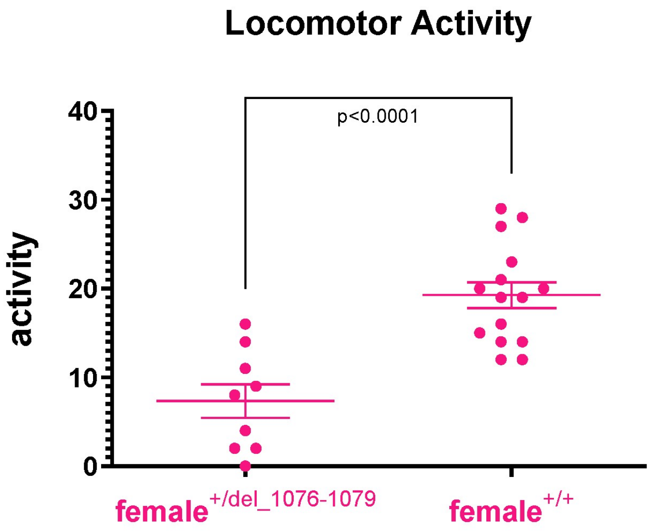


**Supplementary Figure 21**:Differences in the locomotor activity (scored as squares crossed) which is significantly lower in *Ehmt2*$^{+/del\_1076\text{-}1079}$ mice. Statistical significance was calculated using two-tailed Student's t-test. Average values ±SEM are shown.

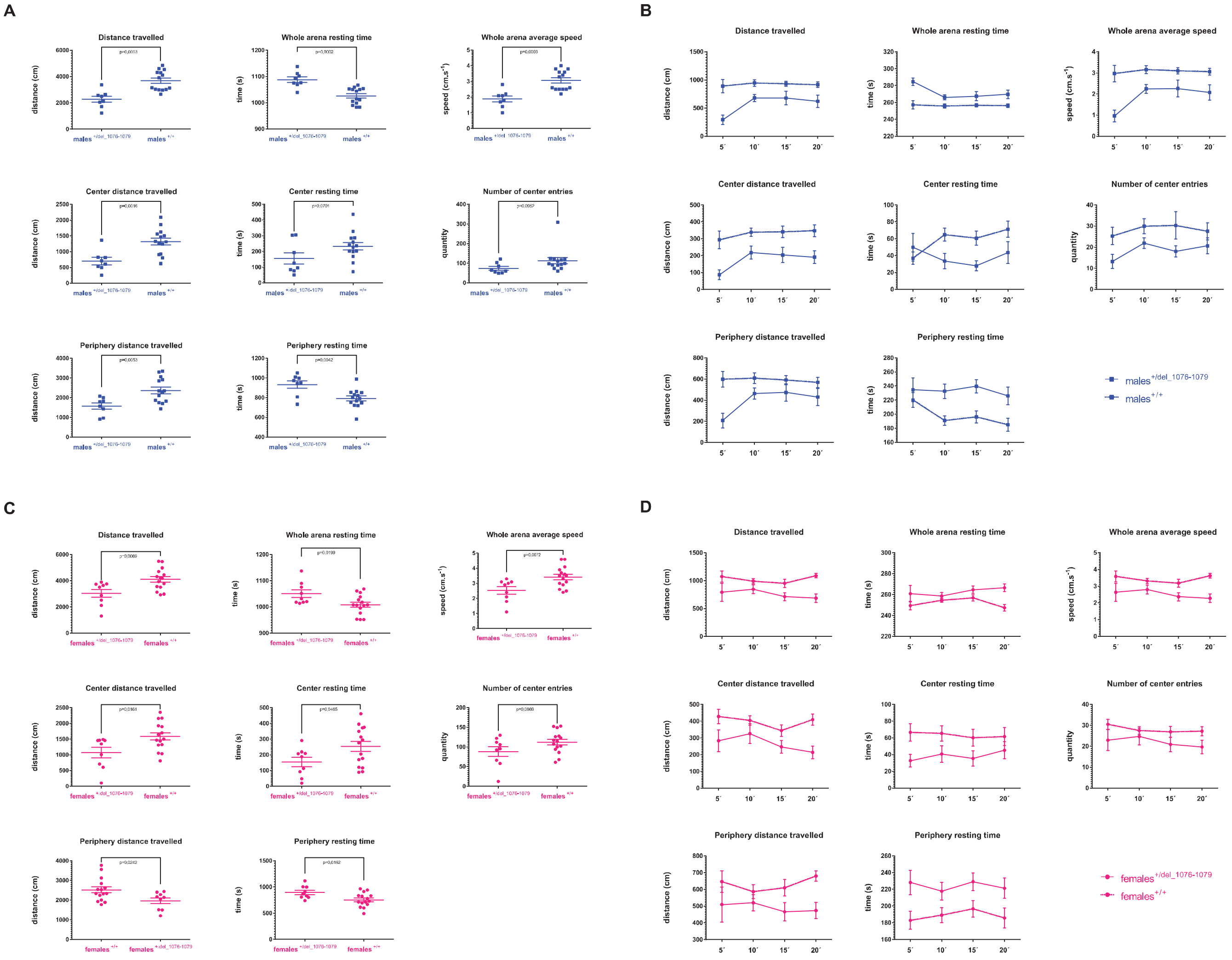


**Supplementary Figure 22**: Summary data (20 minutes) for individual open-field measures are shown for males (A) and females (C). Corresponding interval data (4x5 minutes) are outlined for both sexes (B and D).  Statistical significance was calculated using two-tailed Student's t-test in A and C. Average values ±SEM are shown in A-D.

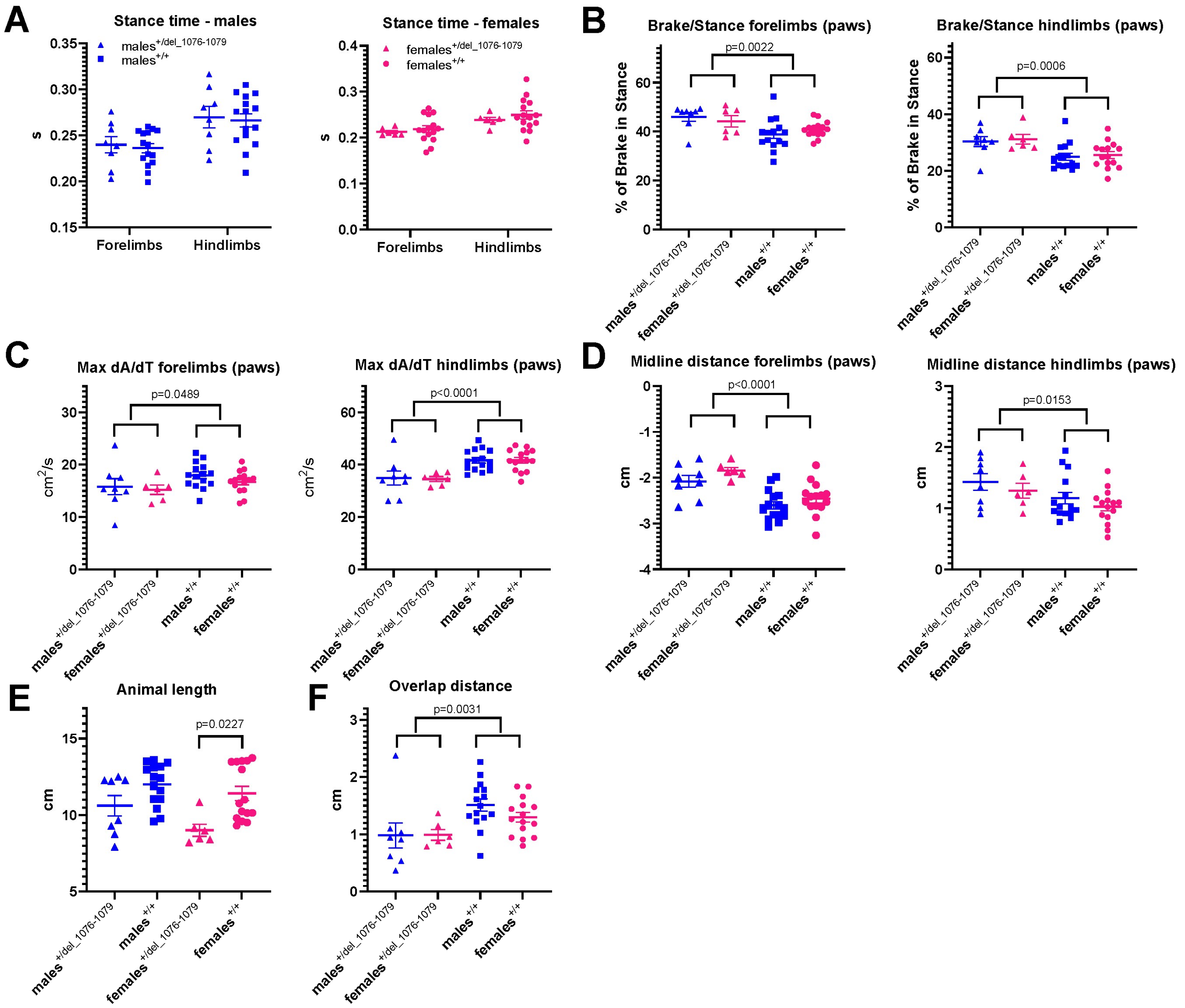


**Supplementary Figure 23**:*Ehmt2*$^{+/del\_1076\text{-}1079}$ mice exhibit significant alterations in gait parameters compared to *Ehmt2*$^{+/+}$ mice.

A. Stance duration (the time during which the paw remains in contact with the treadmill belt) did not differ between the two groups. B. However, the braking phase - „landing“ of the paw to the ground—was significantly prolonged in both the forelimbs and hindlimbs of *Ehmt2*$^{+/del\_1076\text{-}1079}$ mice indicating reduced stability and a need for more precise control of load distribution during stance initiation. C. Difficulties with deceleration and limb loading in *Ehmt2*$^{+/del\_1076\text{-}1079}$ mice were supported by a reduced Maximal rate of change (Max dA/dT) in paw contact area with the treadmill belt during the braking phase, particularly in the hindlimbs. D. Regarding the posture-related parameters, we observed that the midline distance (the mediolateral distance from the body's transverse midline to the paws) was significantly reduced in *Ehmt2*$^{+/del\_1076\text{-}1079}$ mice for the forelimbs (negative midline distance on the *y*-axis is due to the directionality of the measurement) but increased for the hindlimbs. Because midline distance correlates positively with body length, the reduced body length observed in *Ehmt2*$^{+/del\_1076\text{-}1079}$ mice E. may partly explain the decreased forelimb midline distance, particularly in females, but partly could be a result of reduced strength to extend the forelimbs to the extent that the *Ehmt2*$^{+/+}$ mice are capable of. However, this morphological difference cannot account for the increased hindlimb midline distance in the mutant mice, which looks like they stretch their strides too far back or carry the paw further back than normal indicating worsened coordination of gait phases and reduced stability during locomotion. F. Finally, *Ehmt2*$^{+/del\_1076\text{-}1079}$ mice exhibited decreased overlap distance, meaning that hind paws failed to reach the forepaw placement, resulting in a larger gap and indicating impaired inter-limb coordination. Statistical significance was calculated using two-tailed Student's t-test. Average values ±SEM are shown.

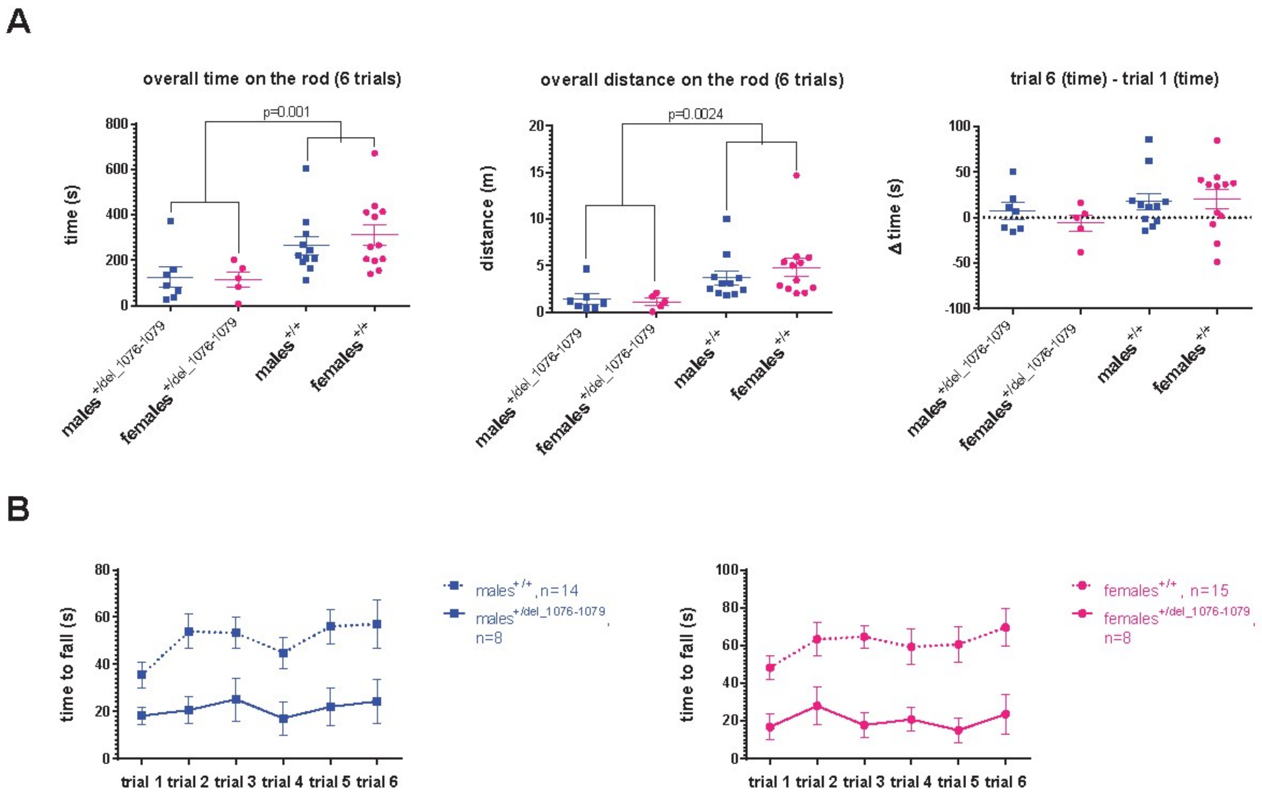


**Supplementary Figure 24**: A. Summary data (6 trials) for time on the rod and distance travelled on the rod are shown. The difference in time-to-fall in trial 6 and trial 1 (Δ time) is not significantly different between *Ehmt2*$^{+/del\_1076\text{-}1079}$ and *Ehmt2*$^{+/+}$ mice. B. Individual interval data for times-to-fall are shown for comparison to overall data.

Statistical significance was calculated using two-tailed Student's t-test in A. Average values ±SEM are shown in A and B.

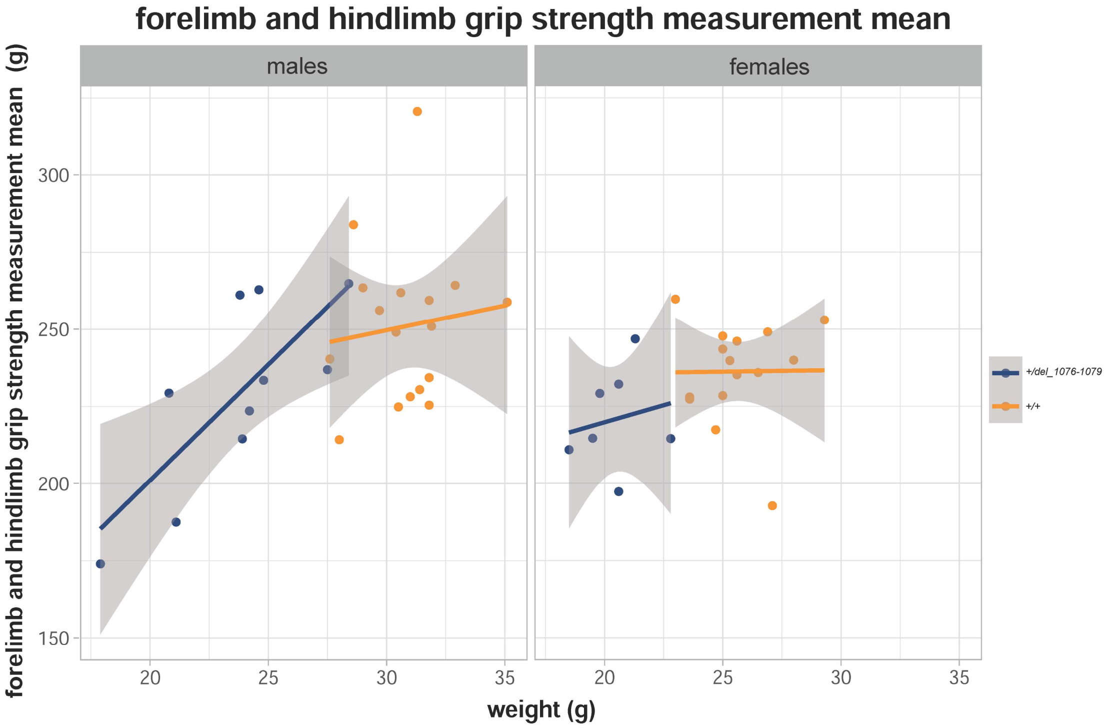


**Supplementary Figure 25**: Grip strength data were analyzed as follows. Triplicate measurements for each mouse were averaged, and a linear mixed-effects model was applied to assess group differences. Genotype, sex, and their interaction were included as fixed effects, with body weight as a covariate. The date of measurement was included as a random effect to account for variability between sessions. To evaluate group differences, contrasts were specified to compare mutant (females and males combined) versus wild-type animals (also combined), as well as within each sex separately. Additionally, the data were visualized with body weight on the x-axis and grip strength on the y-axis, plotted separately for each sex to match the structure of the statistical model.

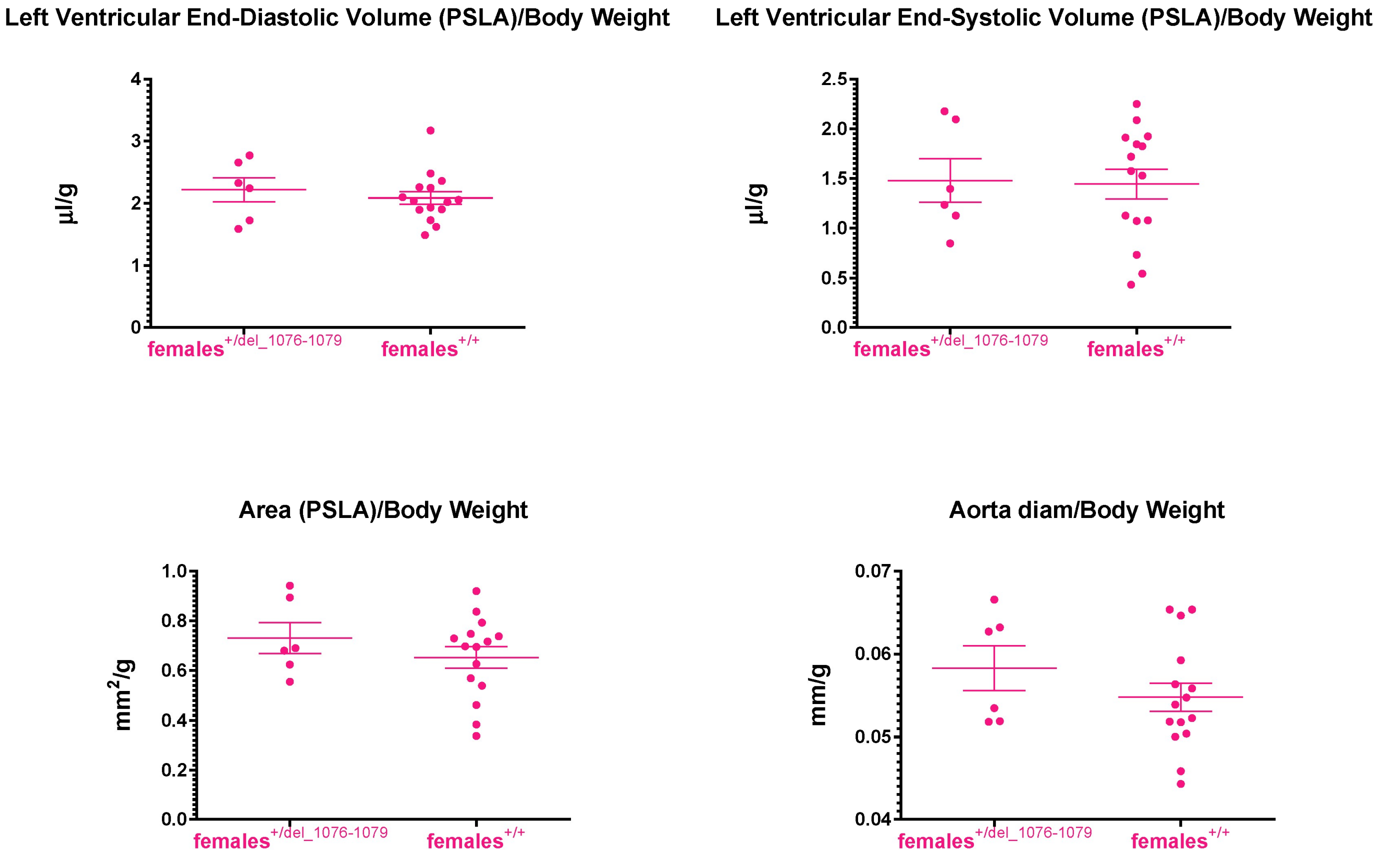


**Supplementary Figure 26**: Left ventricular morphology in female Ehmt2$^{+/del_1076-1079}$ mice assessed by echocardiography

Transthoracic echocardiography revealed that female Ehmt2$^{+/del_1076-1079}$ mice did not exhibit any significant alterations in left ventricular morphology compared to Ehmt2$^{+/+}$ controls. Left ventricular end-diastolic and end-systolic volumes normalized to body weight were comparable between groups, with no significant differences detected. Similarly, no differences were observed in the parasternal long-axis (PSLA) area normalized to body weight or in the aortic diameter normalized to body weight. Statistical analysis was performed using a two-tailed Student's t-test; none of the comparisons reached statistical significance (significance threshold: $p < 0.05$).

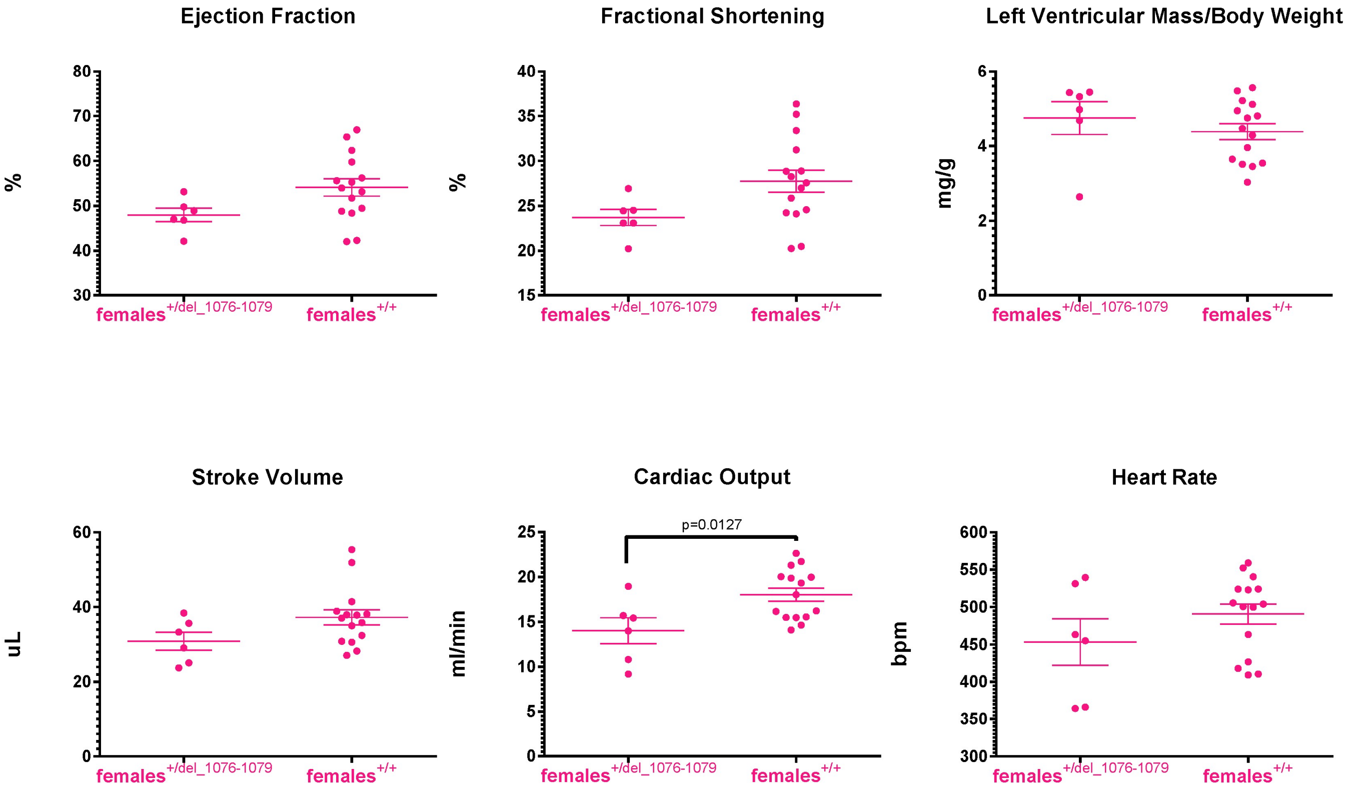


**Supplementary Figure 27**: Cardiac functional parameters in female $Ehmt2^{+/del\_1076-1079}$ mice assessed by echocardiography

Transthoracic echocardiography showed that female $Ehmt2^{+/del\_1076-1079}$ mice did not show significant impairment in functional cardiac parameters compared to $Ehmt2^{+/+}$ controls. Although a mild downward trend in ejection fraction and fractional shortening was observed in mutant animals, these differences did not reach statistical significance. Left ventricular mass normalized to body weight and stroke volume were also comparable between groups. Cardiac output was significantly reduced in $Ehmt2^{+/del\_1076-1079}$ mice; however, this decrease was attributable to a lower heart rate rather than impaired ventricular performance. Overall, echocardiographic findings indicate preserved cardiac structure and function in female $Ehmt2^{+/del\_1076-1079}$ mice under basal conditions. Statistical analysis was performed using a two-tailed Student's t-test. Statistical significance was defined as $p < 0.05$.

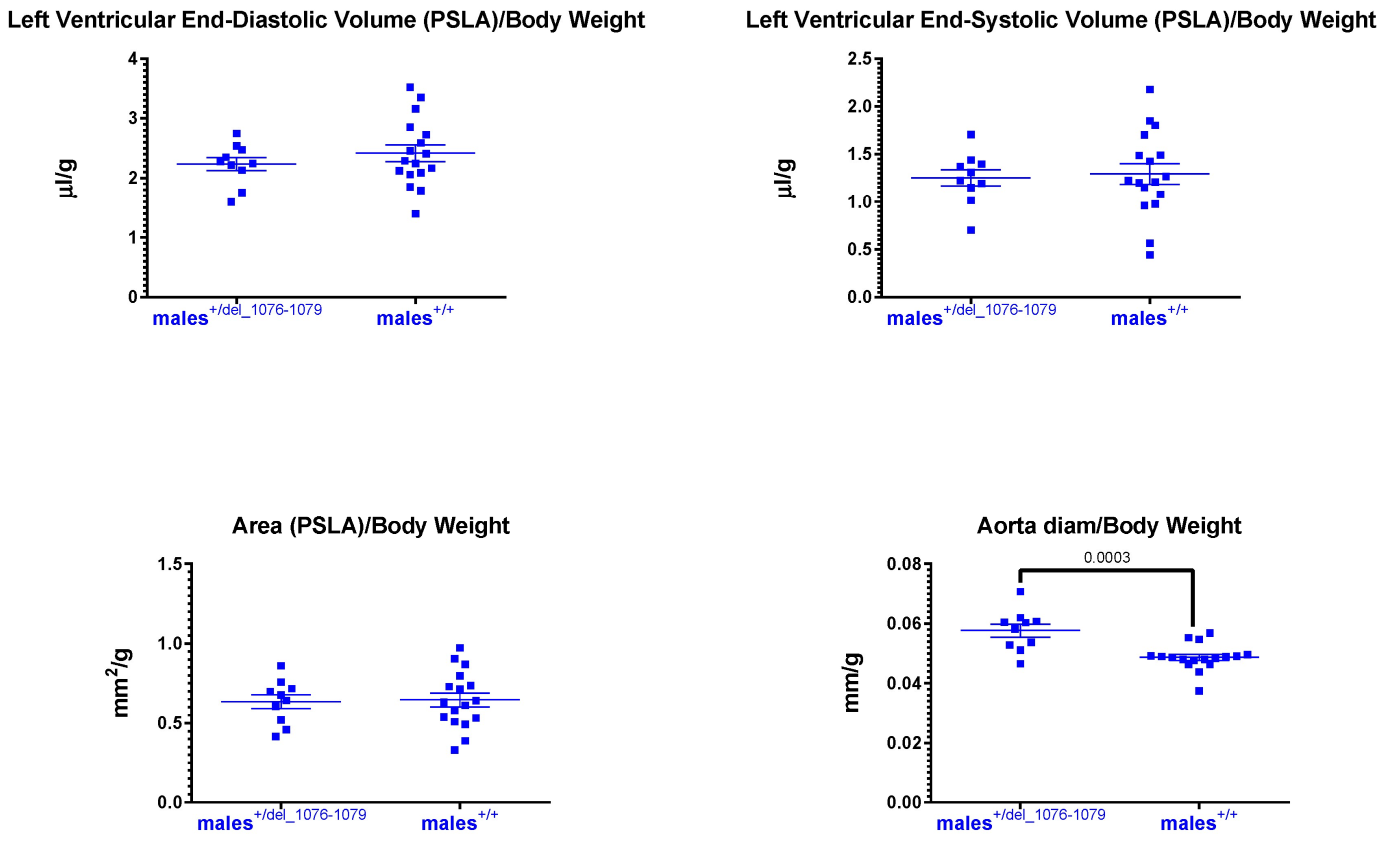


**Supplementary Figure 28**: Left ventricular morphology in male Ehmt2$^{+/del_1076\text{-}1079}$ mice assessed by echocardiography

Transthoracic echocardiography revealed that male Ehmt2$^{+/del_1076\text{-}1079}$ mice did not exhibit significant alterations in left ventricular morphology compared with Ehmt2$^{+/+}$ controls. Left ventricular end-diastolic and end-systolic volumes normalized to body weight were comparable between groups, with no significant differences detected. Similarly, no differences were observed in the parasternal long-axis (PSLA) area normalized to body weight. In contrast, the aortic diameter normalized to body weight was significantly increased in Ehmt2$^{+/del_1076\text{-}1079}$ males. This alteration was not accompanied by changes in left ventricular morphology. Statistical analysis was performed using a two-tailed Student’s t-test. Statistical significance was defined as $p < 0.05$.

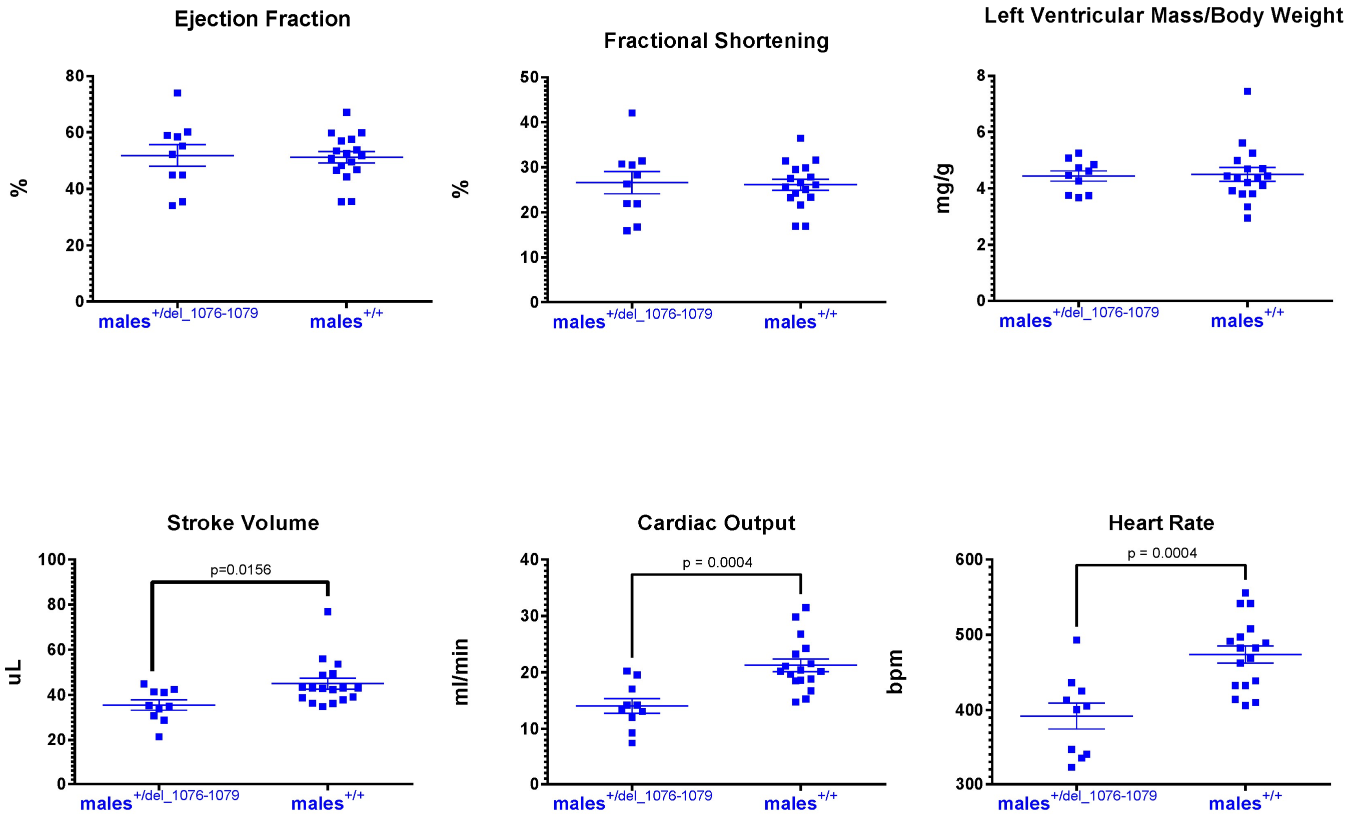


**Supplementary Figure 29**: Cardiac functional parameters in male $Ehmt2^{+/del\_1076-1079}$ mice assessed by echocardiography

Transthoracic echocardiography showed that male $Ehmt2^{+/del\_1076-1079}$ mice exhibited no significant differences in ejection fraction or fractional shortening compared with $Ehmt2^{+/+}$ controls. Left ventricular mass normalized to body weight was also unchanged. In contrast, stroke volume was significantly reduced in $Ehmt2^{+/del\_1076-1079}$ mice, accompanied by a significant reduction in cardiac output and heart rate. The reduction in stroke volume occurred in the absence of changes in ejection fraction or left ventricular mass, indicating preserved myocardial contractility. The decrease in cardiac output was primarily driven by the lower heart rate. Statistical analysis was performed using a two-tailed Student’s t-test. Statistical significance was defined as $p < 0.05$.

**A**

Database of Genomic Variants

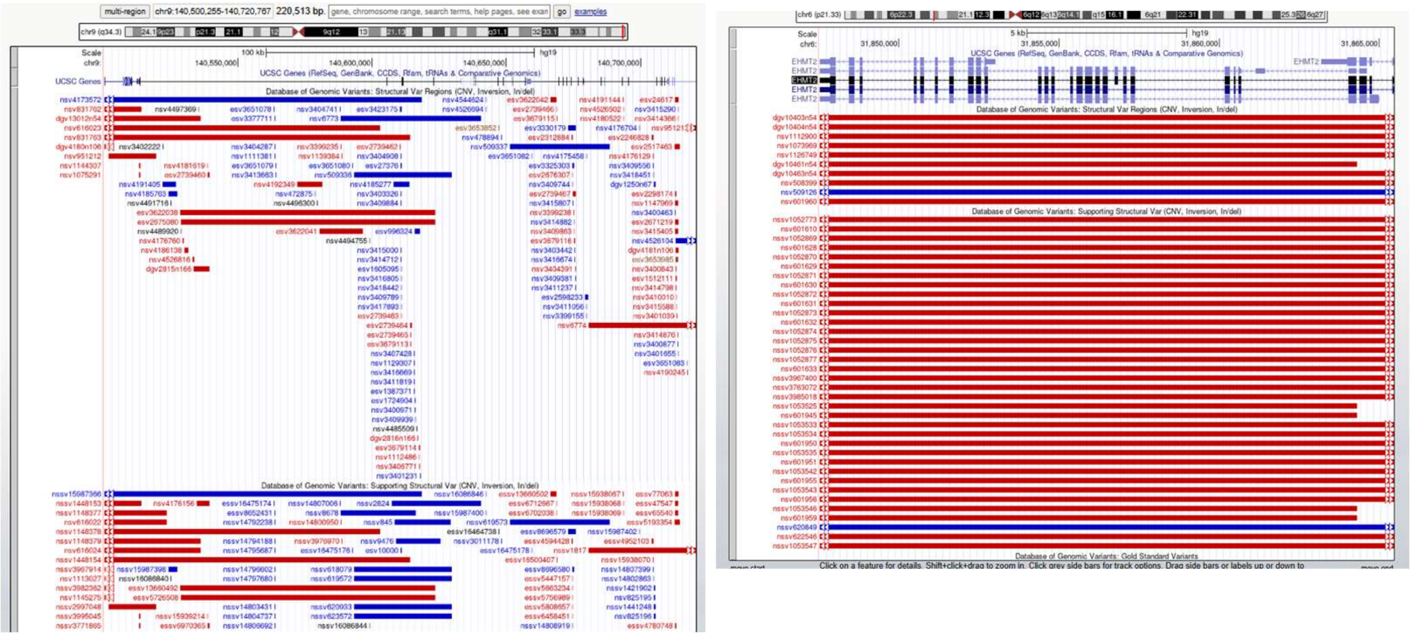

**EHMT1** **EHMT2**

**B**

DECIPHER

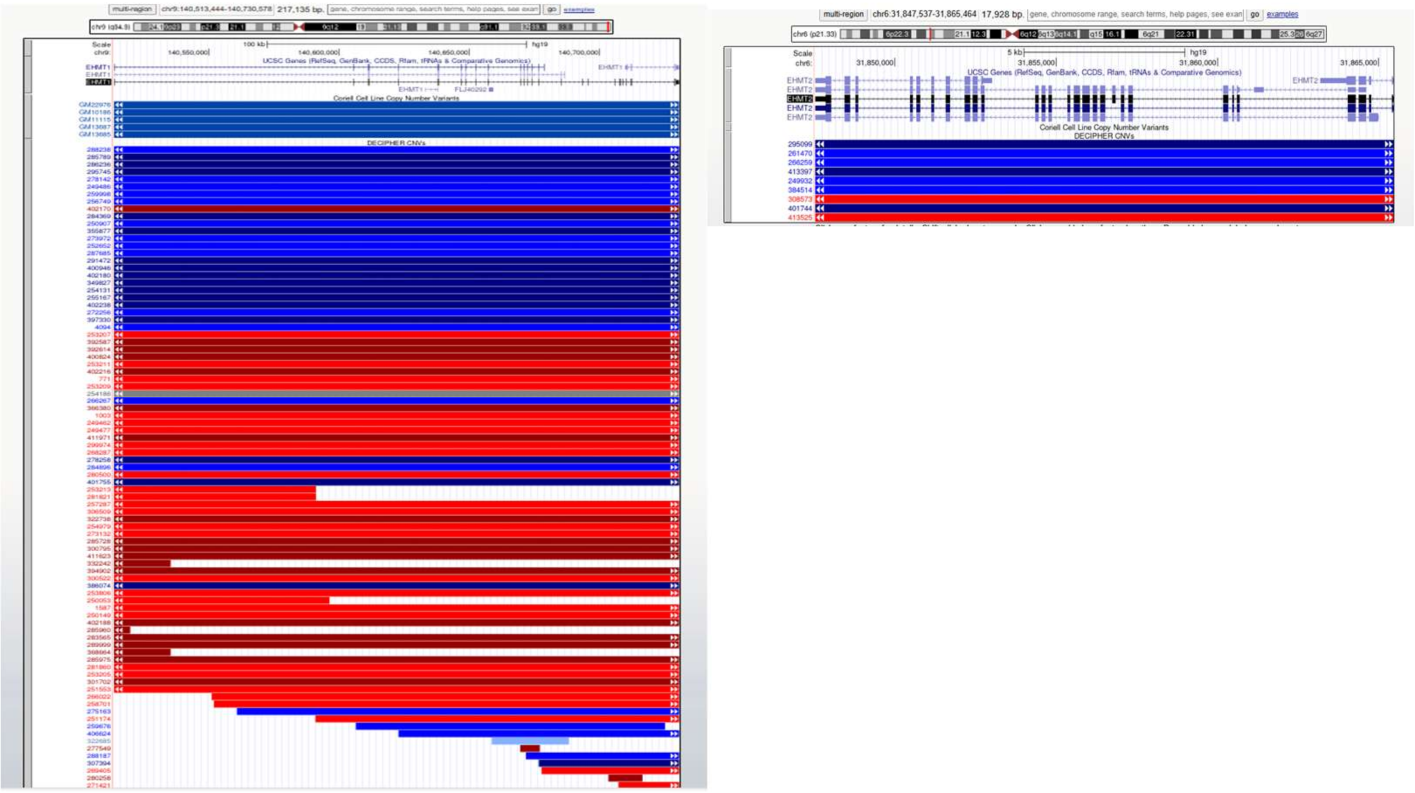

**Supplementary Figure 30.** A. Database of Genomic Variants (DGV) shows sporadic gene deletions in *EHMT1* and frequent deletion events in *EHMT2.*
B. DECIPHER database shows genic deletions in *EHMT1* but not in *EHMT2.*

**Supplementary Video 1:** Representative ventral view video illustrating the altered and unstable gait pattern observed in Ehmt2$^{+/del_1076-1079}$ mutant mice.

**Supplementary Video 2:** Ehmt2$^{+/+}$male animal as a comparison to Ehmt2$^{+/del_1076-1079}$ male in Supplementary video 1. Ventral view video of typical Ehmt2$^{+/+}$ mouse gait pattern.
