## Supplementary Material for "*De novo EHMT2* variants cause an autosomal dominant *EHMT2*-related Kleefstra syndrome via loss of G9a methyltransferase activity"

### *1. Clinical description of the patients with EHMT2 de-novo variants.*

**Proband 1 (P1)** is a girl born at term from the second gravidity, with positive 2nd trimester screening but normal amniocentesis. Birth parameters were normal. The family history is unremarkable. At the age of six months, she underwent surgery for pylorostenosis and was found to be intolerant to cow milk. Stenosis of arteria pulmonalis was found by cardiologic examination. Central hypotonic syndrome was apparent from the age of 6 months, neurological examination ruled out a neuromuscular disorder. The girl shows facial dysmorphism – flat face, hypertelorism, coarse features, high forehead, thin lips, short neck, dysplastic helices; and has unusually big toe. Currently, she suffers from severe developmental delay, cognitive impairment and signs of autism. Emotion-induced stereotypic movements of hands, bruxism and self-mutilation in anger are present in her behavior. EEG showed unspecific bilateral temporal-occipital-parietal focal activity. Central atrophy and atrophy of frontal lobes, enlarged megacisterna magna with arachnoid cyst in posterior fossa, corpus callosum thinning and widening of perioptic CSF spaces were found on MRI. She understands the instructions, but she is unable to speak and is fully dependent on another person care. She is able to walk independently only short distances, while using wheelchair for longer ones. Fine motor skills are suboptimal. She attends a school for children with special needs. Growth and sexual maturation are delayed, with all the growth parameters (height, weight, head circumference) being bellow the third percentile.

**Proband 2 (P2)** was remitted to the undiagnosed rare diseases program SpainUDP after testing negative for candidate gene panels. The proband suffered from global developmental delay characterized by slow development of gross motor milestones and expressive language delay, hypotonia without weakness, coarctation of aorta, nephrocalcinosis and renal medullar cystic disease. The proband had brachycephaly, plagiocephaly with very flat occiput, broad face and midface hypoplasia, synophrys, sparse medial eyebrows, epicanthus, lateral deviation of upslanted palpebral fissures, mildly everted lower eyelids, palpebral ptosis, anteverted nares, smooth philtrum and carp-like mouth. In addition, the proband had dental diastema, microdontia, persistent fetal fingertip pads and scoliosis. From the clinical point of view, the patient's phenotype overlaps significantly with the clinical characteristics of Kleefstra syndrome.

The proband is a male born in 2014 to a 35-year-old mother after a pregnancy complicated by polyhydramnios and intrauterine growth restriction detected at 34 weeks of gestation. The mother was infected by parvovirus B19 during the third trimester of pregnancy, but the newborn's serology was negative for parvovirus (and also for cytomegalovirus). Delivery was induced at 37 weeks of gestation and caesarean section was urgently carried out since a double entanglement and a real knot in the cord were observed. Birth measurements were 2.440 kg of weight (p‹3), 50.5 cm of height (p10-50), a cephalic perimeter of 34.5 cm (p10-50) and an Apgar score of 5/8. The neonate was hospitalized for 15 days after childbirth due to fetal distress assessed during the pregnancy, and clinical evaluation indicated he had jaundice, moderate neonatal depression, membranous septal defect, patent foramen ovale, bilateral cryptorchidism, micropenis, connatal anemia and neonatal hypotonia. At the age of 20 days, he was diagnosed with coarctation of the aorta after an episode of cyanosis and apneas, which required hospitalization and surgical aortic resection and anastomosis. During this hospitalization period, a first electroencephalogram (EEG) showed a pathological profile, although no seizures were observed. However, a second EEG recording revealed no epileptic activity. Neurological examination showed that the baby had dysmorphic facial features and axial hypotonia and therefore he was referred to the neuropediatrician for a follow-up. Three months after birth, he was hospitalized again due to a second episode of cyanosis and apneas, which resolved after non-invasive ventilation. Cerebral ultrasound, video EEG and brain resonance magnetic imaging were performed, but results were normal. Abdominal ultrasound revealed an increased echogenicity of the kidneys and the presence of cysts in the renal medulla, suggesting a renal medullar cystic disease. Anthropometric measurements at that time were 4.590 kg of weight (p‹3), 55 cm of height (p‹3) and a cephalic perimeter of 38 cm (p‹3). At the age of 10 months, his neuropediatrician reported axial hypotonia, unstable sitting, postural plagiocephaly, lower limb hyperreflexia and bilateral inguinal hernia. Weight and height were age appropriate. When reviewed at age 2, he was diagnosed with global developmental delay (mainly characterized by slow development of gross motor milestones and expressive language delay), hypotonia without weakness and nephrocalcinosis. At the age of 3 years, he still showed complete lack of development of speech and certain motor stereotypies. In addition, he underwent orchidopexy for moving his undescended testicles into the scrotum and repairing the associated inguinal hernia (subsequently, this surgical operation has been repeated two more times because it tends to reoccur). At the age of 6 years, he was hospitalized due to an episode of absence seizure, although this type of episodes have not repeated. EEG was normal and brain resonance magnetic imaging only revealed the presence of partially empty sella turcica, which is being followed-up by his endocrinologist. Currently, he has a severe global developmental delay with no speech, although he is able to understand the speech of others and uses a tablet to communicate needs, wishes or thoughts. He shows multiple stereotypic hand movements. His social behavior is normal and he likes establishing social interactions with children and his parents. At the gross motor level, he has unsteady walk with frequent falls, which is complicated by equinovalgus foot deformity and internal rotation and deformity of the left side of his body, where elevated muscle tone and myotatic reflexes are observed. There is no history of ophthalmological or hearing abnormalities. At the craniofacial level, he has brachycephaly, plagiocephaly with very flat occiput, broad face and midface hypoplasia. Hair on the neck extends more inferiorly than usual. He has synophrys, sparse medial eyebrows, epicanthus, lateral deviation of upslanted palpebral fissures, mildly everted lower eyelids, palpebral ptosis (right eye), anteverted nares, smooth philtrum and carp-like mouth. In addition, he has dental diastema, microdontia, persistent fetal fingertip pads and scoliosis. His current anthropometric measurements are 29 kg of weight (p24, -0.71 SD) and 140 cm of height (p68, 0.47 SD). Prior to his entry in the Spanish Undiagnosed Rare Diseases Program SpainUDP, a Prader-Willi test, an array-CGH (750 k) and a clinical exome panel for neurodevelopmental disorders including 6,110 genes yielded negative results. At the age of 4.8 years, he was admitted into the SpainUDP program and the standard criteria for diagnosis established by this program was performed. Deep phenotyping suggested Kleefstra syndrome as a possible diagnosis.

**Proband 3 (P3)** is the first child of non-consanguineous Portuguese parents without any medical history. Pregnancy was marked by discovery of congenital cardiopathy (intra ventricular and auricular communication). She was born at term (38 weeks of gestation) with normal parameters: weight was 2824 g (30th p), length was 49 cm (60th p) and occipito-frontal circonference (OFC) was 34 cm (57th p). Neonatal period was marked by acute renal failure with favorable outcome. Ultrasound showed renal asymmetry with an atrophic and cystic left kidney and a right kidney with moderate hyperechogenicity. She also had a hiatal hernia without any complications. Her psychomotor development was normal with independent walk at 11 months and first words at 8 months. She benefited of atrioventricular canal surgery at 4 years old without any complications. At the age of 6.5 years, she presented with suddenly impaired right upper limb function followed by right lower limb function. Consequently, she suffered with numerous falls. Furthermore, she developed progressive dysarthria, paroxysmal movements (hyperkinesia and choreoathetosis of upper limbs) and perioral dyskinesia. Brain MRI was normal as well as etiological assessment (vascular, infectious, inflammatory, autoimmune). At examination, we noticed some dysmorphic features, i.e. synophrys, long eyelashes, hypertelorism. She was treated with L-DOPA with gradual improvement of symptoms. At the age of 7 years (last examination), she doesn`t present with developmental delay or autistic signs. Her cognitive, motor and verbal abilities are normal.

**Proband 4 (P4)** is a girl born at 39$^{th}$ gestational weeks. Pregnancy was normal and family history was unremarkable. At birth, unilateral postaxial polydactyly of the left foot was noticed. There was facial dysmorphism with frontal bossing, brachycephaly, large anterior fontanelle, short philtrum, and dysplastic cup-shaped and low-set ears. There was a mild pectus excavatum and tapered fingers. There was developmental delay with speech delay, and behavioral anomalies with self-mutilation and aggressivity episodes. The girl walked at age two years. Cardiac examination showed an atrial septal defect.  Brain MRI showed prominent folial pattern in the cerebellar vermis. Evolution showed microcephaly (OFC < - 2 SD), moderate deafness, and absent speech.

**Proband 5 (P5)**, a male, is the second child born to healthy unrelated parents. The pregnancy resulted from *in vitro* fertilization, marked by hypoplasia of nasal bones and single umbilical artery at ultrasound and, maternal diabetes during the 3$^{rd}$ trimester. He was born at 41+6 weeks gestation by vaginal delivery with normal birth parameters and an Apgar of 10/10. He presented feeding difficulties in the neonatal period, ventricular muscular septal defect that required surgery at 7 months of age. Posterior urethral valves were surgically repaired at at 20 months of age after an episode of pyelonephritis and febrile seizure. Motor development was delayed as he sat unaided after 9 months and walked unaided at 24 months. He spoke meaningfull words, associated words into sentences and had good social interactions at the age of 2 years. He could swallow and had non-severe chronic constipation. Growth was in the normal range. Facial features involved synophrys, microstomia and crumpled ears. He also had stenosis of one lacrimal duct and frequent mid-ear infections.

**Proband 6 (P6)** is the second child of healthy, unrelated, white Scottish parents. Antenatal ultrasounds demonstrated a small left ventricle, dilated stomach and duodenum and polyhydramnios. The child was delivered at 39+2 weeks gestation by elective caesarean section. She was born in good condition, with Apgar scores 9 at 1 minute and 9 at 5 minutes. Birthweight was 2840g. Abdomen was distended but soft after birth, with Xray showing distended, gaseous bowel loops. Nonetheless, she passed meconium shortly after birth and a suction rectal biopsy found no evidence of Hirschprung disease. Echocardiogram showed a hypoplastic aortic arch, dysplastic pulmonary valve with mild narrowing and an acute angle turn of both branch pulmonary arteries. At 8 weeks of life, coarctation of the aortic isthmus was repaired by resection and end-to-end anastomosis. The child went on to have significant developmental delay. MRI of brain was reported as unremarkable, except for mild asymmetry of the ventricles and prominence of the extra-axial spaces. EEG at 2 years 7 months revealed abnormal background rhythm ("persistent rhythm 4 Hz theta activity over the temporal lobes. An 8 Hz central mu rhythm was seen while brief passive eye closure evoked a 3.5 Hz posterior rhythm"). Extensive metabolic biochemical investigations were unrevealing.

Noisy breathing was noted in early life and sleep studies were abnormal, with clustered desaturations, which persisted despite adenotonsillectomy. She commenced nocturnal BiPAP therapy from around 2.5 years of age. Recurrent abdominal distension has also been noted. Upper gastrointestinal endoscopy and sigmoidoscopy revealed normal mucosa, but evidence of faecal impaction. Symptoms have persisted despite aperients. On most recent review at 3 years 10 months, the child can sit independently and stand with support. She has not spoken any words, but vocalises readily and parents feel these are often meaningful in terms of tone. She can express happiness and enjoys social interactions. There are tongue-clicking stereotypies and stimming behaviors. On examination, she had a brachycephalic head shape with midface retrusion, flat facial profile and prognathism. Head circumference measured 49.2cm (48th centile). Corners of the mouth were downturned, and primary dentition was not fully erupted. Her sclerae appear grey. There was generalised cutis marmorata, and follicular hyperkeratosis affecting the upper arms.

### *2. Methods of identification of EHMT2 variants*

**P1** was ascertained from a series of patients with paediatric-onset rare diseases with unknown genetic basis who underwent whole-exome sequencing. Informed consent for genetic analyses was obtained for all individuals, and genetic studies were performed as approved by the Institutional Review Board of the First Faculty of Medicine of the Charles University, Prague, the Czech Republic. The patient's parents provided written informed consent for the participation in the study, clinical data and specimen collection, genetic analysis and publication of relevant findings.

Genomic DNA extracted from leukocytes of patient and her parents was used for whole-exome sequencing. Exome enrichment was performed on individually barcoded samples using SeqCap EZ MedExome Probes (Roche) and sequencing was performed on Novaseq 6000 platform (Illumina) with 100bp paired-end reads. Reads were aligned to the hg19 reference genome using Novoalign version 3.02.13 (Novocraft) with default parameters.

After genome alignment, conversion of SAM format to BAM and duplicate removal was performed using Picard Tools (2.20.8). The Genome Analysis Toolkit, GATK (3.8)19 was used for local realignment around indels, base recalibration, variant recalibration, and variant calling. Variants were annotated using the GEMINI framework20 and filtered based on the population frequencies using several public databases and an in-house database of population-specific variants. Identification of candidate variants was performed for autosomal dominant (de novo variants) and autosomal recessive inheritance patterns. Variants were further prioritized according to the functional impact and conservation score.

Sanger sequencing was used for segregation studies to confirm the variant status in the whole family.

**P2** was recruited by the undiagnosed rare diseases program SpainUDP[1] at the Institute of Rare Diseases Research (IIER), Spanish National Institute of Health Carlos III (ISCIII). Peripheral blood samples and skin biopsies were collected from patient and his parents to perform trio-based whole-exome sequencing and to establish fibroblast cultures. Informed consents were signed by the patient's legal representatives. This research project was approved by the ISCIII Research Ethics Committee.

Whole exome sequencing and data analyses were performed in the proband and his unaffected parents. Genomic DNA was extracted from peripheral blood using the Qiagen QIAamp DNA kit. Whole Exome Sequencing (WES) libraries were prepared using the Nimblegen MedExome + ChrMit as enrichment kit and HS2000 v4, 2×100bp sequencing platform in ND095 family and Nextera Felx DNA Library Prep and Illumina NextSeq500 in ND120 family. Data analysis was performed using two different standardized protocols as previously described[1]. These included an in-house analysis and a parallel analysis using the Genomic Analysis module of the RD-Connect Genome-Phenome Analysis Platform (GPAP)13. All identified rare variants were further analysed, checking all available scientific evidence through detailed searches in public databases (including Gene-Card, NCBI, UniProt, OMIM, Pubmed and ExAc). Sanger sequencing was performed to validate candidate variants.

**P3** underwent trio exome sequencing. Libraries of genomic DNA samples were prepared using the Twist Human Core Exome kit (Twist Biosciences, San Francisco, CA), and were sequenced on a NovaSeq 6000 instrument (Illumina, San Diego, CA) according to the manufacturer's recommendations for paired-end 101-bp reads. A mean depth of 93.34 x was reach and 97.1 % of the refseq exons were covered at least by 10 reads.

Variants were identified using a computational platform of the FHU Translad, hosted by the University of Burgundy Computing Cluster (CCuB). Raw data quality was evaluated by FastQC software (v0.11.4). Reads were aligned to the GRCh37/hg19 human genome reference sequence using the Burrows-Wheeler Aligner (v0.7.15). Aligned read data underwent the following processing steps: (a) duplicate paired-end reads were removed by Picard software (v2.4.1), and (b) base quality score recalibration was done by the Genome Analysis Toolkit (GATK v3.8) Base recalibrator. Using GATK Haplotype Caller, Single Nucleotide Variants with a quality score >30 and an alignment quality score >20 were annotated with SNPEff (v4.3t). Rare variants were identified by focusing on nonsynonymous changes present at a frequency less than 1% in the gnomAD database. Copy Number Variants were detected using xHMM (v1.0) and were annotated using in-house python scripts. They were filtered based on their frequency in public databases (DGV, ISCA, DDD).

**P4** was recruited by the undiagnosed rare diseases program Solve-RD from ERN ITHACA. Chromosome analysis on lymphocytes, array-CGH (Agilent 105 K) and exome sequencing were normal. Genome sequencing revealed a variant in gene *EHMT2* (p.Phe1158Leu).

Whole genome sequencing was performed in **P5** as part of the diagnostic process in France (Plan France Médecine Génomique 2025, PFMG2025) at the genomic laboratory SeqOIA. Genomic DNA was extracted from leukocytes of the patient and his parents using QiaSymphony. Whole genome sequencing libraries were prepared using the mina DNA PCR-free Prep and sequencing was performed on NovaSeq 6000®, Illumina® with 150bp paired-end reads using SBS technology and S4 Flow Cell. Reads were aligned to the GRCh38.92.fa reference genome using bwa-mem2 (v2.2.1). After genome alignment, SNV and indel<50bp were detected using Haplotype caller, GATK (v4.1.9.0). Variants were annotated using the SNPEff (v4.3t) and filtered based on the population frequencies using public databases. Variants were further prioritized according to the inheritance, functional impact and conservation score.

Trio-based whole exome sequencing was performed in **P6** as part of a diagnostic service with the Scottish National Health Service. Libraries were constructed using the Illumina DNA Prep with Enrichment kit and capture was performed using Human Core Exome and Human RefSeq Panel probes from Twist Bioscience. Libraries were sequenced on a NovaSeq 6000 instrument using an S2 flow cell (2x100bp) (Illumina). Initial analysis of the DDG2P gene panel (https://www.ebi.ac.uk/gene2phenotype/panel/DD) using an in-house pipeline revealed no plausible causative variants. Subsequently, prompted by clinical suspicion of Kleefstra syndrome and supportive episignature, targeted analysis was undertaken of EHMT2 using the commercial software platform, Congenica (https://www.congenica.com/). This identified the *de novo* variant (NM_006709.5:c.3485A>G p.(Lys1162Arg)). Confirmatory Sanger sequencing was performed.

1. López-Martín, E. *et al.* SpainUDP: The Spanish Undiagnosed Rare Diseases Program. *Int J Environ Res Public Health* **15**(2018).
