## Supplementary material for "*De novo EHMT2* variants cause an autosomal dominant *EHMT2*-related Kleefstra syndrome via loss of G9a methyltransferase activity": Table 1

| Clinical data of patients with EHMT2 variants. | | | | | | | |
|---|---|---|---|---|---|---|---|
| Study Patient ID | **P1** | **P2** | **P3** | **P4** | **P5** | **P6** | **P7** |
| Inheritance | de novo | de novo | de novo | de novo | de novo | de novo | de novo |
| Zygosity | heterozygous | heterozygous | heterozygous | heterozygous | heterozygous | heterozygous | heterozygous |
| Mutation (hg38) | 6-31881053-CACATCAGCCTCA-C[illegible]12bp) | 6-31881061-C-A | 6-31880790-T-C | 6-31880245-A-G | 6-31880224-ATTTGCT-A | 6-31880232-T-C | 6-31881061-C-T |
| Mutation (NM 006709.5) | c.3225 3236del | c.3229G>T | c.3335A>G | c.3472T>C | c.3487 3492del | c.3485A>G | c.3229G>A |
| Mutation (NP 006700.3) | p.Glu1076 Val1079del | p.Ala1077Ser | p.Asn1112Ser | p.(Phe1158Leu) | p.(Ser1163 Lys1164del) | p.(Lys1162Arg) | p.Ala1077Thr |
| gnomAD V4.1 occurence | 0x | 0x | 0x | 0x | 0x | 0x | 0x |
| AlphaMissense | NA | 0.98 (pathogenic) | 0.94 (pathogenic) | 0.95 (pathogenic) | NA | 0.97 (pathogenic) | 0.99 (pathogenic) |
| REVEL | NA | 0.77 (Pathogenic) | 0.8 (Pathogenic) | 0.81 (Pathogenic) | NA | 0.76 (Pathogenic) | 0.75 (pathogenic) |
| SpliceAI | 0 (no predicted effect on splicing) | 0 (no predicted effect on splicing) | 0.1 (no predicted effect on splicing) | 0 (no predicted effect on splicing) | 0 (no predicted effect on splicing) | 0 (no predicted effect on splicing) | 0 (no predicted effect on splicing) |
| GERP score | 4.59 | 4.59 | 4.35 | 4.75 | 4.75 | 4.75 | 4.59 |
| Gender | female | male | female | female | male | female | male |
| Last follow-up age | 15 years | 10 years | 7 years, 6 months | 16 years | 2 years | 4 years 2months | 3 years 10 months |
| Occipital frontal interference at birth | 34,5 cm | 34,5 cm | 34 cm | 32 cm | 35.5 at 41 WG + 6 days | 33.1 cm | 33 cm |
| Length at birth (cm) | 51 cm | 50,5 cm | 49 cm | 48 cm | 55 cm | not recorded | 47 cm |
| Weight at birth (g) | 3150 g | 2440 g | 2824 g | 2630 g | 3100 g | 2840 g | 2390 g |
| Occipital frontal interference at last evaluation | 51 cm | | 48.5 cm | 51 cm | 48.5 cm | 49.2cm at 3yr 11 mo | ND |
| Height (cm) | 145 cm | 140 cm | 100.8 cm | 163.5 cm | 90 cm | 92.4cm at 3yr 3 mo | 102.2 cm |
| Weight (kg) | 30 kg | 29 kg | 15. 3 kg | 62 kg | 13 kg | 14.05 kg at 3yr 3mo | 14.7 kg |
| **Neurological findings:** | | | | | | | |
| hypotonia | yes | yes | no | yes | yes | yes | yes |
| EEG | Nonspecic bilateral focal activity located temporo - occipito-parietally. | Pathological profile at 15 days, normal in later recordings. | Correctly structured wake and sleep pattern apart from a sometimes slower left hemispherical aspect and acute but subclinical flushes? No focus of electric shock abnormalities. | No epilepsy. | ND | No focal or epileptiform anormalities. | Negative for seizure activity. |
| MRI | Periventricular atrophy, cortical atrophy, megacisterna magna with arachnoid cyst along the left cerebellar hemisphere. | Partially empty sella turcica. | Punctiform FLAIR hypersignal in the left frontal white SB with diffusion restriction, compatible with an ischemic lesion. FLAIR hypersignal from the knee of the corpus callosum on the right, without diffusion restriction, probably a vessel and moderate dilatation of the lateral ventricles. | ND | ND | Mild asymmetry of ventricles and prominence of extra-axial spaces. | Abnormal corpus callosum; mild hypoplasia; mildy enlarged cisterns without Dandy Walker, abnormal subarachnoid space with mildly enlarged bifrontal space, moderate ventriculomegaly without hydrocephalus, and multiple diffuse (mostly subcortical) white matter gliosis. |
| Seizures | No | one episode | No | No | Febrile seizure at 20 months. | No | Febrile seizures at 1y 10mo of age. Seizure-like activity vs sensory seeking behaviors (had twitching, arches head backwards and forewards, extreme belly laughts without provocation - all followed by period of prolonged sleep); unremarkable EEGs. |
| **Psycho-developmental and behavioural findings:** | | | | | | | |
| Development | severe developmental delay | psychomotor delay | normal | moderate developmental delay | motor delay | global developmental delay | psychomotor delay |
| Autism | some autistic signs | normal social behavior | no | no | no | no, sociable and enjoys company | ND |
| Behavior | hyperkinetic movements of hands in emotions, bruxism, self-mutilation in anger | stereotypic hand movements | sudden onset of hyperkinetic movement and some choreoathetosic movements, improvement with L-dopa | behavioral disorder | | stereotypic hand and limb movements, tongue-clicking | atypical hand movements |
| ID | severe mental retardation | severe global developmental delay/intellectual disability | no | moderate developmental delay | | moderate-severe global delay (Non-verbal and non-ambulant at age 4 years). | significant global developmental delay/ skills 4-8 months |
| Motor abilities | walks independently - short distances, longer distances on wheelchair, suboptimal fine motor skills | unsteady walk with frequent falls | normal | walks independently | walked unaided at 24m | at age 4 years, sits independently and stands with support. Not independently ambulant. | significant motor and adaptive functioning delays |
| Cognitive abilities | understands the instructions | understands the speech | normal, no learning difficulties | understands the instructions | | demonstrates receptive understanding, expresses happiness and enjoyment appropriately. | significantly delayed |
| Verbal abilities | absent speech | no speech | complexe sentences | limited language with simple sentences | a few words | absent speech. Some vocalisations are meaningful in terms of tone. | no speech |
| **Cranial, facial and somatic dysmorphisms:** | | | | | | | |
| Truncal obesity | – | – | – | + | – | – | – |
| Short neck | + | – | – | – | – | – | – |
| Brachycephaly | + | + | – | + | – | + | + |
| Plagiocephaly with flat occipit | + | + | – | + | – | + | + |
| Flat face | + | – | – | – | – | + | – |
| Broad face | + | + | + | + | – | + | + |
| High forehead | + | – | – | – | – | + | – |
| Midface hypoplasia | + | + | + | + | – | + | + |
| Prognathia/midface retraction | + | + | + | + | – | + | + |
| Widow's peak/low anterior hairline | + | + | + | – | – | + | – |
| Hypertelorism | + | – | + | – | – | – | + |
| Synophrys | – | + | + | – | + | + | – |
| Sparse medial eyebrows | – | + | – | – | – | – | – |
| Deeply set eyes | + | + | + | + | – | – | – |
| Epicanthus | + | + | – | – | + | – | + |
| Long eyelashes | + | – | + | – | – | + | + |

| | | | | | | | |
|---|---|---|---|---|---|---|---|
| Flat/low nasal bridge | – | – | – | + | – | + | + |
| Short nose | + | – | + | - | + | + | + |
| Beaked nose | – | – | + | + | – | – | – |
| Anteverted nares/Hypoplastic | – | + | + | + | – | + | – |
| Protruding columella | + | + | – | + | + | – | – |
| Short philtrum | + | + | + | + | – | + | + |
| Microstomia | – | – | – | – | + | + | – |
| Carp-like mouth | – | + | – | – | – | + | – |
| Thin lips /thin upper lip | **+ (thin lower lip)** | + | – | + | – | + | + |
| Widely spaced teeth | + | + | + | – | **+ (primary teeth)** | + | + |
| Prominent superior incisors | + | – | – | + | – | – | – |
| Dysplastic/orverfolded helices/ears | + | – | – | + | + | + | + |
| Crumpled ears | + | – | – | + | + | – | – |
| Short nails | + | + | – | – | + | + | – |
| Tapered fingers | + | + | – | + | – | – | + |
| | + | + | – | – | – | | – |
| Pes planovalgus/neurogenic | + | + | – | – | – | – | – |
| 1st toe longer than others/ Sandal gap | + | + | + | + | | – | + |
| Unilateral postaxial polydactyly of | – | – | – | + | – | – | – |
| **Cardiologic findings:** | stenosis of arteria pulmonalis | membranous septal defect, patent foramen ovale, coarctation of the aorta (surgery required) | cardiac anomalies (intra ventricular and auricular communication), surgery required | atrial septal defect | ventricular septal defect (surgically repaired) | coarctation of aortic isthmus; dysplastic pulmonary valve | patent foramen ovale in infancy, resolved |
| **Hearing impairment:** | no | no | no | moderate deafness (35 dB) | no | no | no |
| **Visual impairment:** | hypermetropia, glasses | no | no | no | ? | incipient myopia at last review | alternating extropia; hyperopia with astigmatism, bilateral; maculopathy, bilateral; premature macula |
| **Skeletal anomalies** | no | no | no | no | ? | no | cutanesous syndactyl 2nd/3rd toes |
| **Gastrointestinal anomalies:** | pylorostenosis, surgery | no | no | no | chronic constipation | antenatally detected large stomach and duodenum. Recurrent abdominal distension. Hirschbrung disease excluded on biopsy | GERD, constipation, umbilical hernia |
| **Other:** | growth and sexual maturation disorder | kidney cysts in the renal medulla, undescended testicles, inguinal hernia | kidney anomalies (cyst on the left one) | no | stenosis of one lacrymal duct, pyelonephritis at 20 m and posterior urethral valves (surgically repaired) | livedo reticularis; follicular hyperkeratosis of upper limbs; delayed closure of anterior fontanelle; delayed eruption of primary dentition; obstructive sleep apnea on BiPAP therapy | cryptorchidism at birth, resolved without surgery; Seizure-like activity vs sensory seeking behaviors (had twitching, arches head backwards and forewards, extreme belly laughts without provocation - all followed by period of prolonged sleep); unremarkable EEGs |
| **Genetic analyses:** | | | | | | | |
| Microarray | 15q26.1 microduplication, inherited from the mother | ND | ND | normal | 400 kb | normal SNP microarray | normal male |
| Karyotype and results of other genetic testing | 46, XX, FISH22 negative, William-Beuren syndrome, neuromuscular disorders, SMA, mtDNA deletions, Prader-Willi syndrome, Rett syndrome - all negative | Prader-Willi syndrome, aCGH, clinical exome sequencing - all negative | ND | COH1 gene negative | ND | karyotype 46, XX | negative trio exome; negative mitochondrial genome seq & del/dup |
| Biochemical testing | ND | ND | ND | ND | ND | Extensive metabolic screens negative. | abnormall LSD at birth, normal at our evaluation |
| Family History | negative | negative | negative | negative | | negative | maternal uncle deceased; brain teratoma |
| Other notes | NA | polyhydramnion and intrauterine growth restriction, maternal infection by parvovirus during the last trimester of pregnancy | NA | NA | IVF | Antenatal polyhydramnios, large stomach and duodenum. | IUGR; born at 37 weeks gestation, admitted to NICU for respiratory distress for 25 days |
